# Evolution of chromatin accessibility associated with traits of cichlid phenotypic diversity

**DOI:** 10.1101/2025.10.09.681187

**Authors:** Tarang K. Mehta, Angela L. Man, Graham Etherington, Alan M. Smith, Adrian Indermaur, Walter Salzburger, Domino Joyce, Federica Di-Palma, Wilfried Haerty

## Abstract

The radiations of cichlid fishes in East African Lakes Victoria, Malawi, and Tanganyika showcase a remarkable example of rapid adaptive speciation, with over 2000 species evolving diverse morphological and ecological adaptations within the last few million years. Understanding the molecular basis of this phenotypic diversity remains a key challenge. Building on prior evidence of gene regulatory network (GRN) rewiring underpinning adaptive traits, we profiled chromatin accessibility (ATAC-seq) and matched transcriptomes across forebrain, retina, liver, and testis tissues in five representative cichlid species using optimised protocols. We show extensive divergence in chromatin accessibility corresponding to phylogenetic lineages and tissue identity, with many regulatory regions exhibiting accelerated nucleotide evolution. Transcription factor binding site (TFBS) variation correlates with both chromatin accessibility and differential gene expression, particularly in genes linked to sensory systems. Building on this, we reconstructed tissue- and species-specific GRNs and show that motif-supported network inference reveals pervasive but tissue-dependent rewiring, with the strongest candidate edges concentrated among accessible, highly expressed genes linked to adaptive traits. By integrating TF footprinting with regulatory motif turnover analyses, we demonstrate that dynamic nucleotide changes are associated with GRN rewiring, concordant with ecological niche and lineage-specific adaptations. Our findings highlight that regulatory variation at conserved and novel TFBSs associates with genes linked to phenotypic innovation across radiating and non-radiating East African cichlids. This study provides foundational epigenomic evidence linking GRN divergence to key mechanisms facilitating rapid adaptive diversification in this iconic vertebrate radiation.

## Introduction

Seminal studies analysing vertebrate protein evolution concluded that evolutionary changes in ‘regulatory systems’ are a key driver of morphological diversity ^1–4^. These systems comprise *cis*-regulatory elements (CREs) such as promoters, enhancers, and silencers, which harbour transcription factor binding sites (TFBSs) that spatiotemporally control gene expression during development ^5^. While several locus-specific studies have demonstrated the role of CRE evolution in morphological change ^6–10^, novel sequencing methods now provide genome-wide views of chromatin accessibility and TF-binding ^11^. The assay for transposase-accessible chromatin using sequencing (ATAC-seq) is particularly powerful due to its speed and simplicity, enabling chromatin accessibility profiling from ultra-low input ^12^ or even single cells ^13^. Beyond insights into mammalian CRE evolution ^14^, ATAC-seq allows direct assessment of *cis*-regulatory divergence in adaptive radiations ^15–18^.

Ray-finned fishes represent the most species-rich vertebrate radiation, and among them, the East African cichlids are a striking case of explosive speciation ^19^. Within the past few million years ^20,21^, one or a few ancestral lineages gave rise to more than 2000 species in the African Great Lakes (Victoria, Malawi, Tanganyika). These lineages occupy diverse ecological niches, exhibiting extensive phenotypic diversity in colouration, skeletal morphology, trophic structures, and behaviour ^22^. Previous studies have demonstrated that the rapid evolution of regulatory elements ^23^ and regulatory changes driving gene regulatory network (GRN) rewiring ^24^ have contributed to cichlid diversification. Other work has shown evolutionary changes in noncoding regulatory regions in the Lake Victoria radiation ^23,25^ and linked gene expression divergence to speciation and trait diversification in Lake Tanganyika ^26^.

Although still relatively unexplored, epigenomic studies in cichlids have begun to illuminate regulatory evolution. Initial work mapped active promoters in *O. niloticus* fin development and regeneration using H3K4me3 ChIP-seq ^27^, and our recent work integrated ATAC-seq and RNA-seq analyses to identify regulatory loci underlying salinity and osmotic stress acclimation in tilapia ^28^. Other studies have connected *cis*-regulatory divergence with dietary ecology in Lake Malawi cichlids ^29^, speciation processes ^30^, and sexual dimorphism in tilapia ^31^, as well as jaw shape variation to regulatory divergence in Malawi species ^18^. However, no study has yet compared *cis*-regulatory sequence evolution with tissue- and lineage-specific chromatin accessibility and transcription across multiple African cichlid lineages. Here, we present the first genome-wide, multi-tissue comparative epigenomic analysis across multiple East African cichlid species, linking sequence conservation, regulatory accessibility, and transcriptional divergence to identify how regulatory evolution could shape adaptive phenotypic diversity.

## Results

### Cichlid chromatin accessibility is enriched in noncoding regulatory regions

To characterise variation in chromatin accessibility associated with phenotypic diversity in cichlid fishes, we performed ATAC-seq on four tissues that link to traits under natural/sexual selection—forebrain, retina, liver, and testis— from five representative East African cichlids used in comparative studies ^32^ and spanning the major haplochromine radiation and the more distantly related lamprologine lineage: *Metriaclima zebra* (Lake Malawi)*, Pundamilia nyererei* (Lake Victoria)*, Astatotilapia burtoni* (Lake Tanganyika associated, but also inhabiting multiple rivers) from the haplochromine radiation, *Neolamprologus brichardi* from the lamprologine lineage of Lake Tanganyika, and *Oreochromis niloticus* as a riverine and lacustrine freshwater reference species (Fig. 1a and Supplementary Table S1; see ‘*Materials and Methods*’). This dataset enabled genome-wide annotation and comparison of conserved and species specifc open chromatin regions across the phylogeny(Fig. 1d). Across all species and tissues—using consensus peaks across replicates (see ‘*Materials and Methods*’)—on average 78-99% of open chromatin peaks mapped within 5 kb of a transcription start site (TSS), and 10-20% of these peaks were located in the core promoter region (±100 bp ^33^ of the TSS) (Fig. 1b-c, Supplementary Fig. S1-S2, Supplementary Table S2-S3; see ‘*Supplementary information’*). On average, genome-wide annotation revealed that across all five species, peaks overlapped intronic regions most frequently (40%), followed by exons (23.5%) and intergenic regions (17.4%), with smaller fractions assigned to 5′ UTRs (6.5%), promoter regions within 5 kb upstream of the TSS (10%), 3′ UTRs (2.5%) and conserved noncoding elements (0.1%) (Supplementary Fig. S2, Supplementary Table S3). This overall distribution is broadly consistent with the annotation profiled reported for human ATAC-seq datasets of over 300 samples ^34^.

**Figure 1.**
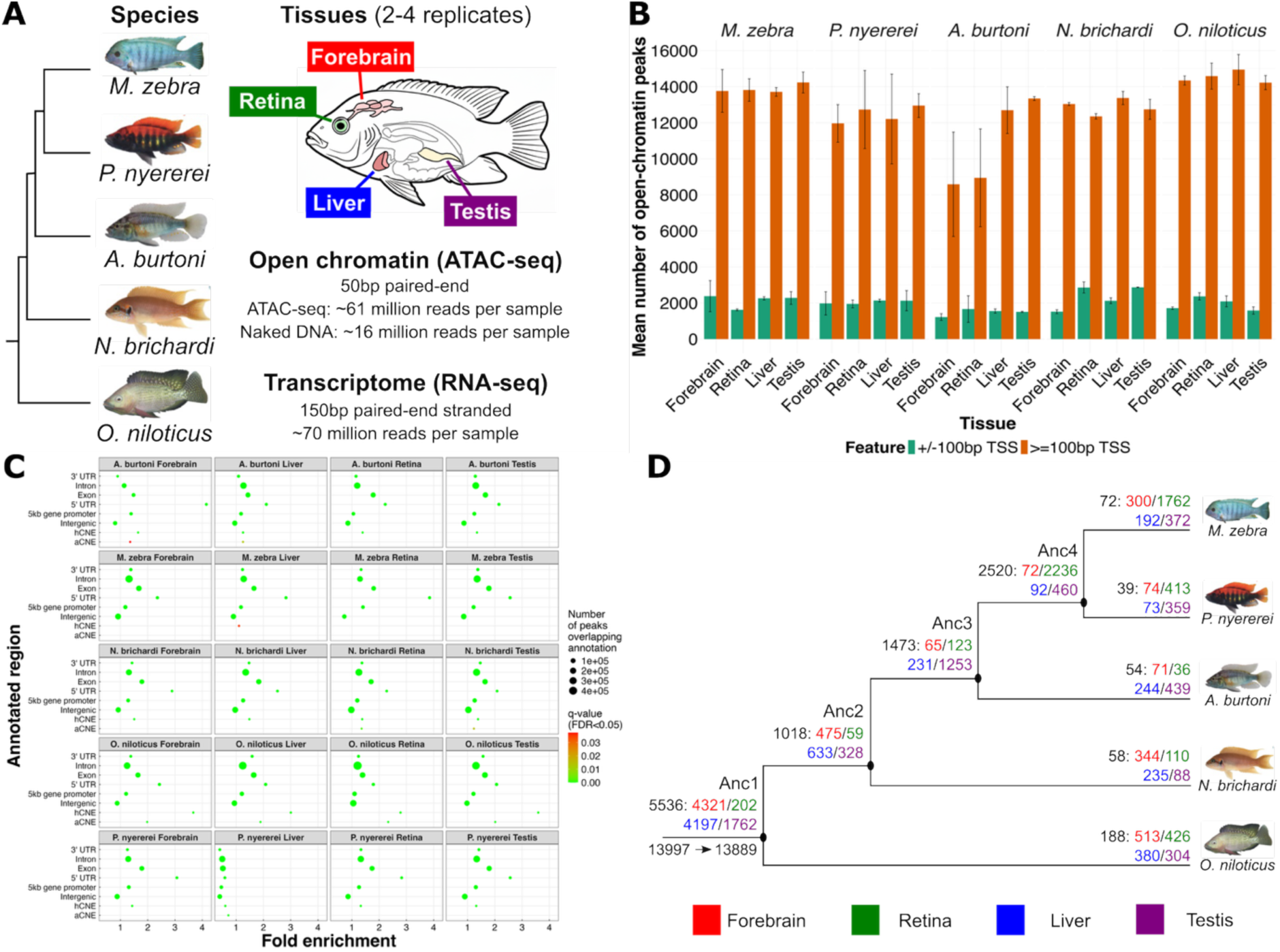
The open chromatin landscape of five cichlid species. **(A)** Summary of the species, tissues, and sequencing design of this study. See ‘*Materials and Methods*’ and Supplementary Table S1. **(B)** Mean number of open chromatin peaks across the five species and four tissues with standard deviation (error bars), separated by peaks that are located either +/- 100 bp from a Transcription Start Site (TSS) or >=100 bp from a TSS. **(C)** Fold enrichment of tissue-specific open chromatin peaks overlapping coding and noncoding regions in the five cichlid genomes. Circles show enriched annotated region (y-axis) of significance (FDR <0.05, to right) and fold enrichment (x-axis) values of all annotated peaks. Number of peaks overlapping each annotation shown by size of each circle. **(D)** Number of 1-to-1 orthologous gene with associated open chromatin peaks (overlapping up to 5kb gene promoter) shown along the phylogeny, indicating in black the count of orthologous genes with positionally conserved peaks at ancestral nodes or those specific to each species, and in red, green, blue, or purple the count present in forebrain, retina, liver, or testis, respectively.

### Core regulatory elements reveal conserved and clade-specific chromatin landscapes

To define the core gene regulatory architecture shaping cichlid diversity, we mapped promoter open chromatin peaks of species-tissue replicates to 13,889 1-to-1 orthologous genes across forebrain, retina, liver, and testis of five species, revealing an enrichment of broadly shared processes such as epithelium development together with tissue-specific functions including synapse organisation (forebrain), axon extension (retina), myotube differentiation (liver), and dephosphorylation (testis) (Fig. 1d; Supplementary Fig. S3; *see ‘Materials and Methods’*). We first examined positionally “conserved” promoter peaks, defined as open chromatin peaks with overlapping summits across species (see *‘Materials and Methods’*). In total, we identified 5,536 orthologous genes, representing 40% of all 13,889 1-to-1 orthologous genes, with such positionally conserved promoter peaks shared across all five species within the same tissue (202**–**4,321 genes per tissue), indicating positionally maintained regulatory elements along the phylogeny (“conserved” at ancestral node 1, *Anc1*, Fig. 1d; see *‘Materials and Methods’*). These genes are enriched for fundamental biological processes such as ‘cell migration’ (Supplementary Fig. S4; Supplementary Table S4), and include key developmental and housekeeping regulators, for example *irx2*, a core vertebrate neural differentiation gene ^35^. Within *P. nyererei*, *M. zebra,* and *A. burtoni* (haplochromine species), we identified 1,473 non-redundant genes (65–1,253 orthologous genes per tissue) with positionally conserved promoter peaks across tissues (“conserved” in haplochromine ancestor - ancestral node 3, *Anc3*, Fig. 1d). Gene ontology analysis linked these positionally conserved peaks to core vertebrate functions such as sensory system development (Supplementary Fig. S4; Supplementary Table S4) and to promoter activity of fundamental genes, for example *pax6*, a master regulator of eye development in all seeing animals ^36^. Together, these evolutionarily maintained core regulatory elements could underpin shared biological functions and key aspects of cichlid development, while clade-specific conservation identifies ancestral regulatory programs that could shape adaptations in species of tested lineages.

### Conserved and species-specific promoter accessibility reveals tissue-specific regulatory variation

We next explored unique patterns of chromatin accessibility that distinguish species and tissues by classifying non-overlapping promoter peaks as ‘unique’, that is, peaks whose summits were not encompassed within a shared interval between species (see *‘Materials and Methods’*). Across all five species and four tissues, species-specific promoter peaks were associated with a total of only 39–188 non-redundant orthologous genes (Fig. 1d). However, these ‘unique’ peaks likely contribute to regulatory diversity, especially in the retina, where the five species together showed a total of 2,747 non-redundant orthologous genes with unique gene promoter peaks compared with 1,124–1,562 in the other tissues (Fig. 1d). Unique peaks also showed high variability at both the ancestral and species level (ancestral coefficient of variation, CV = 162%, species-specific CV = 122%; Fig. 2a; see ‘*Supplementary information’*). Notably, the retina exhibits the fewest positionally ‘conserved’ promoter peaks in only 202 orthologous genes (*Anc1* node, Fig. 1d), representing only 1.8–2.9% of the 6,981–11,382 1-to-1 orthologous genes that are expressed in retina tissue across the five species. This high variation in chromatin accessibility supports previous findings of dynamic, lineage- and clade-specific rewiring of visual system regulatory networks in the same species ^24,37^.

**Figure 2.**
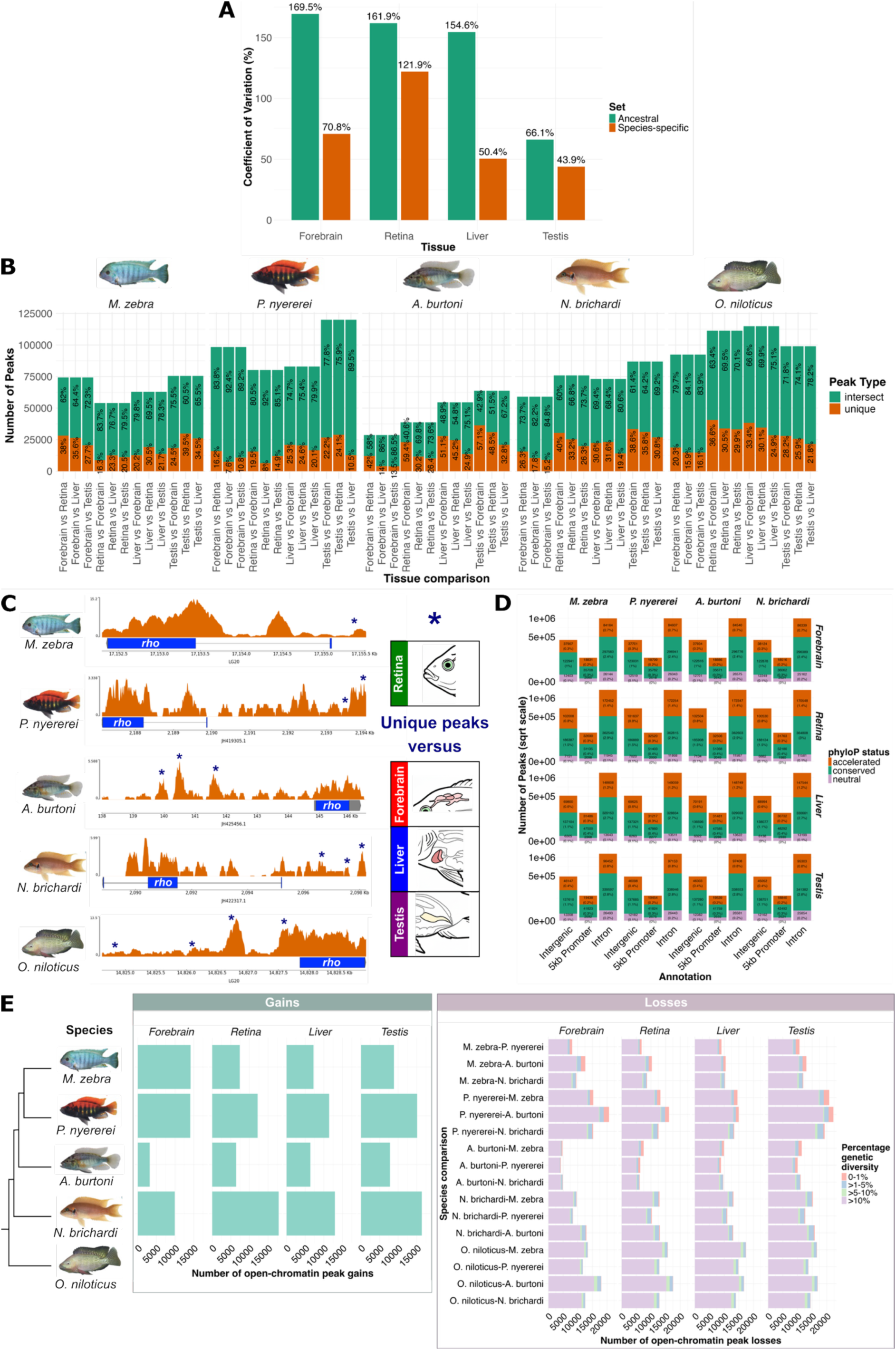
Evolution of gene promoter peaks across species and tissues. **(A)** Coefficient of Variation (CV) in the number of promoter peaks in orthologous genes per tissue, comparing the ancestral node of all species (Anc1 in Fig. 1d; green bars) with species-specific sets (orange bars); CV values shown above each bar. **(B)** Number and percentage of shared and unique gene promoter peaks between all pairwise tissue comparisons within species. Peaks were aggregated using the highest -log10 *q-value* for fully overlapping replicates, with each reference tissue peak counted once if overlapping, resulting in asymmetry for reverse comparisons, where a single reference peak overlapping *n* comparison peaks is counted *n* times (see Supplementary Table S5). **(C)** Retina-specific peaks (navy stars) versus other three tissues in the *rho* gene promoter region of each species. **(D)** Stacked plot (square root scale) of phyloP-classified conserved, neutral, and accelerated peaks found in intergenic, promoter, or intronic regions across four species and tissues, using *O. niloticus* as the reference. True peak counts and percentages of total peaks per annotation are indicated within bar stacks; low-count annotations (UTRs, exons, CNEs) omitted for clarity and presented in Supplementary Table S6. **(E)** Number of open chromatin peak gains and losses in 1-to-1 orthologous gene promoters, with species-specific gains and pairwise losses determined by lack of overlapping peaks in promoter alignments. Pairwise losses are binned by percentage genetic diversity (0–1%, >1–5%, >5–10%, >10%) between the reference and aligned region in the other species.

To further dissect regulatory specificity, we performed pairwise comparisons of chromatin peaks between tissues within each species, defining promoter peaks as ‘conserved’ only when both tissues’ peak summits fell within the shared interval and as ‘unique’ otherwise. Promoter peaks showed high conservation across all 60 tissue-tissue comparisons (mean 72.7% shared ± 11.3 SD, range 41-92%; Supplementary Table S5). Remarkably, retina tissue displayed a substantial proportion of unique peaks across all comparisons (mean 29.8% ± 10.9 SD, n=30; range 8-59%), the highest among all tissues (forebrain 27.3%, liver 26%, testis 26.2%), albeit with no significant difference across them (ANOVA *p*=0.55; Supplementary Table S5). Nonetheless, this overall prevalence of retina-specific chromatin peaks mirrors findings in human retina, where sensory tissues exhibit distinct and highly dynamic accessible regions ^38,39^, and the identification of ∼41% retinal specific peaks in retina and retinal pigmented epithelium comparisons ^40^.

Conserved peaks across pairwise tissue comparisons identify genes implicated in shared tissue functions—such as axonal guidance, e.g., *nrp1a*^41^ in forebrain/retina of all species—while unique peaks are often associated with tissue-specific genes, e.g., the dim-light vision gene *rhodopsin* (*rho*) in the retina of all species (Fig. 2c). These findings indicate that, while a core of conserved regulatory elements is important for shared functions, tissue-specific peaks could be important for evolutionary divergence in the five species—particularly in traits under natural and sexual selection, such as sensory systems.

### Accelerated divergence of active gene promoter sites contribute to cichlid regulatory diversity

To investigate the scale of gene regulatory evolution along the cichlid phylogeny, we calculated rates of nucleotide substitutions ^42^ at gene promoter chromatin peaks, using the phylogenetic outgroup species, *O. niloticus*, as an unbiased reference for comparisons with the four other species (see ‘*Materials and Methods*’). Notably, compared to other noncoding (intergenic and intronic) regions, gene promoters showed the highest proportion (30–39%) of open chromatin peaks undergoing accelerated evolution (Fig. 2d), exceeding rates observed under a neutral model—underscoring the rapid divergence of *cis*-regulatory elements at gene promoter sites (Supplementary Fig. S5-S6; Supplementary Table S6-S7; see ‘*Supplementary information’*). Among 5,536 orthologous genes (202–4,321 genes across tissues) with positionally ‘overlapping’ promoter peaks across all species (Anc1, Fig. 1d), over half (3,061 orthologous genes, 55%) demonstrated accelerated evolution (see ‘*Materials and Methods*’), with significant enrichment for genes tied to key developmental processes such as retinal development and cell junction assembly (Supplementary Fig. S7). This prevalent pattern reveals that even deeply conserved gene promoter peaks—often retained due to the functional importance of associated genes—can undergo sequence divergence in at least one lineage, suggesting regulatory sites may remain spatially conserved while their underlying sequence evolves to facilitate clade-specific functional adaptations.

To further quantify regulatory evolution, we measured gain and loss of promoter peaks across each lineage in relation to the corresponding genetic diversity (Fig. 2e, Supplementary Table S8 and S9). With no previous knowledge of expected divergence at open chromatin sites, we use overlapping *O. niloticus* peak summits as a reference in all comparisons (Supplementary Table S8, see *Materials and Methods*). Pairwise comparisons based on overlapping promoter peaks of 1,956-4,983 1-to-1 orthologous genes revealed that 37–66% (mean of 52%) of peaks across all species and tissues retained strong sequence conservation (less than 10% divergence, Supplementary Table S8), consistent with 85–89% pairwise nucleotide identity across cichlid genomes ^24^ (see *Materials and Methods*). However, species- and tissue-specific differences in gene promoter peak gains were also evident, and unbiased by genome completeness or annotation quality (see ‘*Supplementary information’*); for example, *N. brichardi* exhibited the most gains in retina (52% of all 13,889 1-to-1 orthologous genes), while *M. zebra* showed the highest gains in forebrain tissue (45% of 1-to-1 orthologous genes)—often enriched (Benjamini-Hochberg [BH] ^43^ adjusted *p-value* <0.05) for relevant tissue-specific functions, such as neurogenesis (in *M. zebra)* or monatomic ion transport (in *A. burtoni;* Supplementary Fig. S8). Peak loss was similarly variable across species, with the majority (76–91%) of losses occurring in regions with elevated sequence divergence (greater than 10% difference; Fig. 2e, Supplementary Table S8), suggesting that sequence evolution frequently underpins regulatory site turnover, including the potential gain or loss of transcription factor binding sites.

### Divergence of transcription factor activity underpins adaptive regulatory innovations

Transcription factors are central drivers of gene regulatory innovations, shaping networks associated with cichlid phenotypic diversity ^24,28,37^. To systematically profile TF-mediated regulation, we took advantage of our experimental ATAC-Seq data to map between 54,000 and 226,000 TF footprints within gene promoters (representing 8–13% of each genome) across four tissues and five species, using 1,442 unique TF motifs spanning cichlid ^37^ and vertebrate ^44,45^ lineages. Profiling of footprints with 1,442 vertebrate motifs yielded between 530,000 and 2,200,000 non-redundant TFBSs (Supplementary Table S10a-b, see *Materials and Methods*), with a mean of 1.4 million non-redundant promoter TFBSs per sample; this is remarkably consistent with the 3.6 million footprints mapped to the ∼3× larger human genome^46^.

Consistent with previous work ^24,28,37^, we found that TFBSs are significantly enriched (BH ^43^ adjusted *p-value* <0.05) in noncoding regulatory regions, especially gene promoters and conserved noncoding elements (CNEs; Fig. 3a and Supplementary Table S11). Notably, TFBSs are over four-fold more abundant in highly conserved CNEs (hCNEs with sequence identity ≥90% over ≥30bp, see *Materials and Methods*), identifying TF binding sites maintained by selective constraint. In some tissues and species, e.g., *O. niloticus* forebrain and *N. brichardi* retina (Fig. 3a and Supplementary Table S11), TFBSs are more than two-fold enriched in accelerated CNEs (aCNEs with sequence identity <90% over ≥30bp and significantly deviating from a neutral model, see *Materials and Methods*)—highlighting a set of novel, potentially lineage-specific regulatory sites, such as the previously characterised NR2C2-*sws1* interaction in *N. brichardi* ^37^.

**Figure 3.**
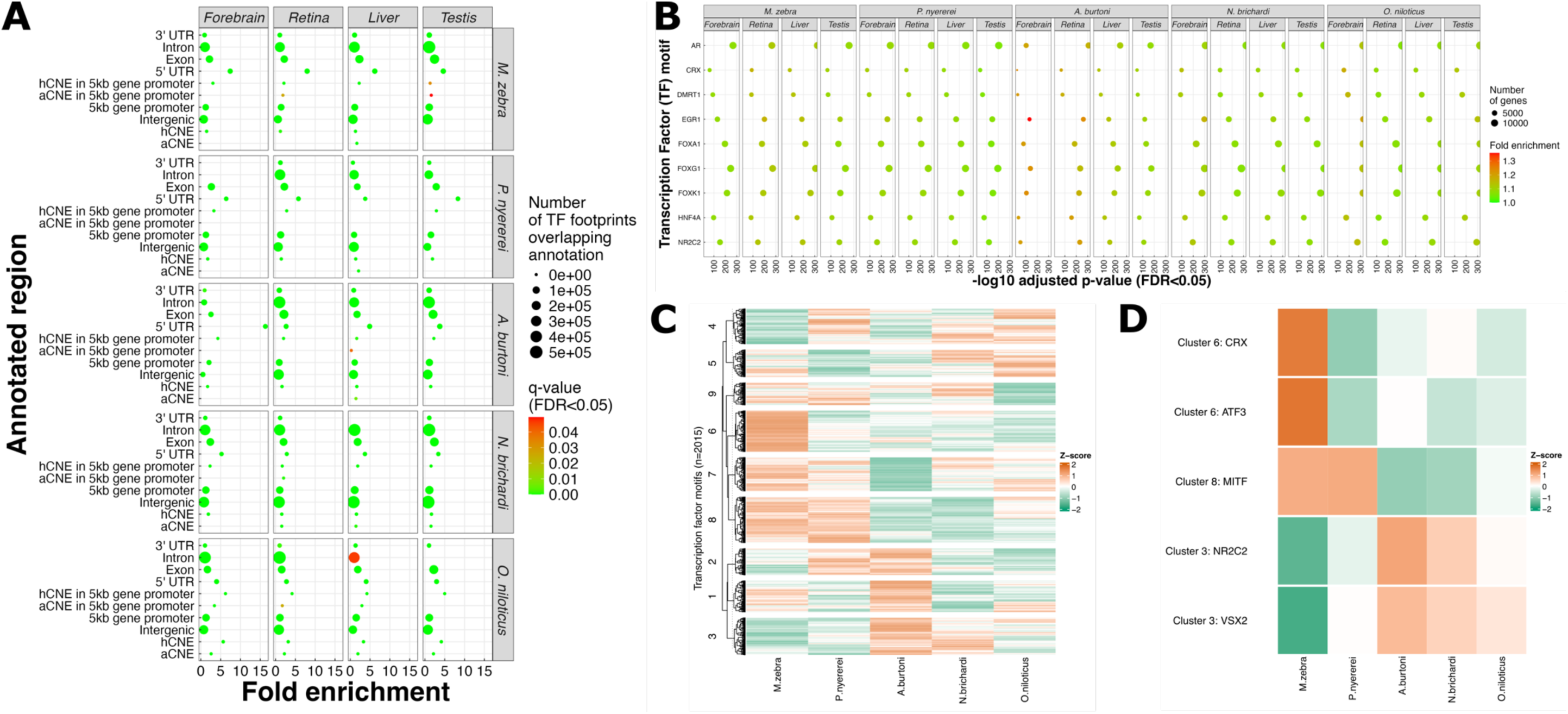
Transcription factor motif profiling of chromatin accessibility evolution in five cichlids. **(A)** Fold enrichment of TF motifs overlapping coding and noncoding regions across species and tissues. Circles show enriched annotated region (y-axis) of significance (FDR <0.05, to right) and fold enrichment (x-axis) values of all annotated peaks. Number of TF motifs overlapping each annotation shown by size of each circle. **(B)** Enrichment of nine functionally associated TF motifs in gene promoter peaks across species and tissues. Circles show TF motif (y-axis) with fold enrichment (heatmap to right) and significance of enrichment as -*log_10_* FDR <0.05 (x-axis) across each species and tissue. Number of enriched genes for each TF motif shown by size of each circle. **(C)** Clustering of TF activities in gene promoter regions of retina tissue across the five species. Each row represents a single, clustered TF, and each column a species. Mean TF bit-score of motif match across gene promoters (as a measure of activity) from HINT-ATAC ^113^ are *Z*-score transformed across rows. Green colour indicates low activity and orange colour indicates high activity. **(D)** TF activities and cluster assignment of five eye-related TFs in gene promoter regions of retina tissue across the five species. Each row is an eye-related TF, and each column a species. Mean TF bit-score of motif match across gene promoters (as a measure of activity) from HINT-ATAC ^113^ are *Z*-score transformed for each TF. Green colour indicates low activity and orange colour indicates high activity.

To explore TF functional relevance, we identified 824 (57%) of the 1,442 input vertebrate TF motifs significantly enriched (BH ^43^ adjusted *p-value* <0.05) in active promoter regions across tissues (Supplementary Table S12), many with known roles in vertebrates, as well as cichlid adaptive traits. Topmost enriched examples across species include: *foxg1* and *egr1* in the forebrain (implicated in neural diversity ^47^ and behaviour ^48^), *crx* and *nr2c2* in the retina (opsin gene regulation ^37,49^), *hnf4a* and *foxa1* in the liver (diet ^29^ and hepatogenesis ^50^), and *ar* and *dmrt1* in the testis (sex differentiation ^51^ and reproductive traits ^52^; Fig. 3b, Supplementary Table S12).

Clustering tissue-specific TF activity within each species (see *Materials and Methods*) revealed distinct co-active groups of TFs (Fig. 3c, Supplementary Fig. S9-S11), reflecting both conserved and divergent patterns of transcriptional regulation (see ‘*Supplementary information’*). In retina tissue (Fig. 3c), clear differences emerged among species: for example, *M. zebra* exhibited strong co-activity (Z-score of 1.6 versus −1 to 0.05 in other species, Fig. 3d) of *crx* and *atf3*, which are key regulators of retinal development ^24,53^ and opsin expression ^54^ (Supplementary Fig. S12), while *P. nyererei* and *M. zebra* shared high *mitf* activity (Z-score of 1 versus - 0.1 to −1 in other species, Fig. 3d), related to retinal pigment epithelium formation^55^ (Supplementary Fig. S12). Conversely, *A. burtoni* and *N. brichardi* showed higher activity for *nr2c2* and *vsx2* (Z-score of 1.1 and 0.6 versus 0 to −1.5 in the other species; Fig. 3d; Supplementary Fig. S12), implicating divergent regulatory programs for thse genes associated with cichlid visual system adaptation ^37^.

Taken together, these results demonstrate that divergence in TF binding activity—through both conserved and lineage-specific regulatory motifs—could enable regulatory plasticity of genes underlying cichlid adaptive traits, reinforcing associations between regulatory innovation and cichlid phenotypic diversification.

### Dynamic regulatory site divergence is associated with regulatory rewiring in cichlids

To directly link the observed divergence of TF binding activity with genetic changes in regulatory elements, we examined nucleotide variation at active TFBSs in orthologous gene promoters. Using the phylogenetic outgroup species, *O. niloticus*, as the reference (see *Materials and Methods*), we found that the vast majority (76–78%) of TF motifs in the other four cichlid species only partially positionally overlap (<100%) with the reference, and nearly all motifs (99.1–99.9%) differed in at least one position or context (Fig. 4a, Supplementary Table S13). This points to pervasive divergence of TF binding landscapes, even at orthologous promoter regions across species.

**Figure 4.**
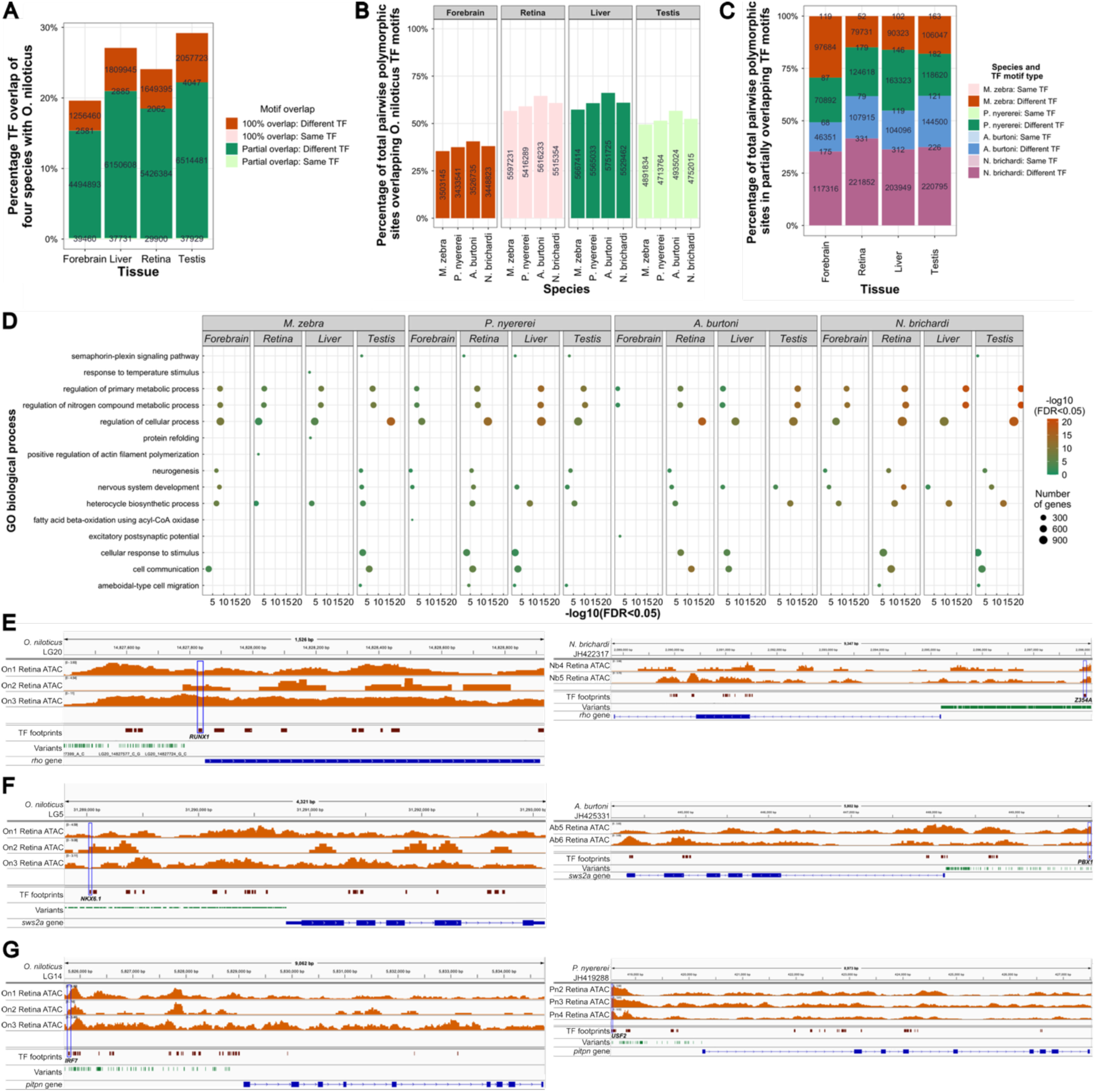
Nucleotide divergence of actively bound TFs across five cichlids. **(A)** Percentage TF motif overlap of same or different motifs of four species versus *O. niloticus* reference across four tissues. Number of TF motifs within each category are shown within each bar. **(B)** Percentage of total pairwise polymorphic sites overlapping *O. niloticus* TF motifs in the four species and tissues. Number of pairwise polymorphic sites in each species tissue versus *O. niloticus* shown within each bar. **(C)** Percentage of total pairwise polymorphic sites in partially overlapping same or different *O. niloticus* TF motifs in the four species and tissues. Number of pairwise polymorphic sites overlapping same or different TF motifs in each species tissue versus *O. niloticus* shown within each bar. **(D)** Gene ontology (GO) biological process enrichment of accessible gene promoter regions containing pairwise polymorphic variants versus *O. niloticus* across the four species and tissues. GO terms are a subset of enrichment -log10 (FDR<0.05) of >1.3. Circles show enriched term (y-axis) with significance of enrichment as -*log_10_* FDR <0.05 (x-axis and heatmap to right) across each species and tissue. Number of enriched genes for each GO term shown by size of each circle. Species chromosome tracks of **(E)** *O. niloticus* (left) and *N. brichardi* (right) *rho* gene; **(F)** *O. niloticus* (left) and *A. burtoni* (right) *sws2a* gene; and **(G)** *O. niloticus* (left) and *P. nyererei* (right) *pitpn* gene. Coding sequence shown as blue annotations including upstream gene promoter regions, accessible peaks across retina tissue biological replicates (orange peaks), TF footprints (maroon mark), and pairwise polymorphic variants (green marks). Candidate actively bound TF footprints with pairwise variation are labelled highlighted with blue box.

To quantify the genetic basis of these regulatory differences, we identified 9–10 million pairwise polymorphic sites within aligned promoter regions of 10,699 out of the 13,889 1-to-1 orthologous genes across the five species (Supplementary Table S14a). On average, 53% of promoter variants (range 35–66%) fell within and affected predicted *O. niloticus* TF binding motifs in promoters of 6,842–8,200 orthologous genes per tissue (Fig. 4b, Supplementary Table S14b). These motifs carried only 43% as many variants per affected gene promoter versus those promoters without variants overlapping motifs (paired t-test across 16 species-tissues: t= −16.8, *p* = 3.9 x 10^-11^), indicating purifying selection. We defined a ‘principal set’ of 1–5% of all promoter variants (1,385-1,439 of the 1,442 non-redundant input vertebrate TF motifs in 749–2,747 genes) as those disrupting partially (<100%) overlapping motifs between species and *O. niloticus,* where 99.9% of motifs diverged (Supplementary Table S14c). Notably, 6–13% of principal set variants (175–1,361 motifs in up to 618 orthologous genes) occurred in accelerated peaks (Supplementary Table S14c). These regions likely represent hotspots of regulatory innovation driving species-specific regulatory changes.

Functional annotation of ‘principal set’ genes affected by TFBS divergence revealed significant enrichment (BH ^43^ adjusted *p-value* <0.05) for biological processes central to vertebrate traits, such as neurogenesis in the forebrain and cellular process regulation in the retina (Fig. 4d). This includes known cichlid adaptive trait genes like, for example, in the *bmpr1* and *rho* promoters of *P. nyererei* and *N. brichardi,* a total of four nucleotide changes (versus *O. niloticus*) alter motifs from CTCFL/RUNX1 to ZN768/Z354A, implicating regulatory evolution in their functions of jaw morphology ^56^ and dim-light vision ^57,58^ (Fig. 4e). Similarly, in *M. zebra* and *A. burtoni*, five and six respective variants in the *cntn4* and *sws2a* promoters result in turnover from ZNF263/NKX6.1 to MEIS1/PBX1 motifs, potentially affecting how these two genes that are linked to either morphogenesis ^23^ and visual system ^59^ divergence ^24,37^ are regulated (Fig. 4f). Both discrete (single-nucleotide change) and complex motif rewiring (multiple nucleotide changes)—such as the replacement of IRF7 with USF2 motifs in the promoter of the retinal function related gene, *pitpn* ^60^ (Fig. 4g) and STAT1 with a TEAD4 motif in the promoter of the liver development gene, *foxa2* ^50^—were also observed in additional tissue- and lineage-specific contexts (see ‘*Supplementary information’*).

In summary, dynamic nucleotide divergence at active regulatory binding sites could underpin the rewiring of gene regulatory networks in cichlids, providing a rapid molecular mechanism for the evolution of adaptive traits across lineages.

### Chromatin accessibility divergence correlates with transcriptional regulation of genes linked to cichlid adaptive traits

While transcriptional variation can underlie cichlid diversification ^26^, the precise role of epigenetic regulation in driving tissue-specific transcriptomic divergence across East African lineages remains understudied. To address this, we integrated chromatin accessibility and RNA-sequencing data from matched tissues across the five cichlid species (see ‘*Materials and Methods*’, Fig. 1a, Supplementary Table S1). Among the 23,044–23,568 protein-coding genes per genome, we quantified expression (as transcripts per million, TPM) for 13,121–21,163 genes (56–75% of all coding genes per species). Across species and tissues, 20,652–226,889 accessible chromatin peaks (47–94% of identified peaks) were associated with the expression of 6,657–18,787 genes (29–70% of protein-coding genes per genome; Supplementary Table S15). These differences reflect tissue- and species-specific regulatory complexity e.g., testis and forebrain have the highest numbers of associated peaks (129,254–166,802) and genes (13,374–15,991) on average, whereas liver (90,452 peaks and 10,331 genes) and retina (83,902 peaks and 12,122 genes) show fewer (Supplementary Table S15), rather than absence of gene expression.

Assessing regulatory relationships, we confirmed a weak but significant positive correlation between promoter chromatin accessibility (within 5 kb of the TSS) and gene expression level in 18 out of 20 species-tissue datasets (Spearman’s R=0.014 to 0.21, *p*<0.05; Supplementary Fig. S13–S17). Exceptions included *A. burtoni* (R=-0.02, *p*<2e-7) and *M. zebra* (R=-0.003, *p*=0.53) liver, where the correlation was weak or absent (Supplementary Fig. S13 and S15). Overall, weak positive correlation is consistent with prior findings in other *O. niloticus* tissues e.g., gill ^28^ and reflect the multifaceted roles of accessible regions consistent with gene activation, repression, or poised states ^61^.

To dissect regulatory modes, we categorised genes based on peak accessibility—high (HA) or medium/low (MA)—and by expression level—high (HE) or medium/low (ME; see ‘*Materials and Methods*’). Of all identified non-redundant peaks, 5,222–68,245 (12–28%) mapped to 1,864–5,404 genes (8–20% of coding genes per genome; Supplementary Fig. S18–S22, Supplementary Table S16). In retina tissue, where adaptive regulatory divergence is marked (Fig. 4), most peak-gene relationships (37–41%) fell into the HA-HE category (1,408–4,842 peaks and 902–2,020 genes per species; Supplementary Fig. S18–S22, Supplementary Table S16). The remainder were distributed as HA-ME (23–24%), MA-HE (20–24%), and MA-ME (15–16%; Fig. 5a, Supplementary Fig. S18–S22, Supplementary Table S16).

**Figure 5.**
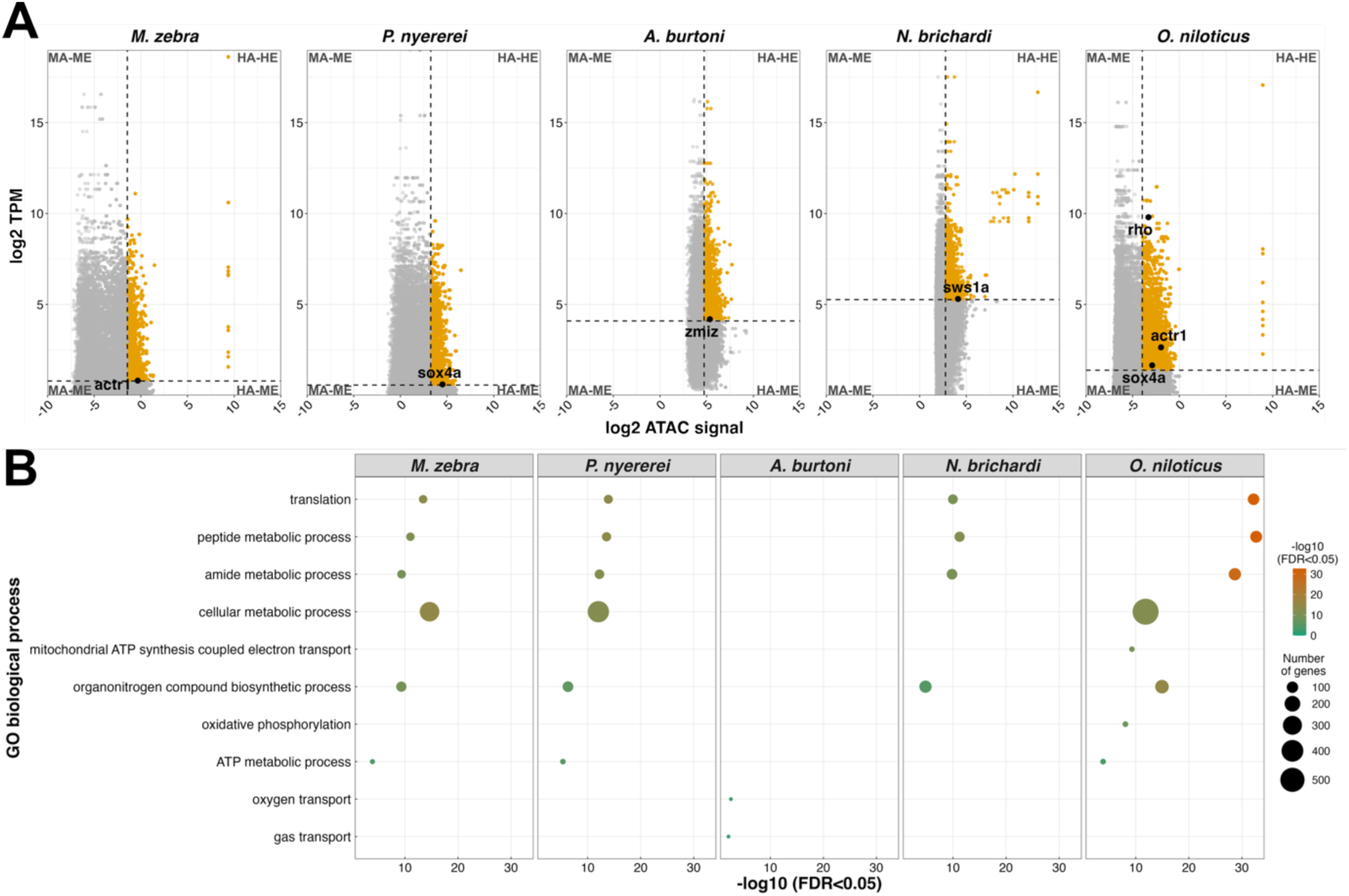
Accessible promoter peaks accounting for gene expression changes in retina tissue across five cichlid species. **(A)** Scatterplots of average *log2* gene expression (as transcripts per million - TPM, y-axis) versus average *log2* ATAC signal (up to 5 kb gene promoter peak counts, x-axis) for 1-to-1 orthologous genes. Genes are categorised by combined accessibility and expression levels into four groups: high accessibility and high expression (HA-HE; both above the 70th percentile, and shown with orange dots), high accessibility and medium-low expression (HA-ME; accessibility above 70th percentile and expression below 50th percentile), medium-low accessibility and high expression (MA-HE; accessibility below 50th percentile and expression above 70th), and medium-low accessibility and medium-low expression (MA-ME; both below 50th percentile). Representative candidate genes are labelled within each plot. **(B)** Gene ontology (GO) biological process enrichment for HA-HE genes across species. Ten selected enriched GO terms (–log10 FDR > 2.1; full set in Supplementary Fig. S23), with circles sized by number of enriched genes and positioned according to GO term (y-axis) and enrichment significance (–log10 FDR<0.05, x-axis and heatmap).

Examining the HA-HE group in retina tissue revealed significant positive correlations in *M. zebra* (R=0.1, *p*<2.2e-16), *N. brichardi* (R=0.08, *p*=6.9e-08), and *O. niloticus* (R=0.2, *p*<2.2e-16), while *P. nyererei* (R=-0.03, *p*=8.5e-04) showed weak negative correlation and in *A. burtoni*, no correlation (Supplementary Fig. S18–S22), paralleling trends in other tissue regulatory categories (Supplementary Fig. S19–S20). Such patterns, as also found in *O. niloticus* gill ^28^, indicate that HA-HE genes are consistent with activated, repressed, or poised states in retina tissue across the five species however, this would need to be validated with relevant histone mark data. Functional enrichment (BH ^43^ adjusted *p*<0.05) among these 902–2,020 HA-HE retina genes identified both shared (e.g., cellular metabolic process in *M. zebra, P. nyererei, O. niloticus*) and species-specific (e.g., oxygen transport in *A. burtoni*, mitochondrial ATP synthesis in *O. niloticus*) GO terms (Fig. 5c, Supplementary Fig. S23), highlighting critical processes for retinal function and visual evolution in fishes ^62^. The HA-HE regulatory group captured key vision-associated genes (Fig. 5a), such as the retinal regulator *actr1* ^63^ (*M. zebra, O. niloticus*), eye development gene *sox4* ^64^ (*P. nyererei, O. niloticus*), eye morphogenesis gene *zmiz* ^65^ (*A. burtoni*), the dim-light vision gene *rho* (*O. niloticus*) with regulatory TFBS divergence ^24^ (Fig. 4e), and the visual opsin gene *sws1* (*N. brichardi*), linked to ecological GRN rewiring in cichlids ^37^.

In summary, conserved and lineage-specific patterns of chromatin accessibility in promoter regions are associated with the transcriptional activity of genes linked to adaptive traits, suggesting that regulatory divergence at the epigenetic level could contribute to visual system differences and adaptation in East African cichlids.

### Motif-supported GRNs reveal pervasive regulatory rewiring

As previous studies have demonstrated that rapid evolution of regulatory elements ^23^ and GRN rewiring ^24^ may contribute to cichlid diversification, we built on this—integrating the matched chromatin accessibility and transcriptional data—to infer tissue- and species-specific GRNs that could dissect regulatory relationships underlying adaptive divergence across the five cichlid species (see ‘*Materials and Methods*’). GRNs were first reconstructed using GENIE3 ^66^, a random forest-based method that frames GRN inference as a set of regression tasks to predict each gene’s expression from all others, yielding ranked TF–target gene edges by variable importance, without requiring prior knowledge of interactions. We selected GENIE3 for its strong performance on benchmark datasets ^66^ and then incorporated motif footprints from accessible promoter regions as an orthogonal filter for regulatory support.

Using expression data alone, large tissue-specific networks were reconstructed (ranging from ∼117k to 11.6M edges per species-tissue combination; Supplementary Table S17a), but the top 12,000 edges per network—the minimum number allowing fair comparison across all datasets without exhausting smaller ‘filtered’ networks below—yielded mostly species-specific structure (Supplementary Fig. S24). GRN rewiring analysis (see ‘*Materials and Methods’*) identified that, of the non-redundant top edges, 97.6–99.7% (47,708–59,373 edges) were species-specific across tissues, 0.3–2.4% (146–1,380 edges) shared in subsets of species, and none conserved across all five species (Fig. 6a). Based on gene expression edges only, Jaccard similarity matrices confirmed low GRN top-edge overlap between species (all <0.03; Supplementary Fig. S25), while multidimensional scaling (MDS) ordinations revealed tissue-dependent separation in GRN space, with mostly different species clustering across tissues (forebrain–*M. zebra/A. burtoni/N. brichardi*, retina–*M. zebra/P. nyererei*, liver–*M. zebra/A. burtoni*, testis– *M. zebra/A. burtoni*; Supplementary Fig. S26).

**Figure 6.**
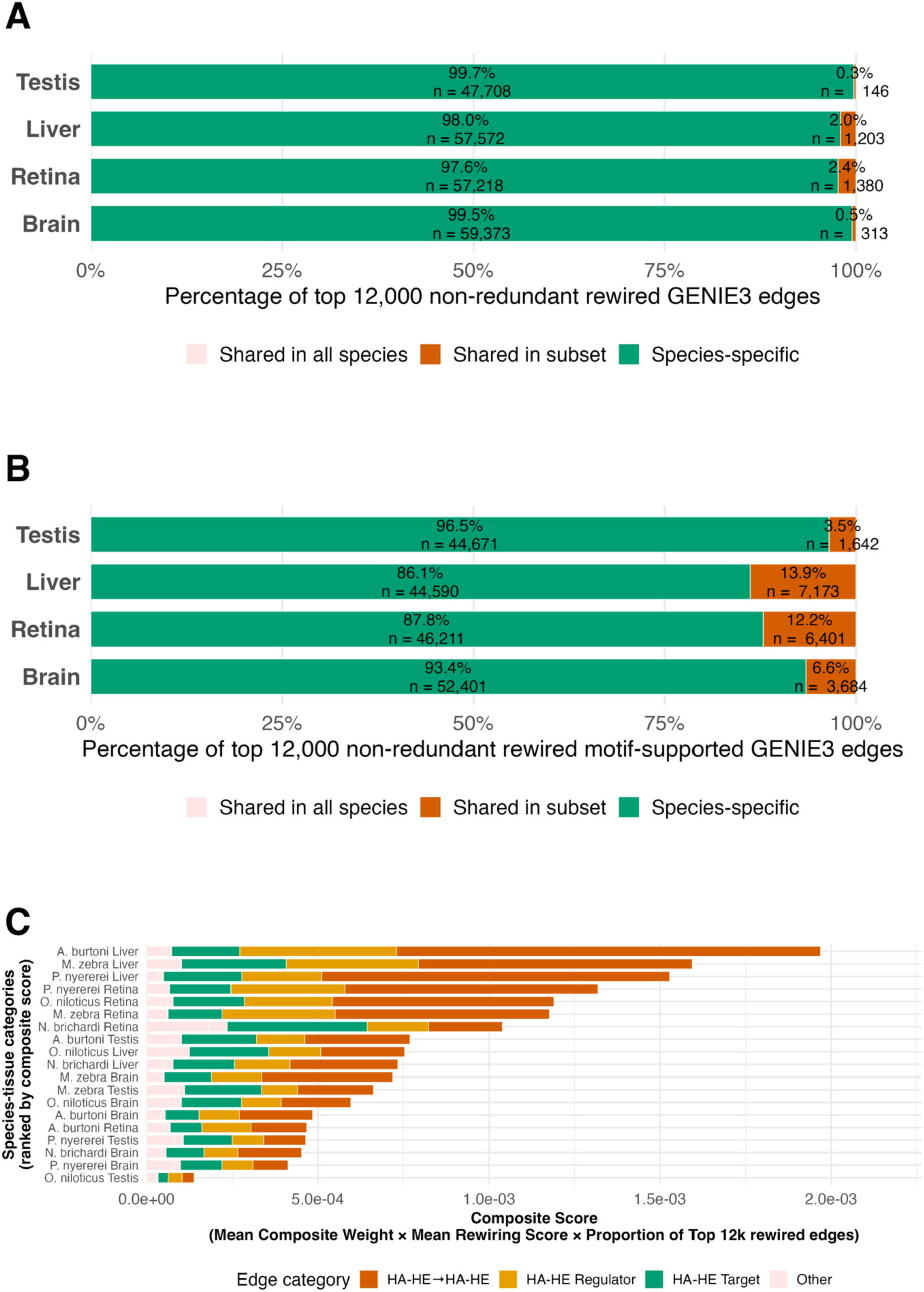
Species-specific and HA-HE-associated GRN rewiring across tissues and ranked composite scores in the five cichlids. **(A)** Distribution of the top 12,000 non-redundant GENIE3 rewired edges across Forebrain, Retina, Liver and Testis, showing the proportion of species-specific edges (green) and the smaller fractions shared in subsets of species (orange) or conserved across all species (pink). **(B)** Distribution of the top 12,000 non-redundant motif-supported GENIE3 rewired edges across the same four tissues, highlighting the same enrichment for species-specific rewiring (green) and the limited share of edges shared in subsets (orange) or across all species (pink). **(C)** Ranked composite score plot of species-tissue categories (y-axis) based on mean composite weight × mean rewiring score × proportion of top 12k rewired edges (x-axis), showing that the highest-ranked categories are dominated by HA-HE-associated edges. The stacked bars distinguish HA-HE → HA-HE (orange), HA-HE regulator (yellow), HA-HE target (green), and other (pink) categories, and the ranking is shown for all 96 species-tissue categories.

Refining networks by motif footprints—normalising maximum orthologous motif bit-scores and computing composite weights (GENIE3 importance × normalised motif bit-score, see ‘*Materials and Methods’*)—identified ∼14k to 1.5M edges across species tissues, yielding sparser (7–11-fold reduction), motif-supported GRNs compared to expression only networks (Supplementary Table S17b). Jaccard similarity (all <0.13; Supplementary Fig. S27) and MDS ordinations identified clearer phylogenetic patterns of conserved regulatory edges (e.g., liver–*M. zebra/P. nyererei/ A. burtoni*) and tissue-specific divergence (e.g., *P. nyererei* and *O. niloticus* are isolated in forebrain and testis GRN space, while *A. burtoni, N. brichardi* and *O. niloticus* are separated in retina; Supplementary Fig. S28). GRN rewiring analysis showed a 5–12-fold increase (compared to expression only GRNs) in shared-subset (1,642–7,173) edges across tissues however, most non-redundant top edges, 86.1–96.5% (46,314–56,085 edges), are species-specific (Fig. 6b); this is supported by ranking rewired edges by species-specificity, weight delta, and rank advantage relative to orthologs (see ‘*Materials and Methods’*; Supplementary Fig. S29-S32).

To identify candidate edges associated with regulatory divergence, we classified GRN orthologs by chromatin accessibility (high [HA]/medium-low [MA]) and expression (high [HE]/medium-low [ME]) status in each species-tissue context, then tested for enrichment among all rewired edges (see ‘*Materials and Methods’*). Edges where both regulator and target exhibit high accessibility and expression (HA-HE → HA-HE), were lowly enriched in only two samples (1.1-fold in *A. burtoni* forebrain, BH ^43^ adjusted *p* = 1.4 × 10^-26^; 1.4-fold in *O. niloticus* testis, BH ^43^ adjusted *p* = 8.4 × 10^-90^), and MA-ME-only edges in only five samples (1–1.3-fold, BH ^43^ adjusted *p* < 0.01) (Supplementary Table S18; Supplementary Fig. S33). Meanwhile, HA-HE Regulator and HA-HE Target edges also showed no to low enrichment in 11 (1–1.4-fold, BH ^43^ adjusted *p* < 10^-4^) and 12 (1–1.3-fold, BH ^43^ adjusted *p* < 0.04) samples respectively, with the highest mean enrichment in HA-HE Target edges of retina tissue (1.2-fold across four species, BH ^43^ adjusted *p* < 5.4 × 10^-8^) (Supplementary Table S18; Supplementary Fig. S33). However, all HA-HE associated edges (as either or both regulator and target) consistently occupy the upper tails of composite weights and rewiring scores across all rewired edges (Supplementary Fig. S34-S37), and when ranking based on a composite of these scores and proportion of top 12,000 species-tissue rewired edges occupied by category, the top 30 (out of 96 species-tissue categories) are all HA-HE associated edges (Fig. 6c; Supplementary Table S19).

The overall top 10 species-tissue categories are HA-HE associated edges in liver (*A. burtoni, P. nyererei, M. zebra*), retina (*P. nyererei, O. niloticus, M. zebra, N. brichardi*), and forebrain (*M. zebra*) (Fig. 6c; Supplementary Table S19), highlighting the importance of dual high accessibility/expression signals in motif-supported GRNs and their potential rewiring for functions of these tissues.

All rewired HA-HE → HA-HE edges were then prioritised to identify the top 50 candidates by composite weight × rewiring score, species-specificity (all unique to one species-tissue) and strong motif support (mean motif bitscore = 14.4, range 11–23). Within this set, dominated by only retina (27 out of 50) and liver (23 out of 50) edges, we identified candidate rewired edges that could be associated with regulatory divergence of tissue-specific traits of the five species, including *O. niloticus* retina CEBPA-*mgme1* (composite weight = 0.052, rewiring score = 13.8) which could be linked to retinal morphology ^67^ and *A. burtoni* liver HNF1A*-foxk1* (composite weight = 0.068, rewiring score = 8.7) that could be linked to liver metabolic processes ^68^ (Supplementary Table S20).

Overall, motif-supported GRNs exhibit rapid regulatory evolution, with 86–97% of top edges being species-specific and thus, rewired along the phylogeny, yet tissue-dependent phylogenetic signal is retained in shared edges. High accessibility and expression (HA-HE) edges—integrating chromatin accessibility, transcriptional activity, and motif support—are consistently rewired across tissues, identifying a focused set of candidate edges that could be linked to tissue function across the species. Building on the prior observation that chromatin accessibility patterns predict transcriptional divergence in adaptive trait genes, these GRN results prioritise regulatory relationships that could contribute to ecological adaptations at the network level.

### Genetic variation in active TFBSs segregates with phylogeny and ecology in East African cichlids

To interrogate functional regulatory divergence, we genotyped TFBS variants of HA-HE and GRN rewired genes across the five cichlid species in deep phylogenies of related riverine and lacustrine species. Our aim was to identify orthologous gene promoter variants that reflect niche- or clade-specific adaptation. To prioritise 1-to-1 orthologous genes, we ranked TFBSs in promoter regions using high tag counts (TC, number of reads) and bit-scores (BS), selecting statistically significant sites above the species-tissue mean (BH ^43^ adjusted *p*<0.0001; see ‘*Materials and Methods*’).

In retina tissue, we identified 896,634–2,247,267 unique TFBSs in 93,139–226,058 promoter footprints, spanning 13,774–21,553 genes across the five species. Of these, 108,193–354,906 (10–20%) TFBSs in 18,726–59,883 (15–32%) footprints within 6,844–13,886 (41–76%) genes exhibited above-mean TC and BS (Fig. 7a, Supplementary Table S21), comparable to findings in *O. niloticus* gill ^28^. Within this candidate set, we focused on TF–target gene (TF-TG) relationships relevant to vision and ecological adaptation ^59^, especially where TFBS variants correlated with transcriptional divergence, e.g., *O. niloticus* RUNX1–*rho*, *P. nyererei* USF2–*pitpn*, *N. brichardi* NR2F6–*sws1* (Fig. 7a).

**Figure 7.**
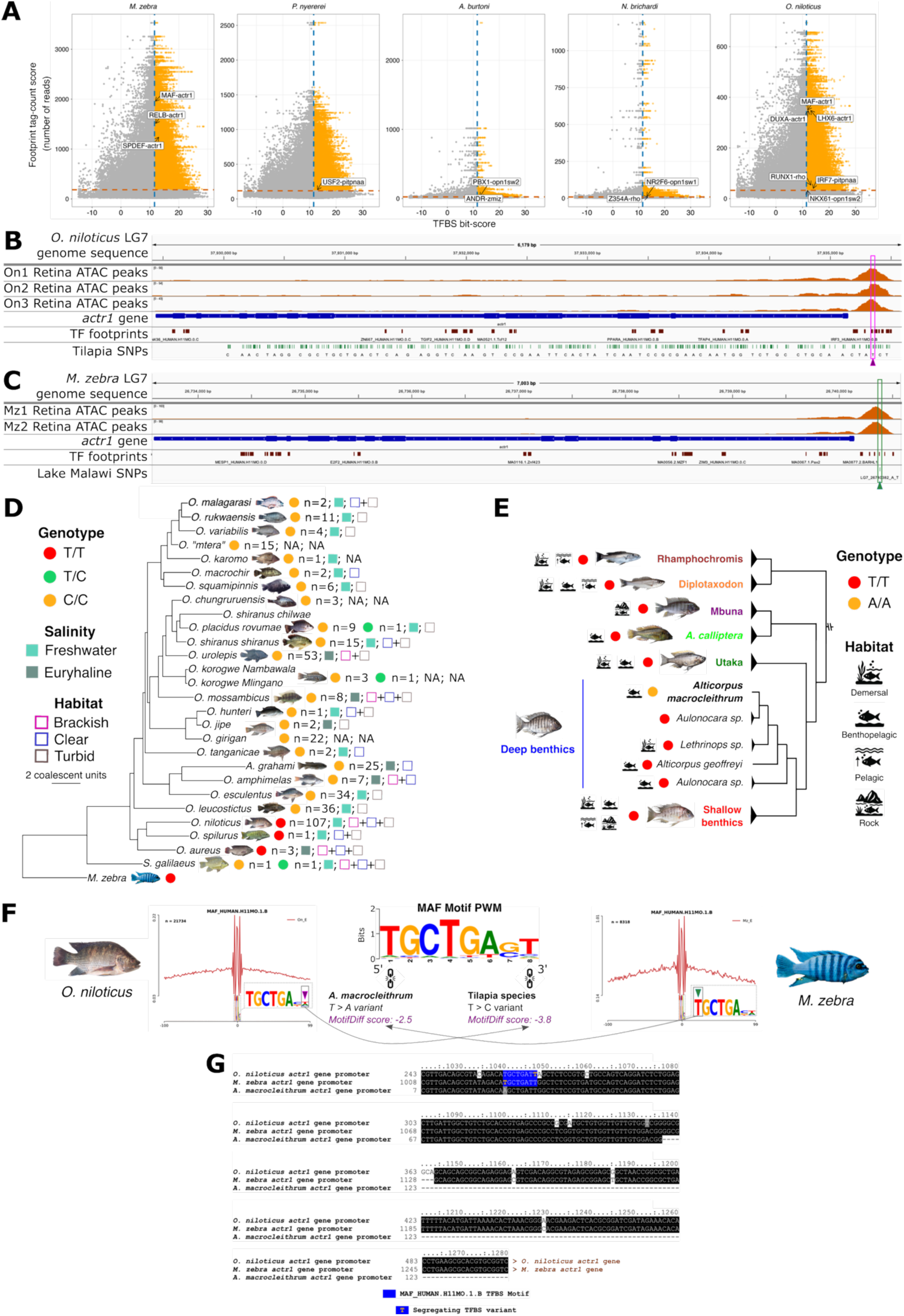
Chromatin accessibility and MAF transcription factor motif variation in the *actr1* promoter of *O. niloticus* and *M. zebra*. **(A)** Scatterplot of tag-count (TC) scores of transcription factor (TF) footprints (y-axis) versus motif bit-scores of predicted TF binding sites (x-axis) detected in accessible retina chromatin. TFBSs exceeding the mean TC (orange dotted line) and mean bit-score (blue dotted line) are shown in orange, while those below these thresholds are shown in grey. Representative TF–target gene candidates are highlighted with arrows. **(B)** Genome track of *O. niloticus actr1* on LG7 with gene annotations (blue), accessible chromatin peaks across three retina replicates (orange), TF footprints (maroon), tilapia SNPs (green), and a polymorphic MAF motif site (pink box/arrow). **(C)** Corresponding *M. zebra* track showing two retina replicates (orange), TF footprints (maroon), Lake Malawi cichlid SNPs (green), and polymorphic MAF motif site (green box/arrow). **(D)** SNP genotypes at the site in (B) mapped onto the tilapia phylogeny ^28,70^: homozygous reference (red), alternate (green), and heterozygous (yellow). Species’ habitats and salinity adaptations annotated. **(E)** SNP genotypes at the site in (C) across the Lake Malawi phylogeny ^69,93^: homozygous reference (red) and alternate (yellow), with habitats indicated. Expanded genotype-phenotype-ecotype phylogeny in Supplementary Fig. S38. **(F)** MAF footprint profiles in *O. niloticus* (left) and *M. zebra* (right), showing signal intensity (red line) centered on footprint (position 0, x-axis). Sequence variation from panels A (purple arrow) and B (green arrow) indicated, with *MotifDiff* score of these variants indicated on input position weight matrices (PWM) of MAF motif used for prediction shown centrally. **(G)** Predicted MAF motifs (blue shading) and polymorphic sites (orange text) in *O. niloticus* and *M. zebra actr1* gene promoter. Orthologous alignment with *A. macrocleithrum* shown below with corresponding *M. zebra* MAF site substitution shown in light grey shading. Additional alignments across other Lake Malawi cichlid genomes are shown in Supplementary Fig. S39.

To assess variant segregation across deeper cichlid phylogenies, we overlapped pairwise variants in highly accessible (HA) TFBSs in promoters of highly expressed (HE) genes of the five cichlids, with population SNP data from 73 Lake Malawi species (134 individuals) ^69^ and 21 other tilapia species (575 indviduals) ^70^, respectively (see ‘*Materials and Methods*’). Notable divergent TF–TG pairs include MAF–*actr1*, SPDEF–*actr1*, and RELB–*actr1* in Lake Malawi’s *M. zebra*; PBX1–*sws2* in the riverine and Lake Tanganyika species *A. burtoni*; and RUNX1–*rho*, MAF–*actr1*, LHX6–*actr1*, and DUXA–*actr1* in *O. niloticus*. Focusing on the photoreceptor gene *actr1*, which is under divergent selection in Lake Malawi’s twilight zone ^71^ and predicted to be regulated by the lens-associated ^72^ TF MAF, we found that the active MAF binding site is positionally conserved in *actr1* promoters of *O. niloticus* (Fig. 7b) and *M. zebra* (Fig. 7c), yet exhibits distinct segregating variants: within tilapia, the last motif position harbours a T/T allele conserved in *O. spilurus* and *O. aureus* but heterozygous or C/C segregating in 21 ecologically diverse tilapia species (Fig. 7d); while in Lake Malawi cichlids, the first motif position shows T/T conservation in 66 species but a homozygous A/A variant segregates in the deep benthic *Alticorpus macrocleithrum* (Fig. 7e-g, Supplementary Fig. S39; see ‘*Supplementary information’*), a species occupying a specialised ecological niche, with unique sensory and morphological adaptations ^73^.

Both variants are predicted to disrupt MAF binding (MotifDiff scores: tilapia −3.8, *A. macrocleithrum* −2.5; Fig. 7f; Supplementary Table S22–S23), reducing the likelihood of canonical MAF regulation in these lineages. Conversely, closely related tilapias (*O. niloticus, O. spilurus,* and *O. aureus*) and most Lake Malawi species retain the ancestral motif and bind MAF, supporting conserved MAF-*actr1* photoreceptor regulation across habitats. The observed patterns support niche-specific divergence, whereby TFBS variants in *actr1* are associated with either clade-level conservation or ecological specialisation. Phylogenetic independent contrast (PIC) analysis shows minimal correlation change after accounting for phylogeny (Supplementary Fig. S40; see *‘Materials and Methods’*), reinforcing that segregating variants could reflect true lineage and niche adaptation.

In summary, our results show that functionally active TFBSs in visual system genes segregate by both phylogeny and ecological niche across radiating and non-radiating cichlid lineages. Distinct TFBS variants in conserved regulatory elements—such as at the *actr1* promoter—are recurrent in ecologically specialised species across broad taxon sampling (73 Lake Malawi species, 21 tilapias; Fig. 7d-f, Supplementary Fig. S38–S39, indicating that regulatory variation could be a key contributor to adaptive innovations and rapid phenotypic diversification in East African cichlids.

## Discussion

Our study provides the first comparative analysis of chromatin accessibility and GRN divergence in East African cichlid fishes, offering novel insights into the molecular basis of cichlid adaptive phenotypic diversification. Building on previous genomic, transcriptomic, and epigenomic work in cichlids ^24,26,28–30,37^, we demonstrate that dynamic regulatory variation is a major contributor to rapid lineage- and niche-specific adaptation in this iconic vertebrate radiation.

Evolutionary change in non-coding regulatory sequences is increasingly recognised as a central driver of phenotypic diversification, particularly during rapid radiations ^24,25,29,30,37,74^. Our findings support and extend this view, revealing both deep conservation and pronounced divergence in open chromatin regions and TF binding site (TFBS) landscapes across East African cichlid species, tissues, and broader ecological clades. Thousands of promoter-associated accessible sites are conserved across all five species and four tissues, yet over half exhibit signatures of accelerated nucleotide evolution, indicating that even constrained elements are modified or co-opted in some lineages—paralleling observed patterns in other vertebrates ^14,75,76^. Divergence is often tissue-specific, notably in retinal visual systems genes—a key trait in cichlid diversification ^19^.

Most TFBSs in promoters of 1-to-1 orthologous genes are not perfectly conserved in sequence or position, showing high turnover along the phylogeny. The *actr1* photoreceptor gene illustrates this, with independent substitutions in different phylogenetic groups altering predicted MAF TF binding probabilities in patterns consistent with niche-specific shifts. Such variation offers a mechanism for GRN rewiring, whereby changes in regulatory connectivity, i.e., TF–target gene relationships, alter transcriptional outputs and potentially adaptive phenotypes, consistent with previous functional studies in cichlids ^24,25,37,74,77^. Given that TFBS annotations here rely on experimental footprinting of vertebrate motifs from several reference databases ^44,45,78,79^, turnover patterns imply potential evolutionary novelty that need to be functionally validated; complementary mobility shift ^37^ and high-throughput reporter ^80^ assays will be essential to confirm their regulatory significance.

Integrating chromatin and transcriptomic profiles, gene promoter accessibility correlates significantly—though often weakly—with gene expression in most within species–tissue combinations, consistent with findings in *O. niloticus* gill tissue ^28^. This is expected as accessible regions do not always correlate with active transcription ^81,82^, as they can mark repressed or poised genes; confirmation of such states requires complementary ChIP-seq assays (H3K27me3, H3K4 methylation) ^83^. Our analyses are necessarily promoter-centric: although intronic and distal peaks display broadly similar accessibility patterns across species and tissues, their assignment to specific target genes remains lower-resolution and thus, definitive mapping will require 3D chromatin-contact data such as Hi-C to supplement GRN inferences. Still, highly accessible promoters of highly expressed genes are enriched for functions such as retinal physiology, metabolism, and oxygen transport—processes linked to coordinated evolution of eye morphology and visual pathways in Lake Malawi niche specialisation ^84^. This combined evidence links chromatin divergence not only to molecular evolution, but to observed ecological and morphological diversification at the level of candidate regulatory networks.

Our GRN analyses extend these observations by showing that motif-supported network inference captures a more resolved layer of regulatory divergence. Compared with expression-only networks, incorporating promoter TF footprints reduced network complexity and increased the interpretability of species-specific rewiring, while retaining tissue-dependent patterns of conservation and divergence. This approach prioritised candidate TF–target relationships supported by both transcriptional evidence and accessible chromatin, rather than co-expression alone, and therefore provides a stronger basis for identifying direct regulatory changes likely to underlie adaptive phenotypes. In particular, the enrichment of rewired high-accessibility, high-expression edges in all tissues suggests that network-level regulatory changes are associated with ecological and functional diversification in cichlids e.g., retinal morphology ^59^ and liver metabolic processes ^29^. These findings build on our earlier co-expression ^37^ and *in-silico* motif prediction ^24,37^ based GRN studies by adding empirical epigenetic support for inference of regulatory interactions, highlighting how promoter accessibility and TFBS turnover could jointly shape lineage- and tissue-specific GRN evolution in a complex vertebrate system.

While retina tissue was prioritised due to its established role in cichlid sensory adaptation with well-characterised ecological gradients across light enrvironments ^59^, parallel analyses across forebrain, liver, and testis datasets reveal broadly comparable patterns of tissue-specific divergence and chromatin-accessibility/transcriptional correlations. These datasets enable analogous analyses to those performed for retina, including TFBS variant segregation across phylogenies and deeper analysis of rewired regulatory connections in tissue-specific contexts (e.g., genes associated with neurotransmission in forebrain, metabolism in liver, reproduction in testis). This can extend previous studies like, for example, divergence in regulatory regions could predict differences in brain structure and behaviour tied to cichlid ecological adaptations ^85^. Future analyses of our forebrain data could also connect active regulatory sites to genomic divergence, testing whether similar signatures drive adaptive changes in forebrain and behaviour, revealing broader tissue-specific mechanisms of ecological and phenotypic novelty.

Although the results presented here offer the most comprehensive cross-species view of chromatin accessibility and regulatory connection evolution in cichlids to date, some caveats remain. To overcome challenges of optimising untested combined epigenetic and transcriptional protocols in cichlid species, samples comprised lab-reared individuals where developmental stage was controlled where possible by selecting adults in breeding coloration with replicates from the same family and similar age. Differences in maturation timing and coloration among species make precise matching difficult, and we cannot exclude effects of environment or captive breeding on genome dynamics, particularly in forebrain and reproductive tissues. However, all five focal species were reared under standardised conditions comparable to those used in other major genomic and transcriptomic studies on the same cichlid species ^23,24,37^. With these established protocols, future work could involve deeper taxon sampling beyond the five species, which were selected based on availability of genomic resources^23,86,87^, phylogenetic coverage as ‘representative’ lacustrine and riverine species ^23^, and ecological interest ^19^, as well as sampling both sexes and along key developmental stages to track regulatory dynamics over time. Also, while sampling bias was minimised through careful tissue matching, high cell numbers (∼50,000 nuclei per replicate), and stringent peak reproducibility (IDR<0.05), undetected subpopulation differences are possible. In this respect, single-cell joint RNA and chromatin profiling ^88^, combined with emerging models of *cis-*regulatory syntax ^89^, will be important for resolving cell type-specific GRNs and their evolutionary trajectories across lineages.

Further, we acknowledge that some open chromatin peaks classified as species-specific may reflect unresolved alignment of orthologous regulatory regions, local assembly gaps, or lineage-specific indels rather than true gain or loss of regulatory activity. Whilst 1-to-1 ortholog filtering, IDR-based peak calling, and peak-overlap thresholds can reduce these effects, they cannot fully exclude them, so species-specific peaks should be viewed as strong candidates for regulatory novelty rather than unambiguous proof of it. Consistent with this caveat, our open chromatin peak calling versus assembly-quality analysis (see *‘Materials and Methods’*) showed that species with higher-N50 assemblies, notably *O. niloticus* ^86,90^ and *M. zebra* ^86,87^, had fewer gap-associated peaks in the raw comparison (Supplementary Fig. S41), but the adjusted analysis suggested that this pattern largely reflects lower gap burden in the more contiguous assemblies rather than a strong N50 effect independent of gaps (Supplementary Fig. S42). Importantly, the absolute proportion of peaks overlapping gap-masked sequence was low overall, indicating that for most peaks in this study, assembly state is unlikely to be a major driver of the observed lineage-specific chromatin landscape; nevertheless, these results underscore the importance of telomere-to-telomere assemblies for representative cichlid species to further reduce ambiguity in interpreting regulatory novelty. This is all the more relevant in rapidly evolving non-coding regions, where sequence divergence can obscure orthology even when regulatory function is conserved, as shown in recent vertebrate *cis-*regulatory element comparisons ^91^.

Since matching species and tissues are unavailable in published methylomes ^29,30^ and transcriptomes ^26^, we have been unable to integrate our atlas with these datasets reliably. Future integration of comparable data could nevertheless enable preliminary testing for anti-correlated marks associated with gene repression, while mapping transposable element insertions within accessible chromatin could help assess their contribution to regulatory novelty ^92^. Expanding this framework across Lake Malawi cichlid clades, including *A. calliptera*, mbuna, benthics, *Diplotaxodon*, and *Rhamphochromis*, now supported by chromosome-level genome assemblies ^93^, will enable testing of whether GRN innovations identified here and elsewhere ^24,37^, together with segregating structural changes ^93,94^, particularly of sensory systems ^93^, could underpin fine-scale ecological diversification.

Overall, our results demonstrate the patterns of promoter accessibility, accelerated sequence evolution, and TFBS turnover that likely shape phenotypic innovations in East African cichlids. By moving beyond cataloguing sequence change to directly measure and annotate regulatory activity, albeit primarily in gene promoter regions, we provide a mechanistic between *cis-*regulatory evolution and genes associated with adaptive traits in both radiating and non-radiating cichlid lineages. This framework sets the stage for future work disentangling adaptive change from relaxed evolutionary constraint, and functional dissection e.g., high-throughput reporter ^80^ and CRISPR assays ^95^ of how specific regulatory changes drive phenotypic outcomes, deepening our understanding of how (epi)genetic forces drive biodiversity at rapid evolutionary timescales.

## Materials and Methods

### Tissue dissection

All animal procedures were approved by the relevant university and carried out in accordance with approved guidelines. Male *Metriaclima zebra, Pundamilia nyererei* (Makobe)*, Astatotilapia burtoni*, *Neolamprologous brichardi* and *Oreochromis niloticus* individuals (Supplementary Table S1) were sacrificed according to either Home Office (UK) or cantonal veterinary permit nr. 2317 (Switzerland) schedule 1 killing using overdose of MS-222 (tricaine) at The University of Hull, UK; University of Stirling, UK; and University of Basel, Switzerland. Four tissues (brain – forebrain specifically, eye – retina specifically, liver and testis) were dissected from each individual for DNA and RNA extraction. Selection of the four tissues enabled the study of tissue-specific associated traits under natural and/or sexual selection in cichlids: forebrain (behaviour, learning and memory, and social interaction); retina (adaptive water depth/turbidity vision); liver (hepatic function associated with dietary differences); and testis (sexual systems associated with behaviour).

### Cell preparation

From a pool of all tissue cells, around 50,000 random cells were harvested, nuclei counted and then spun at 500×g for 5 min at 4°C. Cells were washed once with 50 μL of ice-cold 1× PBS buffer and pelleted at 500×g for 5 min at 4°C. Cell pellets were gently resuspended in 50 μL of 0.05% cold lysis buffer (10mM Tris-HCl pH7.4, 10mM NaCl, 3mM MgCl_2_, 0.05% IGEPAL CA-630), and immediately pelleted at 500×g for 10 min at 4°C and then stored on ice.

### Transposition reaction and purification

Pelleted nuclei were gently resuspended in a 50 μL transposition reaction mix composed of 25 μL 2× TD Buffer (Illumina), 2.5 μL Tn5 Transposes (Illumina) and 22.5 μL Nuclease-Free H_2_O. Transposition reaction was incubated at 37°C for 30 min and purified using the PCR purification MinElute Kit (QIAGEN), according to manufacturer’s protocol. Transposed DNA was eluted in 10 μL Elution Buffer (10mM Tris buffer, pH 8) and stored at −20°C.

### PCR amplification

Transposed DNA was amplified in a final volume of 50 μL composed of 10 μL Transposed DNA, 10 μL Nuclease Free H_2_O, 2.5 μL 25μM P7 adapter, 2.5 μL 25μM P5 adapter and 25 μL NEBNext High-Fidelity 2x PCR Master Mix (NEB). Transposed DNA was amplified for 5 min at 72°C, 30 secs at 98°C, and 11 cycles of 10 secs at 98°C, 30 secs at 63°C and 1 min at 72°C in a PCR thermocycler.

Amplified transposed DNA was purified using the PCR purification MinElute Kit (QIAGEN) and eluted in 20 μL Elution Buffer (10mM Tris buffer, pH 8), according to manufacturer’s protocol. To remove excess primers for final ATAC libraries, an additional 1× Agencourt AMPure XP bead (Beckman Coulter) clean-up was performed and eluted in 0.1× filtered TE. ATAC libraries were quantified on the Qubit 4 fluorometer (Invitrogen) and size distribution assessed on Agilent Tapestation and/or Bioanalyser.

### DNA extraction

DNA was purified from the pool of all cells of each tissue using the DNeasy Blood and Tissue kit (QIAGEN) according to manufacturer’s protocol. DNA was quantified on the Nanodrop 2000 (Thermo Scientific) and Qubit 4 fluorometer (Invitrogen), and used 1 ng for DNA library preparation using the Nextera XT DNA Library Preparation kit (Illumina), according to manufacturer’s protocol. Library size distribution was assessed on Agilent Tapestation and/or Bioanalyser. DNA libraries, obtained from naked DNA, were used as internal controls to determine background levels of genomic DNA accessibility and Tn5 transposase sequence cleavage bias.

### ATAC and control DNA sequencing

51 ATAC and 51 corresponding naked DNA control libraries (Supplementary Table S1) were equimolar pooled and 50bp paired-end sequenced at Earlham Institute on 2 lanes of the Illumina NovaSeq 6000 platform using an S2 flow cell, generating an average of 61 million (ATAC-seq) and 16 million (naked DNA control) reads per library.

### ATAC-seq and control DNA processing

Sequence adaptors were removed and trimmed for quality from all raw paired-end reads using Trim Galore! (v0.5.0; https://github.com/FelixKrueger/TrimGalore) and FastQC (v0.11.9) ^96^ using default parameters. Read alignment, post alignment filtering and ATAC peak calling were performed according to the ENCODE projects ‘ATAC-seq Data Standards and Processing Pipeline’ for replicated data (https://www.encodeproject.org/atac-seq/). To minimise false inflation of species-specific regulatory peaks arising from divergent or repetitive sequence, we stringently mapped trimmed reads to their respective genome - *M. zebra* UMD2a ^86,87^; *P. nyererei* v1 ^23^; *A. burtoni* v1 ^23^; *N. brichardi* v1 ^23^; *O. niloticus* UMD_NMBU ^86,90^ using bowtie2 (v2.2.6) ^97^, with parameters `‘–k 4 –X2000 –mm`’, retained only high-confidence, properly paired mappings following the ENCODE ATAC-seq pipeline, and outputted in BAM format using SAMtools (v1.9) ^98^. Since ATAC-seq can generate a high proportion (15-50% in a typical experiment) of mitochondrial mapped reads, any reads mapping to their respective mitochondrial genomes were identified using BLAST (v2.3.0) ^99^ and removed from the BAM file using SAMtools (v1.9) ^98^. The resulting BAM files were sorted, and duplicated reads were marked using Sambamba v0.6.5 ^100^. Duplicated, unmapped, non-primary alignment, and failing platform QC reads were filtered using SAMtools (v1.9) ^98^, retaining reads mapped as proper pairs, and fragment length distributions were plotted using Picard (v1.140; https://github.com/broadinstitute/picard/). At each step, the recommended parameters from the ENCODE pipeline were applied. BAM files were converted to *tagalign* files using Bedtools (v2.30.0) ^101^ and Tn5 shifting of ATAC mappings carried out prior to peak calling. Peaks were identified using macs2 (v2.1.1) ^102,103^ with the shifted tag as test and corresponding control DNA as input with parameters `‘-f BED -p 0.05 --nomodel --shift -75 --extsize 150`’. Narrow peaks were used to create coverage tracks using *bedClip* and *bedToBigBed* in the UCSC-tools package (v333; http://hgdownload.cse.ucsc.edu/admin/exe/). Following the ENCODE pipeline, Irreproducible Discovery Rate (IDR) peaks of true replicates were flagged as either true (<0.1) or false (≥0.1) using idr v2.0.4 (https://github.com/kundajelab/idr). The fraction of reads in peaks (FRiP) were calculated using Bedtools (v2.30.0) ^101^ and demarcated as pass (>0.3) or acceptable (>0.2) according to ENCODE guidelines. FRiP was not used as a QC measure and instead, transcription start site (TSS) enrichment was calculated using ATACseqQC (v1.18.0) ^104^, with the TSS enrichment QC requirement of a significant signal value at the centre of the distribution being applied. All samples passed this criterion for further analysis. To avoid naïve approaches of merging peaks across replicates ^105^, for each species sample, a consensus set of peaks across all biological replicates was created based on reproducible (IDR true) peaks, where intersecting peaks are also ranked based on MACS2 peaks significance score ^105^. Consensus peaks were used for subsequent analyses.

### Assessing peak calling bias versus assembly quality

We assessed whether open chromatin peak calling was biased by aforementioned genome assembly quality by summarising peak-level statistics across the four tissues for each species. Peaks were counted as gap-associated in genome contigs/scaffolds only if at least 10% of the peak bases were overlapping ‘N/n’ gap regions, and as scaffold-end proximal if they fell within 100 bp of either scaffold end. For each species, these fractions were averaged across tissues, then merged with species-level assembly contiguity (scaffold/contig N50) ^23,87,90,106^ and genome-wide ‘gap fraction’, calculated as the proportion of all bases in the genome FASTA that were N/n’s. We then tested the association between contiguity and peak bias using Spearman correlation, and repeated the analysis with a partial Spearman correlation controlling for ‘gap fraction’ to assess whether any N50 effect was independent of overall gap burden.

### Multiple genome alignment of five cichlid species

A reference-free multiple genome alignment (MGA) of the five cichlid genomes (same versions as above) was created using Cactus (v2.0.3) ^107^ and outputted in Multiple Alignment Format (MAF). This reference-free approach allowed us to define orthologous genomic regions across species, so that cross-species peak comparisons were made only within homologous loci rather than by single-reference coordinate transfer. Species-specific peaks were therefore called only after orthology-aware comparison of aligned regions, reducing the likelihood that apparent species specificity was driven solely by local assembly or mapping artefacts.

### Identifying conserved noncoding elements (CNEs)

A neutral substitution model was created using the *phyloFit* function of PHAST (v1.5) ^108^ by fitting a time reversible substitution ‘REV’ model and parameters ‘—tree “((((Metriaclima_zebra,Pundamilia_nyererei),Astatotilapia_burtoni),N eolamprologus_brichardi),Oreochromis_niloticus)” --subst-mod REV’.

The five cichlid MGA was split by *O. niloticus* chromosome/scaffold using the *mafSplit* function in the UCSC-tools package (v333; http://hgdownload.cse.ucsc.edu/admin/exe/).

The neutral substitution model and MGAs of each *O. niloticus* reference chromosome/scaffold were used as input to predict conserved noncoding elements (CNEs) using the *phastCons* function of PHAST (v1.5) ^108^ with parameters `‘--target-coverage 0.3 --expected-length 30 --most-conserved --estimate-trees --msa-format MAF`’. We used the conservation score output to define highly conserved CNEs (hCNEs) with sequence identity ≥90% over ≥30bp and pseudo accelerate/diverged CNEs (pseudo aCNEs) with sequence identity <90% over ≥30bp. The evolutionary conservation-acceleration (CONACC) score of each pseudo aCNE was predicted by running *phyloP* ^42^ of the PHAST (v1.5) package ^108^ using the likelihood ratio test (LRT) ‘`--method LRT`’ on the CNE `--features` with their corresponding neutral substitution model in ‘`--mode CONACC`’. Pseudo aCNEs with a negative CONACC score, likelihood ratio <0.05, and significant divergence (altsubscale >1), were defined as significantly deviating from the neutral model, and therefore as true aCNEs. Since CNEs were defined by their position in *O. niloticus* chromosomes as a reference, corresponding CNE coordinates in the other species were defined using a combination of the *mafsinRegion* function of the UCSC-tools package (v333; http://hgdownload.cse.ucsc.edu/admin/exe/) and Bedtools (v2.30.0) ^101^.

### Peak annotation and orthology

Up to 5 kb gene promoter regions were annotated in our previous study ^37^ for *P. nyererei* v1, *A. burtoni* v1 and *N. brichardi* v1, with the same method being applied to the *M. zebra* UMD2a and *O. niloticus* UMD1 genomes in this study. We defined ±100 bp of TSS as the ‘core promoter’ and up to 5 kb upstream as the ‘extended promoter’ based on previously observing a plateau of predicted TFBSs from 5–20 kb upstream of the TSS in the same five species (see Fig. S22 in Additional File 1 ^37^). Narrow peaks either overlapping or most proximal to all annotated features in each genome were mapped using the *intersect* function of Bedtools (v2.30.0) ^101^. A presence-absence matrix was generated based on peaks overlapping in the up to 5kb gene promoter regions for each species tissue, and each peak assigned to a gene accordingly. Gene family relationships/orthologs of all genes in the five species were derived using OMA (v2.3.0) ^109^ with Ensembl gene annotations corresponding to the genome versions provided above as input. A total of 38,445 orthologous genes were identified across the five species, with 13,889 1-to-1 orthologous genes found in all five species. Each ‘gene assigned peak’ was then assigned to its corresponding orthologous gene, and for each species tissue, a single representative ‘best’ peak (over all replicates) was assigned to each orthologous gene based on being 1) a True IDR peak; 2) closest to the gene TSS; and 3) most significant merged peak FDR corrected *p-value* (*q-value*). Peak orthology in each species tissue was then determined using the above derived orthologous genes, and used for peak conservation or uniqueness, as well as gain or loss analysis below.

### Peak gain or loss and genetic diversity

Up to 5 kb gene promoter regions of species genes in all 38,445 orthologous genes were aligned using MAFFT v7.271 ^110,111^. Species peak positions were adjusted according to multiple alignment gaps and overlapping peaks were determined using Bedtools *intersect* (v2.30.0) ^101^. All pairwise overlapping peak regions, defined as ‘conserved’ peaks, were required to encompass both species peaks summit, and non-overlapping ‘unique’ peaks as not, for analysis of conservation or uniqueness, gain or loss, and subsequent analysis of genetic diversity. For subsequent analyses we only include *O. niloticus* as the reference and not the comparative species for loss since it is the base of our phylogenetic tree. To avoid any biases based on difference in genome quality and annotation, multiple alignments of 13,889 1-to-1 orthologous genes were used to calculate pairwise sequence similarity of ‘conserved’ overlapping peak regions using an in-house python script (https://github.com/tmehta12/ATAC_bioinformatics) and categorised into bins of 0-1% (highly conserved), >1-5% (moderately conserved), >5-10% (less conserved) and >10% (variable) genetic diversity. These sequence-similarity thresholds reflect the spectrum of selective pressures observed across the five species’ genomes, ranging from highly conserved coding regions (95 - 99.7% identity) to more variable whole-genome sequences (85 - 89% identity) ^24^. For each reference peak within these sets, any pairwise absence of peak overlap is recorded as a loss. For example, a reference peak in *M. zebra* with overlapping peaks and summits in *P. nyererei* and *N. brichardi*, but not *A. burtoni* is recorded as presence between *M. zebra* - *P. nyererei* and *M. zebra* - *N. brichardi*, but a loss between *M. zebra* - *A. burtoni*. To determine species-specific peak gain in *M. zebra, P. nyererei*, *A. burtoni* or *N. brichardi*, we calculated based on two sets of data: 1) a stringent criterion of only zero pairwise peak overlap; and 2) where there is zero pairwise peak overlap or where either one or both species peak summits are not encompassed within the peak overlap region. Gains or losses and genetic diversity of peak overlap were then collated and counted for each tissue.

Genetic diversity in peak adjacent regions across five species was quantified using a custom Python script (https://github.com/tmehta12/ATAC_bioinformatics) applied to 1-to-1 orthologous promoter alignments spanning ±500 bp from tissue-specific peak boundaries, enabling comparison of diversity in peak versus peak-adjacent regions. Diversity estimates were obtained through resampling, with 100 random lines drawn 50 times for peak-adjacent regions of each species tissue, and mean percentage nucleotide identity was calculated per iteration.

### Measuring evolutionary rate of peaks

The evolutionary conservation-acceleration (CONACC) of each annotated *O. niloticus* peak was predicted by running *phyloP* ^42^ in the PHAST (v1.5) package ^108^ using the likelihood ratio test (LRT) ‘`--method LRT`’ on the peak `--features` with their corresponding neutral substitution model in ‘`--mode CONACC`’.

### Testing the significance of CONACC scores of peaks overlapping genomic regions

The significance of accelerated CONACC scores of annotated *O. niloticus* peaks in each genomic region was tested (Wilcoxon rank sum) against one another for a) all peaks, and b) within each pairwise comparison. The *p-value* was adjusted using Benjamini-Hochberg (BH) ^43^ correction, and recorded as either having a significant (Wilcoxon rank sum test, adjusted *p-value* <0.05) or insignificant (Wilcoxon rank sum test, adjusted *p-value* >0.05) difference in accelerated CONACC scores for peaks in each region.

### Gene ontology (GO) enrichment

GO enrichment analyses of peaks (from 1-to-1 orthologous genes) localised to the up to 5 kb gene promoters for each tissue collated by species was conducted using the ‘g:GOst’ module of g:Profiler (https://biit.cs.ut.ee/gprofiler/gost) ^112^, version e111_eg58_p18_f463989d (2024), using the species-specific databases. We use the false discovery rate (FDR) corrected hypergeometric *p* value with Benjamini-Hochberg (BH) ^43^ correction to assess enrichment of GO terms, with a statistical cut-off of FDR < 0.05.

### Transcription factor (TF) footprinting, enrichment, and clustering

TF footprints were characterised using HINT-ATAC in the Regulatory Genomic Toolbox (v0.13.0) ^113^ using a stringent false positive rate (FPR) of 0.0001, with both species-specific and cichlid-wide position weight matrices (PWMs) as defined in our previous study ^37^, as well as vertebrate PWMs from JASPAR (v9.0) ^44,114^, HOCOMOCO ^45^, GTRD ^78^, and UniPROBE ^79^.

The enrichment of accessible peaks in genomic regions was tested using the Genome Association Tester (GAT) tool ^115^. The accessible peaks were provided as segments of interest to test, with a bed file of annotations to test against, and workspace as the length of each scaffold/chromosome. GAT was ran using the following parameters: *–verbose = 5, −-counter = segment-overlap, −-ignore-segment-tracks, −-qvalue-method = BH –pvalue-method = norm*. We use the Benjamini-Hochberg ^43^ false discovery rate (FDR) to assess enrichment of peaks in annotated regions, with a statistical cut-off of FDR < 0.05. Similarly, we use the false discovery rate (FDR) corrected hypergeometric *p*-value (*q-*value) test to assess localised enrichment of TF footprint motifs in open chromatin peaks overlapping gene promoter regions. Enrichment is tested using a set-based approach whereby TF footprints in active gene promoter regions are compared against a background of TF footprints identified in open chromatin peaks in the whole genome.

For clustering TF activities, the mean of each TF’s bit-score of motif match (as a measure of activity) in all gene promoters for all species and tissues was calculated. Using the mean bit score of each motif match, the within-group sums of squares and silhouette coefficient, as a measure of clustering homogeneity, for k = 1 to 20 clusters was calculated and plotted in R (v4.4.2). K-means clustering was selected over hierarchical clustering due to the volume of the data and to observe cluster differences between species, as opposed to a nested tree. The most suitable ‘k’ for clustering was selected based on coherence between the two methods where 1) the sum of squared distance falls suddenly (‘elbow’ method); and 2) the highest average silhouette width that is above 0.25, representative of structured data ^116^. The bit scores were Z-score transformed across rows and heatmap plotted in R (v4.4.2) using the ComplexHeatmap package (v3.6) ^117^. TF footprint line plots were generated with the ‘differential analysis’ module of HINT-ATAC in the Regulatory Genomic Toolbox (v0.13.0) package ^113^ using bias-corrected signals.

### Nucleotide variation in TF motifs

Using a similar method for detecting peak genetic diversity above, TF motif positions were adjusted according to multiple alignment gaps of gene promoter regions and overlapping TF motifs between *O. niloticus* (as a reference) and the other four species were determined using Bedtools *intersect* (v2.30.0) ^101^. Overlapping TF motifs were classified as either 100% overlap or partial overlap, and whether of the same or different TF motif. An in-house python script (https://github.com/tmehta12/ATAC_bioinformatics) was used to identify all pairwise polymorphic sites in gene promoter alignments and using Bedtools *intersect* (v2.30.0) ^101^, polymorphic sites overlapping *O. niloticus* TF motifs were identified with genomic coordinates of reference or alternate alleles recorded.

### RNA extraction and sequencing

RNA was purified from the remaining random pool of cells of each tissue using the RNeasy Plus Mini kit (QIAGEN) according to manufacturer’s protocol. RNA and DNA content were quantified on the Qubit 4 fluorometer (Invitrogen) and integrity assessed on Agilent Tapestation and/or Bioanalyser, taking samples with RIN≥7 and <15% genomic DNA. A total of 48/51 (94%) samples passed these criteria for selection (Supplementary Table S1). A total of 48 stranded RNA libraries were prepared using the NEBNext Ultra II Direction RNA-seq kit according to manufacturer’s protocol. All stranded RNA-seq libraries were equimolar pooled and 150bp paired-end sequenced at Earlham Institute on 1 lane of the Illumina NovaSeq 6000 platform using an S4 v1.5 flow cell, generating an average of 70 million reads per library.

### RNA-seq processing

Read quality was assessed using FastQC (v 0.11.9) ^96^ and Trim Galore! (v 0.6.5; https://github.com/FelixKrueger/TrimGalore) was used to remove adapters and trim for low-quality paired end reads using default settings. All reads were then mapped to their respective genome - *M. zebra* UMD2a ^87^; *P. nyererei* v1 ^23^; *A. burtoni* v1 ^23^; *N. brichardi* v1 ^23^; *O. niloticus* UMD_NMBU ^90^ using HISAT2 (v 2.2.1) ^118^ with default parameters. Mapping QC was carried out using QualiMap (v 2.2.1) ^119^ with default ‘rnaseq’ parameters. The final BAM file was sorted using samtools (v1.16.1) ^98^ and transcript abundance calculated using ‘htseq-count’ in the HTSeq (v 2.0.2) ^120^ package.

### Identifying ATAC peak and gene expression associations

To prioritise peak-gene relationships, we devised an approach to identify peaks that could regulate their target genes expression. For gene expression, we used an approach applied previously ^28,121^ to calculate transcript per million (TPM) values by first normalising transcript counts by gene length, as calculated using GTFtools ^122^, followed by the library size, as calculated using QualiMap (v 2.2.1) ^119^, and then *log2*(x +1) transforming 1) each genes TPM in each replicate; and 2) the mean TPM for each gene across biological replicates. Each genes mean TPM across biological replicates was used to assess correlations with ATAC signal. The ATAC signal was processed using an approach applied previously ^28,123^ to obtain the number of independent Tn5 insertions in collated gene promoter peaks. The number of insertion sites i.e., peak counts, was counted in collated narrow peaks against a merged BAM file of all biological replicates using Bedtools (v2.30.0) *coverage* ^101^. A peak count matrix was created and normalised using edgeR ^124^ counts per million (CPM) ‘log=TRUE, prior.count=5’, followed by a quantile normalization using the preprocessCore (https://github.com/bmbolstad/preprocessCore) *normalize.quantiles* module in R (v 4.4.2). After merging the ATAC signal of biological replicates using the *log2* average from the normalized counts matrices, the average *log2*-TPM and *log2*-ATACSignal for each gene was plotted with the correlation co-efficient (r) calculated in R (v 4.4.2).

Using an approach devised previously ^28,82^, putative activated, repressed or poised genes were categorised based on assigning each peak-gene relationship to one of four groups according to gene promoter accessibility (*log2*-ATACSignal) and gene expression (*log2*-TPM) values that are less than the 50th (medium-low) or more than the 70th (high) percentiles of all peaks counts or genes expression values in the same species tissue. The four groups are (1) HA–HE (high accessibility and high expression); (2) HA–ME (high accessibility and medium–low expression); (3) MA–ME (medium–low accessibility and medium–low expression); or (4) MA–HE (medium–low accessibility and high expression). To maintain stringent gene groups, any genes not falling into any of the four groups were discarded.

### Inference and comparative analysis of tissue-specific gene regulatory networks

Gene regulatory networks (GRNs) were inferred for each species-tissue combination using GENIE3 (v1.30.0) ^66^ on orthologous gene expression data, yielding TF–TG edges ranked by importance weight. To integrate ATAC-seq footprinting evidence, we filtered GENIE3 networks to retain only edges supported by orthologous motif footprints, extracting maximum motif bit-scores. Composite edge weights were calculated by first calculating normalised motif bit-scores (to match GENIE3’s 0-1 range) followed by composite weight calculation:

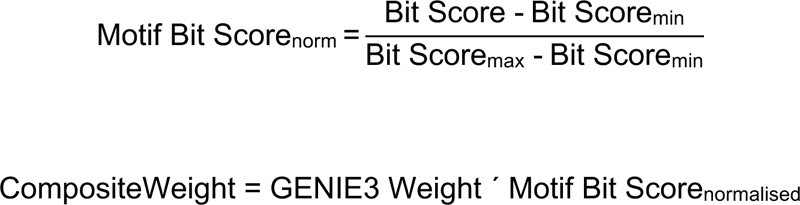

This produced ranked edge lists for each of the species-tissue networks across the five species.

Comparative GRN analysis was performed in R (v4.5.0). For each tissue, the top 12,000 composite-weighted edges per species were selected for Jaccard similarity analysis across species pairs, generating tissue-specific similarity matrices and multidimensional scaling (MDS) ordinations. Rewiring was quantified per species by computing a composite score combining (1) species-specificity (binary: edge absent in other species), (2) composite weight delta relative to orthologous edges, and (3) rank advantage relative to orthologous edges, with top 100 rewired edges identified per species-tissue. All GRN regulators and targets were then classified into accessibility/expression categories (HA-HE, MA-HE, HA-ME, MA-ME), and category enrichment was assessed using one-sided Fisher’s exact tests against the complete set of rewired edges as background, with Benjamini-Hochberg ^43^ false discovery rate (FDR) correction applied within each species-tissue across the four edge classes (FDR < 0.05). For downstream prioritisation, the full distribution of composite weights and rewiring scores across edge categories was examined, and ranked HA-HE candidate edges by a combined score calculated as CompositeWeight × rewiring score.

### Identification of segregating variants in binding sites

Pairwise variants of *M. zebra* and *O. niloticus,* derived from gene promoter alignments above, were overlapped with SNPs in either Lake Malawi ^69,93^ or other tilapia ^70,125^ species respectively, using Bedtools (v2.30.0) *intersect* ^101^. Variants were prioritised based on ordering TF-TG candidates where 1) the TG is HA-HE and a TFBS in the promoter has variation between species; or 2) a TFBS has variation between species; or 3) the TG is HA-HE; and then ordering variants in TFBSs based on ranked reads (tag count, TC) in the footprint and bit-score of the motif match.

### Quantifying variant effect on binding site disruption

The *M. zebra* UMD2a ^87^ and *O. niloticus* UMD_NMBU ^90^ genomes, HOCOMOCO v11 human motifs (in meme format), and chromosome-separated vcf files of Lake Malawi species ^69,93^ or tilapia SNPs ^70,125^ were inputted into *MotifDiff* ^126^ to obtain normalised scores of variant effect to either decrease (score of < 0) or increase (score of > 0) binding probability of all predicted motifs along each chromosome.

### Phylogenetic Independent Contrasts

PICs were carried out to test the effect of fitting species phylogenies to the covariance of segregating TFBSs and species ecologic traits derived from previous studies (habitat ^127,128^ and foraging habit/diet ^127^). This involved (1) coding the genotypes of segregating regulatory sites and ecologic measurements for all species, and (2) using the ape package (v5.8.1) in R (v4.5.0) to apply the PICs test ^129^ on all correlations with the binding site genotype (genotype vs. habitat and genotype vs. foraging habit/diet). PICs assume a linear relationship and Brownian motion process between traits, and thus, for each combination of data, scatterplots were first generated. To test for any correlation changes owing to phylogenetic signal, the regression model was compared between the relationships both excluding and including the respective input species phylogeny.

## Declarations

### Data Availability

The data underlying this article are available in the article, its online supplementary material, or uploaded to Zenodo (https://doi.org/10.5281/zenodo.19821867). All code is available on GitHub at: https://github.com/tmehta12/ATAC_bioinformatics.

### Consent for publication

Not applicable.

## Acknowledgments

The authors acknowledge the support of the Biotechnology and Biological Sciences Research Council (BBSRC), part of UK Research and Innovation. The authors would like to acknowledge Tom Barker, Vanda Knitlhoffer, Leah Catchpole, and Suzanne Henderson of the Genomics Pipelines Group at Earlham Institute for data generation including preparation of RNA-Seq libraries, pooling, and sequencing. The authors would also like to acknowledge the Scientific Computing group, as well as support for the physical HPC infrastructure and data centre delivered via the NBI Research Computing group.

## Author contributions

DJ, AMS, WS and AI sacrificed and dissected fresh fish tissues; TKM prepared tissue-specific cells to perform transposition reactions and purifications for ATAC libraries; TKM and AM performed tagmentation of genomic DNA controls, library amplification and QC; GE ran multiple genome alignments; TKM did all analyses; TKM and WH wrote the manuscript with input from AM, GE, DJ, AMS, AI, WS and FDP.

## Corresponding authors

Correspondence to

