## Supplementary Information for "Evolution of chromatin accessibility associated with traits of cichlid phenotypic diversity"

### Results

#### Cichlid chromatin accessibility is enriched in noncoding regulatory regions

Across all species and tissues, an average of 78–99% of open chromatin peaks occur within 5 kb of a gene's transcription start site (TSS), with 10–20% located in the core promoter region (±100 bp of the TSS) (Fig. 1b, Supplementary Fig. S1, Supplementary Table S2). More specifically, peaks are most strongly enriched in 5′ UTR sequences across all species (fold enrichment = 1.8–4.1, adjusted $P < 0.05$; Fig. 1c). Genome wide, across all species and tissues, open chromatin peaks span several genomic features, overlapping intronic regions (40%), exons (23%), and intergenic regions (17%), with the remainder distributed among promoter regions within 5 kb upstream of the TSS (10%), 5′ UTRs (7%), 3′ UTRs (3%), and conserved noncoding elements (CNEs; 0.04%) (Supplementary Fig. S2, Supplementary Table S3).

#### Core regulatory elements reveal conserved and clade-specific chromatin landscapes

Using the 13,997 1-to-1 orthologous genes across the five species (see '*Materials and Methods*'), we identified 13,889 orthologous genes (99%) with an associated open chromatin peak in the gene promoter region (Fig. 1d). These orthologous genes are enriched for common biological processes such as cell differentiation and epithelium development in each of the four tissues, as well as unique processes such as synapse organisation (in forebrain tissue), axon extension (in retina tissue), myotube differentiation (in liver tissue) and dephosphorylation (in testis tissue) (Supplementary Fig. S3).

In the 13,889 orthologous genes, a total of 5,536 (40%) unique orthologous genes (202-4321 orthologous genes in each tissue) have conserved gene promoter peaks in orthologous genes of all five species (*Anc1* node, Fig. 1d). Examples of which include in forebrain tissue (4321 orthologous genes), *irx2*, a gene associated with neural differentiation and behavioural helper care in a cooperatively breeding cichlid [1]. Examples in the retina tissue (202 orthologous genes) include *atxn1b*, a gene important for visual function [2], and with an overall gain of TFBSs along the same phylogeny [3], has significantly rewired GRNs in cichlids [4]. Along the phylogeny, 1,473 unique orthologous genes (65-1253 orthologous genes in each tissue) have shared peaks in gene promoters of the haplochromines (Anc3, Fig. 1d); in liver tissue (231 orthologous genes), this includes *foxm1,* a gene associated with cell cycle stimulation during liver regeneration [5]. Notably, compared to ancestral nodes, there are fewer (39-188) unique orthologous genes with species-specific peaks in their gene promoters, but variability (36-1762) of orthologous genes with peaks at the per tissue level across species (Fig. 1d); in forebrain tissue (71-513 orthologous genes), this includes *lrrtm1* in *A. burtoni* and *vldlr* in *P. nyererei* associated with modulating synaptic cell adhesion [6] and neural development [7] respectively; in retina tissue (36-1762 orthologous genes), this includes *fgf10* in *M. zebra* and *sws1* in *N. brichardi* that are associated with eye morphogenesis [8] and short wavelength sensitive visual function in cichlids [9] respectively; in liver tissue (73-380 orthologous genes), *cxcr3* in *M. zebra* and *cxcr1* in *O. niloticus,* genes associated with hepatic immune response [10,11]; and in testis tissue (88-439 orthologous genes), *plag1* in *P. nyererei* associated with spermatogenesis [12], and *prdm9* in *A. burtoni* associated with meiotic recombination [13].

**Species- and tissue-specific open chromatin peaks underpin regulatory diversification**

We then focus on the species-specific level where, compared to ancestral nodes (1018-5536 orthologous genes), there are fewer unique orthologous genes (39-188), but tissue-specific variability (36-1762 orthologous genes) of species-specific peaks in gene promoters (Fig. 1d). The number of gene promoter peaks in unique orthologous genes varies between species (Mz = 192-1762 orthologous genes; Pn = 73-413 orthologous genes; Ab = 36-439 orthologous genes; Mz = 88-344 orthologous genes; Mz = 304-513 orthologous genes) and tissues (Forebrain = 1302 orthologous genes; Retina = 2747 orthologous genes; Liver = 1124 orthologous genes; Testis = 1562 orthologous genes).

By calculating the standard deviation (SD), mean, and coefficient of variation (CV) of both 'conserved' ancestral (set 1) and species-specific (set 2) gene promoter peaks in orthologous genes, we identified that all tissues show higher variability of orthologous genes in set 1 (forebrain CV=169%; retina CV=162%, liver CV=155%; testis CV=66%) compared to their counterparts in set 2 (forebrain CV=71%; retina CV=122%, liver CV=50%; testis CV=44%) (Fig. 2a). However, peaks in gene promoters of retina tissue exhibit high variability in both sets, indicating dynamic evolution of chromatin accessibility in retina-related genes both ancestrally and at a species-specific level. In forebrain and liver, there is high variability of orthologous genes at ancestral nodes, but low-moderate variability at the species level, suggesting more dynamic evolutionary processes in forebrain-related and liver-related genes at ancestral levels, but more conserved patterns of chromatin-accessibility at the species level. Testis on the other hand shows the lowest variability of orthologous genes amongst all tissues in ancestral nodes and relatively low variability at species-specific nodes, indicating more consistent evolutionary patterns of chromatin accessibility in testis-related genes across both ancestral and species-specific contexts.

This identified peaks in individual tissues of gene promoters potentially relevant to species-specific function; in forebrain tissue (71-513 orthologous genes), this includes *lrrtm1* in *A. burtoni* and *vldlr* in *P. nyererei* associated with modulating synaptic cell adhesion [6] and neural development [7] respectively; in retina tissue (36-1762 orthologous genes), this includes *fgf10* in *M. zebra* and *sws1* in *N. brichardi* that are associated with eye morphogenesis [8] and short wavelength sensitive visual function in cichlids [9] respectively; in liver tissue (73-380 orthologous genes), *cxcr3* in *M. zebra* and *cxcr1* in *O. niloticus,* genes associated with hepatic immune response [10,11]; and in testis tissue (88-439 orthologous genes), *plag1* in *P. nyererei* associated with spermatogenesis [12], and *prdm9* in *A. burtoni* associated with meiotic recombination [13].

**Accelerated divergence of active gene promoter sites drives cichlid regulatory diversity**

To study the divergence of open chromatin peaks, whether conserved or diverged across tissues and species, we calculated the rate of nucleotide substitutions of each *O. niloticus* peak as a reference against the four other species, compared to a neutral model using phyloP [14] (see '*Materials and Methods*'). Within each annotation, 61-85% of *O. niloticus* open chromatin peaks are conserved with the other four species, whereas 7-35% exhibit accelerated evolution (Supplementary Fig. S5-S6 and Supplementary Table S6). Consistent with our previous findings of regulatory divergence in cichlid gene promoter regions [4,15], the highest proportion (35%) of open chromatin peaks in *O. niloticus* tissues exhibiting accelerated evolution along the phylogeny, are found to overlap gene promoter regions (Supplementary Fig. S5-S6 and Supplementary Table S6). Notably, the proportion of accelerated peaks are significantly different (Wilcoxon rank sum test, (Benjamini-Hochberg adjusted *p-value* $<0.05$) between annotations, especially between comparisons of gene promoter regions with the other annotations (5' UTR, Exon, Intron and 3' UTR) both within *O. niloticus*, and pairwise between *O. niloticus* and the other species (Supplementary Table S7). Within the 13,889 orthologous genes, 12,922 have *O. niloticus* open chromatin peaks that are localised to gene promoter regions and of which, 7,615 (59%) exhibit accelerated evolution in at least one lineage along the phylogeny. A total of 5,536 unique orthologous genes (202-4321 orthologous genes in each tissue) have open chromatin peaks in orthologous gene promoter regions of all five species (Anc1, Fig. 1d) of which, 3,061 (55%) exhibit accelerated evolution along the phylogeny. Overall, this lends further support to focus on open chromatin peaks overlapping gene promoter regions of 1-to-1 orthologous genes to 1) characterise divergence of tissue-specific peaks, including gains or losses and genetic diversity along the phylogeny and 2) study any open chromatin and transcriptional divergence correlations.

To examine the divergence of gene promoter peaks in the phylogeny, gene promoter peak gain and loss for each lineage was quantified and categorised according to corresponding pairwise genetic diversity (Fig. 2e, Supplementary Table S8 and S9). Since there is no previous knowledge of expected divergence at active gene promoter sites in cichlid species, we use *O. niloticus* peaks as a reference where peak summits overlap in all pairwise comparisons (Supplementary Table S8, see *Materials and Methods*). In all comparisons against *O. niloticus*, *N. brichardi* has the most peak gains in retina (52% of all 13,889 1-to-1 orthologous gene promoter regions), liver (44%), and testis (48%) tissue. This is not biased by genome completeness or annotation quality, since *N. brichardi* has one of the fewest annotated protein-coding genes (23,568 – compared to 23,044-28,142 in the other four species [16-18]) and lowest genome contig N50 (13.2 kb – compared to 20kb - 3.1 Mb in the other four species [16-18]), whereas *O. niloticus*, used as the reference here for an unbiased approach, has the most complete genome (3.1 Mb) and highest protein coding gene count (28,142 protein-coding genes) [17].

In forebrain tissue, the most peak gains (45% of 1-to-1 orthologous genes) are instead observed in *M. zebra,* with an enrichment of tissue function, e.g., neurogenesis (Benjamini-Hochberg [19] adjusted *p-value* <0.05), with the fewest peak gains in general found in *A. burtoni,* e.g., 15% in forebrain (Supplementary Table S8), with unique functional enrichments, e.g., monatomic transmembrane transport (Benjamini-Hochberg [19] adjusted *p-value* <0.05), and possibly relevant to the *A. burtoni* model species for behaviour and social neuroscience [20]. Across all four tissues and pairwise comparisons, a total of 27-40% (*M. zebra*), 36-47% (*P. nyererei*), 19-38% (*A. burtoni*), and 30-47% (*N. brichardi*) of the 13,889 1-to-1 orthologous genes exhibit a loss of at least one gene promoter peak (Fig. 2e,

Supplementary Table S8). Overall, peak loss in all species could be attributed to sequence divergence as 76-91% of peak loss regions present more than 10% pairwise sequence divergence (Fig. 2e, Supplementary Table S8). Consistent with previous findings of regulatory site divergence [4,15], more than 10% sequence divergence over a mean peak size of 236 to 419 bp across all samples, could be indicative of the gain or loss of regulatory sites, e.g., TFBSs (mean size of 13 bp across all five species). We therefore conclude that increased nucleotide divergence at active gene promoter regions is associated with chromatin accessibility peak loss and could drive novel divergence of functionally active regulatory sites associated with tissue-specific traits e.g., the visual systems.

**Divergence of transcription factor activity underpins adaptive regulatory innovations**

To identify conserved and species- or clade-specific binding patterns, tissue-specific TF activity was clustered into co-active groups within each species (see *Materials and Methods*). We identified 13 (forebrain, n=1427 unique TF motifs; Supplementary Fig. S9), 10 (retina, n=1424 unique TF motifs; Fig. 3c), 8 (liver, n=1430 unique TF motifs; Supplementary Fig. S10), and 11 (testis, n=1427 unique TF motifs; Supplementary Fig. S11) clusters of binding patterns, with varying levels of orthologous TF binding activity observed across the five species. For example, in forebrain tissue, *prox1* associated with zebrafish neurogenesis [21], is highly activated in cluster 10 of *M. zebra* (Z-score of 1.7) but has low activity in the other species (Z-score of -0.8 to -0.2), whereas *otx2* associated with brain morphogenesis [22], is highly activated in cluster 1 of *A. burtoni* (Z-score of 1.8) but has low activity in the other species (Z-score of -0.6 to -0.3) (Supplementary Fig. S9). Varying activity is also observed in liver tissue, where *p73* associated with liver fuel metabolism [23] is highly activated in cluster 6 of *N. brichardi* (Z-score of 1.8 versus -0.7 to -0.2 in the other species), and *runx3* associated with liver development [24] is highly activated in cluster 3 of *O. niloticus* (Z-score of 1.6 versus -1 to 0.2 in the other species) (Supplementary Fig. S10). This varying activity is also observed in testis tissue, where *pbx1* associated with testis-determination [25] is highly active in cluster 10 of *P. nyererei* (Z-score of 1.8 versus -0.6 to -0.3 in the other species) (Supplementary Fig. S8).

**Dynamic regulatory site divergence drives gene regulatory network rewiring in cichlids**

Across the four tissues and species, 35-66% of pairwise polymorphic sites (in the promoter regions of 6842 – 8200 1-to-1 orthologous genes) map to *O. niloticus* TF motifs (Fig. 4b, Supplementary Table S14b) and thus, could impact TF binding activity. Of these, we define a 'principal set' where 1-5% polymorphic sites (1385-1439 non-redundant TF motifs in the promoter regions of 749-2747 genes) are in partially (< 100%) overlapping TF motifs between each species tissue and *O. niloticus,* where the majority (99.9%) of overlapping TF motifs are different (Fig. 4c, Supplementary Table S14c). Further, 6-13% of these polymorphic sites (175-1361 non-redundant TF motifs in the promoter regions of 18-618 1-to-1 orthologous genes) are in 'accelerated peaks' (Supplementary Table S14c), indicative of accelerated nucleotide divergence in actively bound TFs of the comparison species, relative to neutrally evolving sites in *O. niloticus*.

We observed concrete examples where motif-altering variants result in predicted switching of TF binding sites associated with genes linked to adaptive traits. In the accelerated set (of 24-769 genes), we identified examples of both discrete, e.g., one variant changing an IRF7 site to USF2 in the retinal gene, *pitpn* [26] (Fig. 4g), in *P. nyererei,* and indiscrete nucleotide variation, e.g., seven variants changing a STAT1 site to TEAD4 in the liver development gene [27], *foxa2*, in *N. brichardi.* Altogether, both subtle and significant genetic variations in motifs of actively bound TFs have likely contributed to motif divergence and GRN rewiring of several genes linked to adaptive traits across the five species.

**Chromatin accessibility divergence correlates with transcriptional regulation of genes linked to cichlid adaptive traits**

Upon investigating the correlation between accessible gene promoter regions, including divergent TF binding affinities, and transcriptional changes across species and tissues, we identified that in all five species retina tissue, peak-gene expression relationships were most abundant in the HA-HE category (37-41% of all categorised relationships), with 1408-4842 non-redundant peaks and 902-2020 non-redundant genes (Fig. 5a, Fig. S18-S22, and Supplementary Table S16). In the same tissue, the remainder are categorised as either HA-ME (929-3069 peaks and 588-1325 non-redundant genes: 23-24% of all relationships), MA-HE (1212-5862 peaks and 885-2487 non-redundant genes: 20-24% of all relationships), or MA-ME (790-4290 peaks and 542-1812 non-redundant genes: 15-16% of all relationships) (Fig. 5a, Fig. S18-S22, and Supplementary Table S16).

**Genetic variation in active TFBSs segregates with phylogeny and ecology in East African cichlids**

We hypothesise that genetic variation between the five cichlid species and their respective lake or riverine species counterparts can identify functional variation in TFBSs of orthologous genes associated with niche- and/or clade- specific adaptive traits that differ between the species. To prioritise 1-to-1 orthologous genes, we ranked TFBSs in promoter regions using the best described classification method [28], by selecting TFBSs with tag counts (TC, number of reads) and bit-score (BS) of motif match above the means for all statistically significant predicted TFBSs in each species tissue (Benjamini-Hochberg [19] adjusted *p-value* <0.0001) (see '*Materials and Methods*'). We started with 896,634-2,247,267 positionally non-redundant TFBSs in 93,139-226,058 footprints, characterised in the promoter regions of 13,774-21,553 genes of the five species retina tissues. In this set, we identified 108,193-354,906 (10-20%) predicted TFBSs in 18726-59883 (15-32%) footprints in the promoter regions of 6844-13886 (41-76%) genes above the mean TC and BS of all genes (Fig. 7a; Supplementary Table S21). This is comparable to that identified in *O. niloticus* gill tissue previously [29] and amongst these, we identified candidate transcription factor (TF) – target gene (TG) relationships where dynamic divergence of regulatory sites correlates with transcriptional divergence, concentrating on well-defined cichlid visual system genes that also strongly associate with niche diversity [30]. We then filtered for 13,997 1-to-1 orthologous genes, and then categorised TF-TG candidates where 1) the TG is HA-HE and a TFBS in the promoter has variation between species e.g., *O. niloticus* RUNX1-*rho*; 2) a TFBS has variation between species e.g., *P. nyererei* USF2-*pitpn*; or 3) the TG is HA-HE e.g., *N. brichardi* NR2F6-*sws1* (Fig. 7a).

We examined the photoreceptor gene, *actr1*, under divergent selection in Lake Malawi cichlids from the twilight zone [31], and predicted to be regulated by MAF, a transcription factor involved in lens development [32]. The active MAF binding site is conserved in the orthologous *actr1* gene promoter of *O. niloticus* (Fig. 7b) and *M. zebra* (Fig. 7c) with segregating variants in the last (tilapia species, Fig. 7d) and first (a Lake Malawi species*,* Fig. 7e, Supplementary Fig. S38) positions of the binding site (Fig. 7f-g). The homozygous variant (T/T) in the last position is 1) conserved in the same clade species, *O. spilurus* and *O. aureus*; but 2) heterozygous or homozygous segregating in all 21 distantly related tilapia species with diverse salinity and habitat adaptations, for example, *O. urolepis* and *O. mossambicus* (both C/C) (Fig. 7d). Conversely, the homozygous variant (T/T) in the first position is 1) conserved with all 66 other Lake Malawi species; but 2) homozygous segregating in the deep benthic, *Atlicorpus macrocleithrum* (A/A, Fig. 7e and 6g, Supplementary Fig. S39), that occupies a niche utilising enlarged sensory pores on the lower head and a uniquely adapted elongated cleithrum on the chest [33]. This variant was confirmed by identifying the 1) *A. macrocleithrum* segregating variant in multiple reads [34,35] (Fig. 7g, Supplementary Fig. S39); and 2) corresponding *actr1* gene promoter and MAF TFBS region in chromosome-level genomes of representative Lake Malawi caldes (*A. calliptera, Tropheops* sp. 'mauve'*, Aulonocara stuartgranti,* and *Ramphochromis* sp. 'Chilingali') genomes [35] (Supplementary Fig. S39).
Both MAF TFBS variants are predicted to be disruptive, where MotifDiff scores of less than 0, indicate a significant decrease of binding probability [36] of MAF to the *actr1* gene promoter of 1) 21 tilapia species with diverse salinity and habitat adaptations (Fig. 7f, *MotifDiff* score of -3.8); and 2) the Lake Malawi deep benthic species, *A. macrocleithrum* (Fig. 7f, *MotifDiff* score of -2.5). Overall, this suggests

that MAF 1) could be a core regulator of *actr1* in the closely related clade of *O. niloticus*, *O. spilurus* and *O. aureus* (Fig. 7d), as well as most Lake Malawi species (Fig. 7e) where photoreception in diverse habitats, influencing feeding, social, and reproductive behaviour is vital for overall fitness; but 2) is unlikely to regulate *actr1* in the rest of the tilapia phylogeny, as well as the uniquely adapted LM deep benthic species, *A. macrocleithrum* (Fig. 7), possibly indicating niche-specific divergence of photoreceptor regulation across cichlid phylogenies. Further, phylogenetic independent contrast (PIC) analysis [37] of the *actr1* (Supplementary Fig. S40) genotype against ecology of each of the 73 Lake Malawi species, highlights very little change in correlation once the phylogeny is taken into account and a regression model fitted (*see Materials and Methods*).

In summary, we identified active TF binding sites of visual system genes that segregate according to phylogeny and ecology of radiating and non-radiating cichlid fish species. The presence of distinct variants at two sites within a positionally conserved TFBS, segregating independently across both a non-radiating tilapia lineage and the rapidly radiating Lake Malawi cichlids, suggests that independent selective pressures may be acting on different positions within conserved regulatory elements in each lineage - indicative of lineage-specific adaptation even at conserved regulatory motifs. This highlights the role of regulatory variation TFBSs as a key contributor to adaptive innovations across East African cichlids.
