## Supplementary Figures for "Evolution of chromatin accessibility associated with traits of cichlid phenotypic diversity"

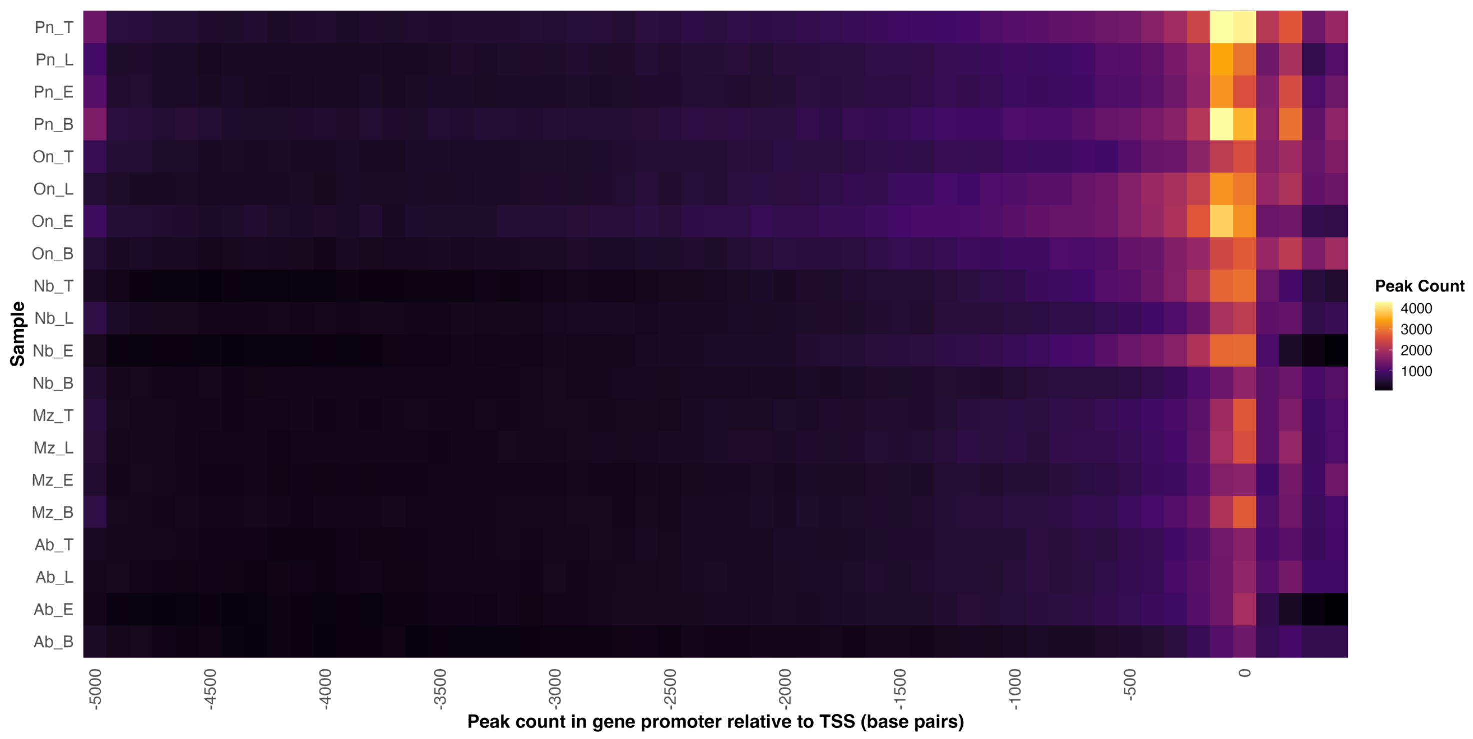


**Fig. S1 – Distribution of narrow open-chromatin peaks relative to TSS.** Heatmap showing the distribution of peak summits relative to the transcription start site (TSS) for sample replicates. Peak counts were calculated using a 100-bp sliding window from the TSS (at 0 bp) along the 5 kb gene promoter region. The colour scale indicates the number of peaks within each window. Species: Mz = *M. zebra*; Pn = *P. nyererei*; Ab = *A. burtoni*; Nb = *N. brichardi*; On = *O. niloticus*; Tissues: B = Forebrain; E = Retina; L = Liver; T = Testis.

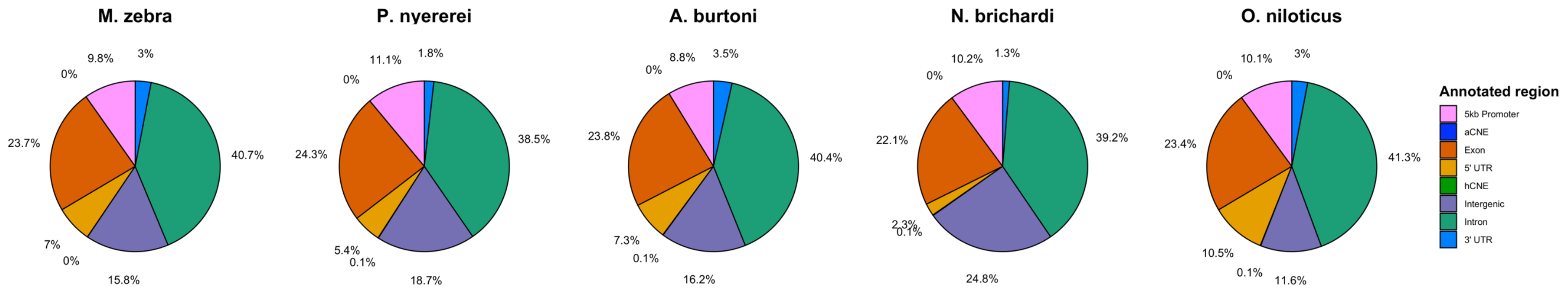


**Fig. S2 – Distribution of narrow open-chromatin peaks in annotated features of four tissues in five cichlids.** Collated percentage of narrow open-chromatin peaks in all four tissue samples of five cichlids shown outside of each annotated feature with legend of annotations to right.

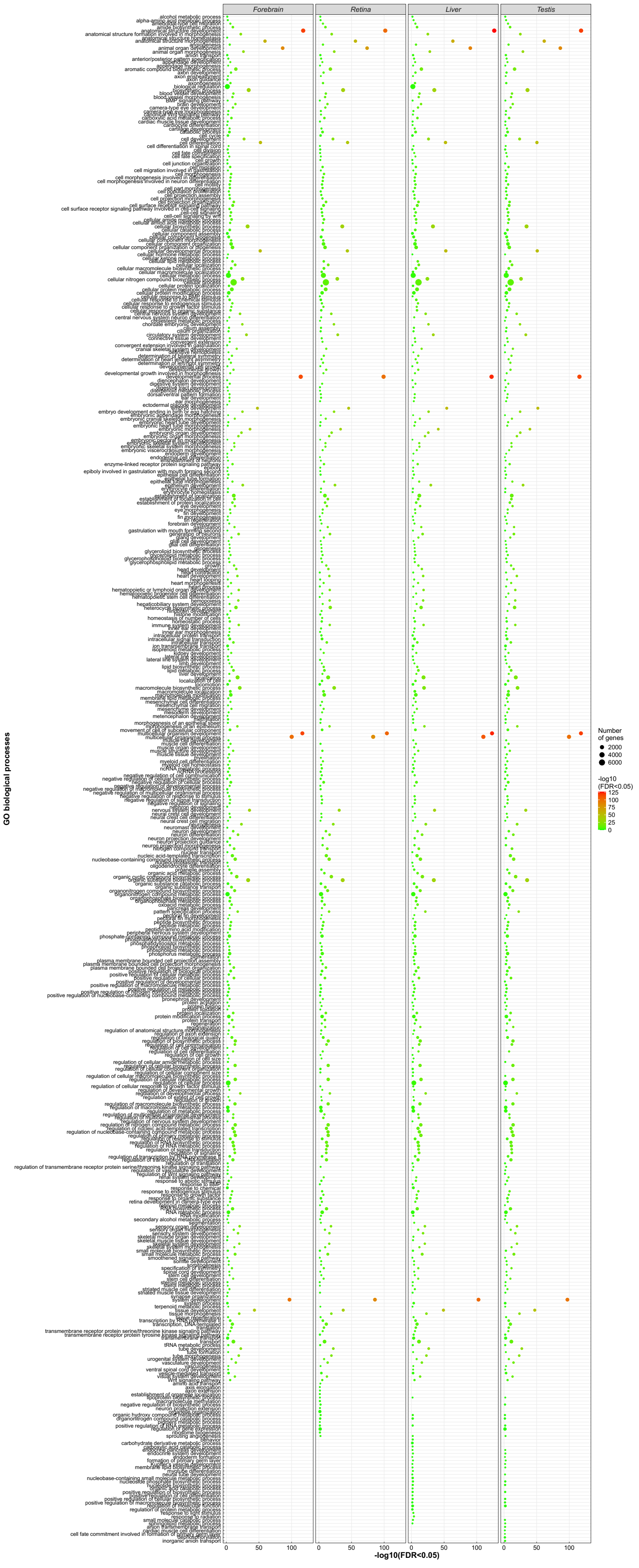


**Fig. S3 – Gene Ontology (GO) enrichment of peaks localised to 5 kb gene promoters of 1-to-1 orthologs collated by tissue.** GO biological processes enrichment of peaks localised to the up to 5 kb gene promoter regions of 13,889 1-to-1 orthologs collated for all five species by tissue. Circles show enriched biological processes (y-axis) of significance (-$log_{10}$ FDR <0.05, heatmap to right) and -$log_{10}$ FDR <0.05 (x-axis) values of each term across each tissue. Number of enriched genes for each term shown by size of each circle.

**(A)**

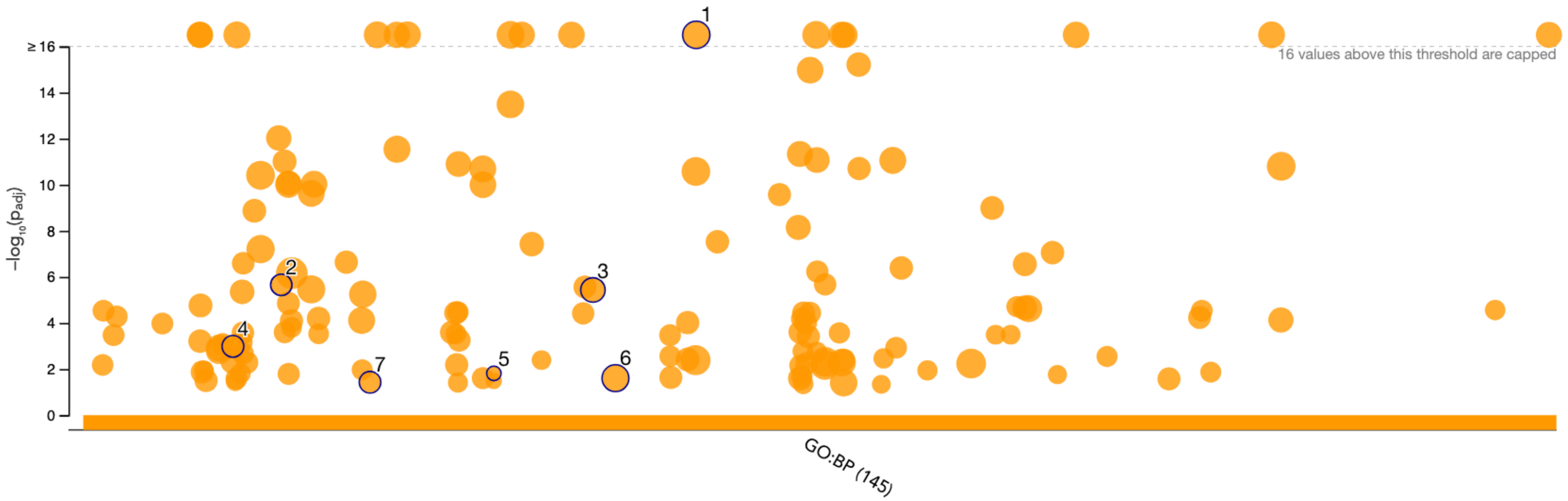


| ID | Source | Term ID | Term Name | $p_{adj}$ (query_1) |
|---|---|---|---|---|
| 1 | GO:BP | GO:0044271 | cellular nitrogen compound biosynthetic process | $8.034\times10^{-28}$ |
| 2 | GO:BP | GO:0009719 | response to endogenous stimulus | $2.214\times10^{-6}$ |
| 3 | GO:BP | GO:0035556 | intracellular signal transduction | $3.706\times10^{-6}$ |
| 4 | GO:BP | GO:0007167 | enzyme-linked receptor protein signaling path... | $1.027\times10^{-3}$ |
| 5 | GO:BP | GO:0032007 | negative regulation of TOR signaling | $1.541\times10^{-2}$ |
| 6 | GO:BP | GO:0036211 | protein modification process | $2.482\times10^{-2}$ |
| 7 | GO:BP | GO:0016477 | cell migration | $3.737\times10^{-2}$ |

**(B)**

| GO:BP | | stats | |
|---|---|---|---|
| Term name | Term ID | $p_{adj}$ | $-\log_{10}(p_{adj})$ 0 ≤16 |
| developmental process | GO:0032502 | $2.812\times10^{-17}$ | |
| anatomical structure development | GO:0048856 | $4.821\times10^{-17}$ | |
| multicellular organism development | GO:0007275 | $3.826\times10^{-16}$ | |
| system development | GO:0048731 | $1.410\times10^{-13}$ | |
| cell differentiation | GO:0030154 | $6.645\times10^{-10}$ | |
| cellular developmental process | GO:0048869 | $7.070\times10^{-10}$ | |
| multicellular organismal process | GO:0032501 | $5.077\times10^{-9}$ | |
| anatomical structure morphogenesis | GO:0009653 | $8.017\times10^{-9}$ | |
| nervous system development | GO:0007399 | $1.771\times10^{-8}$ | |
| neurogenesis | GO:0022008 | $6.599\times10^{-8}$ | |
| animal organ development | GO:0048513 | $1.504\times10^{-7}$ | |
| generation of neurons | GO:0048699 | $6.864\times10^{-7}$ | |
| tissue development | GO:0009888 | $1.088\times10^{-6}$ | |
| neuron differentiation | GO:0030182 | $1.311\times10^{-6}$ | |
| epithelium development | GO:0060429 | $2.479\times10^{-5}$ | |
| embryo development | GO:0009790 | $1.067\times10^{-4}$ | |
| cell development | GO:0048468 | $3.320\times10^{-4}$ | |
| embryonic morphogenesis | GO:0048598 | $5.015\times10^{-4}$ | |
| central nervous system development | GO:0007417 | $6.624\times10^{-3}$ | |
| animal organ morphogenesis | GO:0009887 | $7.899\times10^{-3}$ | |
| sensory system development | GO:0048880 | $1.122\times10^{-2}$ | |
| regulation of cell differentiation | GO:0045595 | $1.396\times10^{-2}$ | |
| plasma membrane bounded cell projection morphogene... | GO:0120039 | $1.500\times10^{-2}$ | |
| neuron projection morphogenesis | GO:0048812 | $1.500\times10^{-2}$ | |
| cell projection morphogenesis | GO:0048858 | $1.500\times10^{-2}$ | |
| cell surface receptor protein tyrosine kinase signaling p... | GO:0007169 | $1.733\times10^{-2}$ | |
| positive regulation of cellular process | GO:0048522 | $2.126\times10^{-2}$ | |
| regulation of RNA metabolic process | GO:0051252 | $2.768\times10^{-2}$ | |
| regulation of developmental process | GO:0050793 | $2.856\times10^{-2}$ | |
| neuron projection development | GO:0031175 | $2.967\times10^{-2}$ | |
| regulation of nucleobase-containing compound metabo... | GO:0019219 | $3.233\times10^{-2}$ | |
| regulation of RNA biosynthetic process | GO:2001141 | $3.874\times10^{-2}$ | |
| regulation of DNA-templated transcription | GO:0006355 | $3.874\times10^{-2}$ | |
| neuron development | GO:0048666 | $3.961\times10^{-2}$ | |
| cell morphogenesis | GO:0000902 | $4.032\times10^{-2}$ | |

**Fig. S4 – Gene Ontology (GO) enrichment of peaks localised to ancestral nodes.** GO enrichment of all tissue peaks at **A)** 5536 1-to-1 ortholog gene promoters positionally conserved across all five species (Anc1); and **B)** 1473 1-to-1 ortholog gene promoters positionally conserved across haplochromine species. Circles and bars show enriched biological processes (x-axis and table) of significance ($-log_{10}$ FDR <0.05, y-axis) of each term in Supplementary Table S4. In (A) table of displayed driver terms (demarcated with numbers on plot) calculated by the g:GOSt module of g:Profiler (https://biit.cs.ut.ee/gprofiler/gost) based on a two-stage algorithm for filtering GO enrichment results by grouping into sub-ontologies (stage 1) and identifying connected components using a greedy search strategy (stage 2).

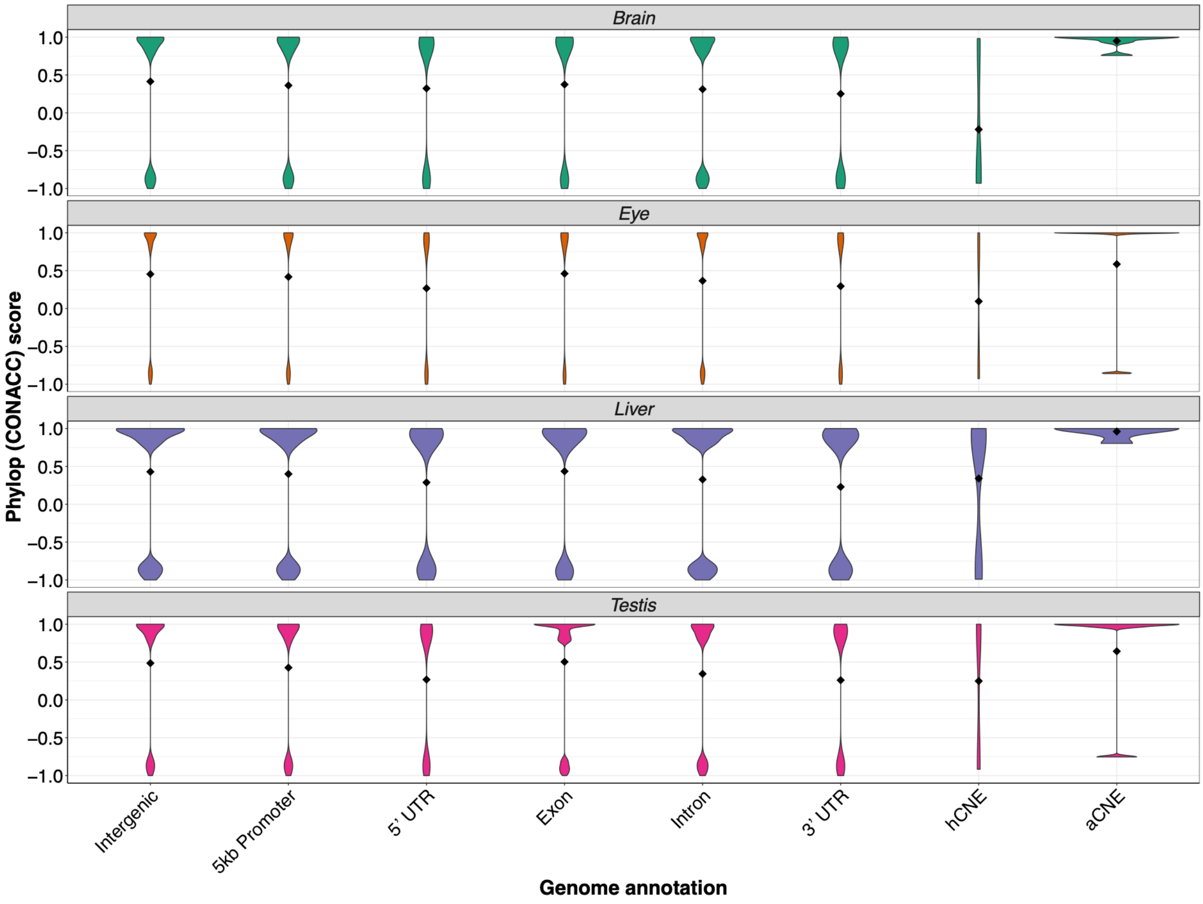


**Fig. S5 – PhyloP conservation-acceleration scores of *O. niloticus* open-chromatin peaks collated by tissues and genomic annotations.** Violin plot of collated pairwise phyloP conservation-acceleration (CONACC) scores (y-axis) of *O. niloticus* open-chromatin peaks in genomic annotations (x-axis) of four tissues (Brain = Forebrain and Eye = Retina). PhyloP scores were calculated using *O. niloticus* as a reference against all species pairwise comparisons and collated; positive scores indicate pairwise evolutionary conservation, whereas negative scores indicate evolutionary acceleration against a neutral substitution model.

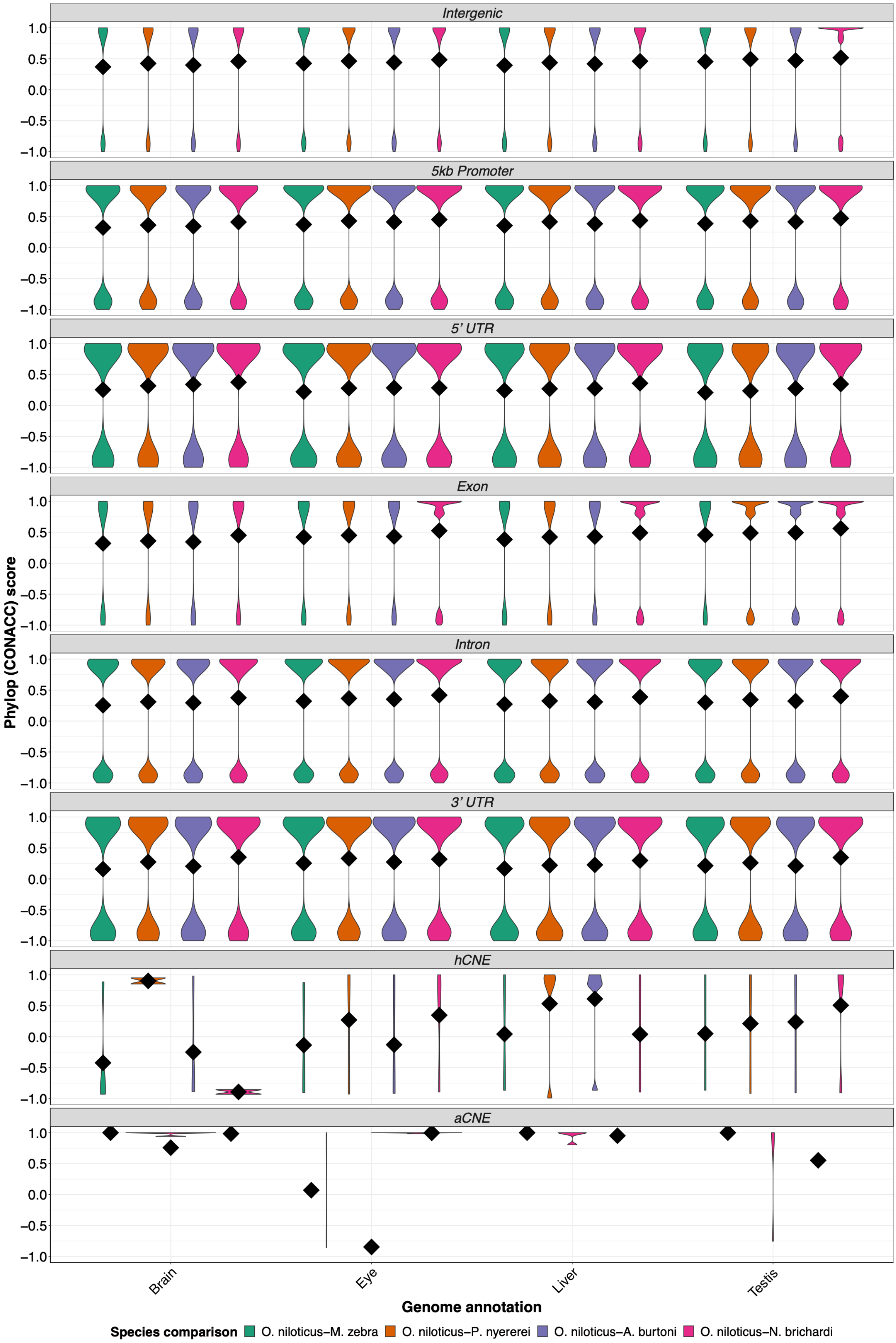

Intergenic
5kb Promoter
5' UTR
Exon
Intron
3' UTR
hCNE
aCNE
Phylop (CONACC) score
1.0
0.5
0.0
−0.5
−1.0
Brain
Eye
Liver
Testis
Genome annotation
Species comparison
O. niloticus−M. zebra
O. niloticus−P. nyererei
O. niloticus−A. burtoni
O. niloticus−N. brichardi

**Fig. S6 – PhyloP conservation-acceleration scores of *O. niloticus* open-chromatin peaks in pairwise species comparisons of tissues and genomic annotations.** Violin plot of all pairwise phyloP conservation-acceleration (CONACC) scores (y-axis) of *O. niloticus* open-chromatin peaks in genomic annotations of four tissues (x-axis). PhyloP scores were calculated using *O. niloticus* as a reference against all species pairwise comparisons and individually plotted; positive scores indicate pairwise evolutionary conservation, whereas negative scores indicate evolutionary acceleration against a neutral substitution model.

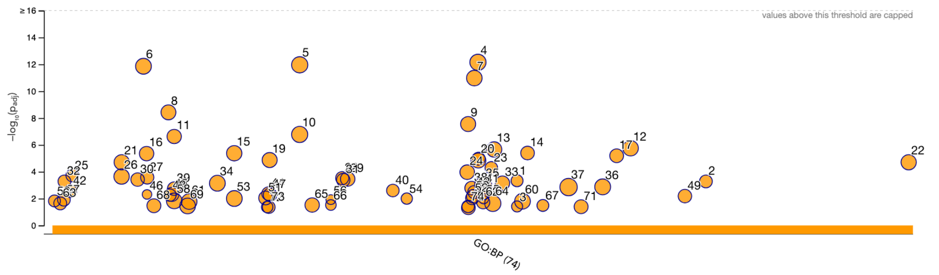


| ID | Source | Term ID | Term Name | $p_{adj}$ (query_1) |
|---|---|---|---|---|
| 1 | GO:BP | GO:0060041 | retina development in camera-type eye | $4.703\times10^{-4}$ |
| 2 | GO:BP | GO:0150063 | visual system development | $5.246\times10^{-4}$ |
| 3 | GO:BP | GO:0060042 | retina morphogenesis in camera-type eye | $3.835\times10^{-2}$ |
| 4 | GO:BP | GO:0048856 | anatomical structure development | $7.112\times10^{-13}$ |
| 5 | GO:BP | GO:0032502 | developmental process | $1.093\times10^{-12}$ |
| 6 | GO:BP | GO:0007275 | multicellular organism development | $1.401\times10^{-12}$ |
| 7 | GO:BP | GO:0048731 | system development | $1.065\times10^{-11}$ |
| 8 | GO:BP | GO:0009653 | anatomical structure morphogenesis | $3.800\times10^{-9}$ |
| 9 | GO:BP | GO:0048513 | animal organ development | $2.817\times10^{-8}$ |
| 10 | GO:BP | GO:0032501 | multicellular organismal process | $1.678\times10^{-7}$ |
| 11 | GO:BP | GO:0009888 | tissue development | $2.376\times10^{-7}$ |
| 12 | GO:BP | GO:0080090 | regulation of primary metabolic process | $1.771\times10^{-6}$ |
| 13 | GO:BP | GO:0051252 | regulation of RNA metabolic process | $2.193\times10^{-6}$ |
| 14 | GO:BP | GO:0060429 | epithelium development | $4.052\times10^{-6}$ |
| 15 | GO:BP | GO:0019219 | regulation of nucleobase-containing compoun… | $4.216\times10^{-6}$ |
| 16 | GO:BP | GO:0007399 | nervous system development | $4.479\times10^{-6}$ |
| 17 | GO:BP | GO:0072359 | circulatory system development | $6.457\times10^{-6}$ |
| 18 | GO:BP | GO:0048880 | sensory system development | $1.072\times10^{-5}$ |
| 19 | GO:BP | GO:0030154 | cell differentiation | $1.348\times10^{-5}$ |
| 20 | GO:BP | GO:0048869 | cellular developmental process | $1.462\times10^{-5}$ |
| 21 | GO:BP | GO:0006355 | regulation of DNA-templated transcription | $1.977\times10^{-5}$ |
| 22 | GO:BP | GO:2001141 | regulation of RNA biosynthetic process | $1.977\times10^{-5}$ |
| 23 | GO:BP | GO:0051128 | regulation of cellular component organization | $5.196\times10^{-5}$ |
| 24 | GO:BP | GO:0048468 | cell development | $1.083\times10^{-4}$ |
| 25 | GO:BP | GO:0002009 | morphogenesis of an epithelium | $1.893\times10^{-4}$ |
| 26 | GO:BP | GO:0006351 | DNA-templated transcription | $2.271\times10^{-4}$ |
| 27 | GO:BP | GO:0007423 | sensory organ development | $2.650\times10^{-4}$ |
| 28 | GO:BP | GO:0035239 | tube morphogenesis | $2.979\times10^{-4}$ |
| 29 | GO:BP | GO:0035556 | intracellular signal transduction | $3.478\times10^{-4}$ |
| 30 | GO:BP | GO:0007010 | cytoskeleton organization | $3.843\times10^{-4}$ |
| 31 | GO:BP | GO:0035295 | tube development | $4.142\times10^{-4}$ |
| 32 | GO:BP | GO:0001654 | eye development | $5.246\times10^{-4}$ |
| 33 | GO:BP | GO:0051641 | cellular localization | $7.085\times10^{-4}$ |
| 34 | GO:BP | GO:0016043 | cellular component organization | $7.189\times10^{-4}$ |
| 35 | GO:BP | GO:0050794 | regulation of cellular process | $1.042\times10^{-3}$ |
| 36 | GO:BP | GO:0071840 | cellular component organization or biogenesis | $1.341\times10^{-3}$ |
| 37 | GO:BP | GO:0065007 | biological regulation | $1.390\times10^{-3}$ |
| 38 | GO:BP | GO:0048646 | anatomical structure formation involved in mor… | $1.664\times10^{-3}$ |
| 39 | GO:BP | GO:0009887 | animal organ morphogenesis | $1.814\times10^{-3}$ |
| 40 | GO:BP | GO:0043010 | camera-type eye development | $2.515\times10^{-3}$ |
| 41 | GO:BP | GO:0050789 | regulation of biological process | $2.746\times10^{-3}$ |
| 42 | GO:BP | GO:0001944 | vasculature development | $2.765\times10^{-3}$ |
| 43 | GO:BP | GO:0048729 | tissue morphogenesis | $3.295\times10^{-3}$ |
| 44 | GO:BP | GO:0030030 | cell projection organization | $4.256\times10^{-3}$ |
| 45 | GO:BP | GO:0009790 | embryo development | $4.862\times10^{-3}$ |
| 46 | GO:BP | GO:0007416 | synapse assembly | $4.913\times10^{-3}$ |
| 47 | GO:BP | GO:0030182 | neuron differentiation | $5.245\times10^{-3}$ |
| 48 | GO:BP | GO:0009719 | response to endogenous stimulus | $5.497\times10^{-3}$ |
| 49 | GO:BP | GO:0120036 | plasma membrane bounded cell projection org… | $6.547\times10^{-3}$ |
| 50 | GO:BP | GO:0048699 | generation of neurons | $6.740\times10^{-3}$ |
| 51 | GO:BP | GO:0022008 | neurogenesis | $9.003\times10^{-3}$ |
| 52 | GO:BP | GO:0048666 | neuron development | $9.270\times10^{-3}$ |
| 53 | GO:BP | GO:0019222 | regulation of metabolic process | $9.861\times10^{-3}$ |
| 54 | GO:BP | GO:0044087 | regulation of cellular component biogenesis | $1.001\times10^{-2}$ |
| 55 | GO:BP | GO:0050808 | synapse organization | $1.023\times10^{-2}$ |
| 56 | GO:BP | GO:0034329 | cell junction assembly | $1.202\times10^{-2}$ |
| 57 | GO:BP | GO:0001568 | blood vessel development | $1.228\times10^{-2}$ |
| 58 | GO:BP | GO:0009889 | regulation of biosynthetic process | $1.494\times10^{-2}$ |
| 59 | GO:BP | GO:0000226 | microtubule cytoskeleton organization | $1.498\times10^{-2}$ |
| 60 | GO:BP | GO:0060255 | regulation of macromolecule metabolic process | $1.570\times10^{-2}$ |
| 61 | GO:BP | GO:0010556 | regulation of macromolecule biosynthetic proc… | $1.666\times10^{-2}$ |
| 62 | GO:BP | GO:0050793 | regulation of developmental process | $1.872\times10^{-2}$ |
| 63 | GO:BP | GO:0000902 | cell morphogenesis | $2.279\times10^{-2}$ |
| 64 | GO:BP | GO:0051179 | localization | $2.285\times10^{-2}$ |
| 65 | GO:BP | GO:0033036 | macromolecule localization | $3.007\times10^{-2}$ |
| 66 | GO:BP | GO:0034330 | cell junction organization | $3.016\times10^{-2}$ |
| 67 | GO:BP | GO:0060972 | left/right pattern formation | $3.250\times10^{-2}$ |
| 68 | GO:BP | GO:0008104 | protein localization | $3.393\times10^{-2}$ |
| 69 | GO:BP | GO:0010468 | regulation of gene expression | $3.550\times10^{-2}$ |
| 70 | GO:BP | GO:0048514 | blood vessel morphogenesis | $3.850\times10^{-2}$ |
| 71 | GO:BP | GO:0070727 | cellular macromolecule localization | $4.030\times10^{-2}$ |
| 72 | GO:BP | GO:0022603 | regulation of anatomical structure morphogene… | $4.246\times10^{-2}$ |
| 73 | GO:BP | GO:0030036 | actin cytoskeleton organization | $4.338\times10^{-2}$ |
| 74 | GO:BP | GO:0048522 | positive regulation of cellular process | $4.792\times10^{-2}$ |

**Fig. S7 – Gene Ontology (GO) enrichment of positionally conserved and accelerated peaks localised to 3061 1-to-1 orthologous gene promoters along the five species phylogeny.** Circles show enriched biological processes (x-axis and table) of significance (-*log₁₀* FDR <0.05, y-axis) of each Nile tilapia term. Table of terms (demarcated with numbers on plot) calculated by the g:GOSt module of g:Profiler (https://biit.cs.ut.ee/gprofiler/gost).

### (A) *M. zebra* Forebrain

| GO:BP | | stats | |
|---|---|---|---|
| Term name | Term ID | $p_{adj}$ | $-\log_{10}(p_{adj})$ 0 … ≤16 |
| developmental process | GO:0032502 | $7.121\times10^{-55}$ | |
| anatomical structure development | GO:0048856 | $4.322\times10^{-53}$ | |
| multicellular organism development | GO:0007275 | $8.851\times10^{-49}$ | |
| system development | GO:0048731 | $6.015\times10^{-40}$ | |
| multicellular organismal process | GO:0032501 | $4.352\times10^{-31}$ | |
| cellular developmental process | GO:0048869 | $5.781\times10^{-30}$ | |
| cell differentiation | GO:0030154 | $9.826\times10^{-30}$ | |
| animal organ development | GO:0048513 | $1.515\times10^{-27}$ | |
| anatomical structure morphogenesis | GO:0009653 | $3.855\times10^{-26}$ | |
| nervous system development | GO:0007399 | $1.000\times10^{-24}$ | |
| tissue development | GO:0009888 | $2.031\times10^{-20}$ | |
| transport | GO:0006810 | $4.073\times10^{-18}$ | |
| neurogenesis | GO:0022008 | $1.968\times10^{-16}$ | |
| localization | GO:0051179 | $4.528\times10^{-16}$ | |
| generation of neurons | GO:0048699 | $6.116\times10^{-16}$ | |
| establishment of localization | GO:0051234 | $6.299\times10^{-16}$ | |
| regulation of cellular process | GO:0050794 | $8.868\times10^{-16}$ | |
| biological regulation | GO:0065007 | $1.606\times10^{-15}$ | |
| regulation of biological process | GO:0050789 | $3.912\times10^{-15}$ | |
| embryo development | GO:0009790 | $2.899\times10^{-14}$ | |
| neuron differentiation | GO:0030182 | $5.190\times10^{-14}$ | |
| transmembrane transport | GO:0055085 | $1.171\times10^{-13}$ | |
| central nervous system development | GO:0007417 | $3.961\times10^{-13}$ | |
| cell development | GO:0048468 | $2.238\times10^{-12}$ | |
| epithelium development | GO:0060429 | $5.389\times10^{-12}$ | |
| regulation of primary metabolic process | GO:0080090 | $1.201\times10^{-11}$ | |
| DNA-templated transcription | GO:0006351 | $1.387\times10^{-11}$ | |
| regulation of RNA biosynthetic process | GO:2001141 | $1.782\times10^{-11}$ | |
| regulation of DNA-templated transcription | GO:0006355 | $1.782\times10^{-11}$ | |
| regulation of RNA metabolic process | GO:0051252 | $2.678\times10^{-11}$ | |
| regulation of nucleobase-containing compound metabo... | GO:0019219 | $4.064\times10^{-11}$ | |
| anatomical structure formation involved in morphogene... | GO:0048646 | $5.329\times10^{-10}$ | |
| regulation of developmental process | GO:0050793 | $5.503\times10^{-10}$ | |
| embryonic morphogenesis | GO:0048598 | $2.075\times10^{-9}$ | |
| animal organ morphogenesis | GO:0009887 | $1.582\times10^{-8}$ | |
| monoatomic ion transmembrane transport | GO:0034220 | $1.681\times10^{-8}$ | |
| monoatomic ion transport | GO:0006811 | $1.921\times10^{-8}$ | |
| sensory organ development | GO:0007423 | $2.333\times10^{-8}$ | |
| sensory system development | GO:0048880 | $2.884\times10^{-8}$ | |
| cell communication | GO:0007154 | $8.026\times10^{-8}$ | |
| signaling | GO:0023052 | $2.253\times10^{-7}$ | |
| gland development | GO:0048732 | $2.751\times10^{-7}$ | |
| tube development | GO:0035295 | $3.638\times10^{-7}$ | |
| enzyme-linked receptor protein signaling pathway | GO:0007167 | $3.993\times10^{-7}$ | |
| cellular process | GO:0009987 | $6.732\times10^{-7}$ | |
| cell fate commitment | GO:0045165 | $9.771\times10^{-7}$ | |
| neuron development | GO:0048666 | $9.879\times10^{-7}$ | |
| cell morphogenesis | GO:0000902 | $1.626\times10^{-6}$ | |
| pattern specification process | GO:0007389 | $1.647\times10^{-6}$ | |
| axon development | GO:0061564 | $1.734\times10^{-6}$ | |
| regulation of cell communication | GO:0010646 | $2.364\times10^{-6}$ | |
| regulation of signaling | GO:0023051 | $2.451\times10^{-6}$ | |
| signal transduction | GO:0007165 | $3.887\times10^{-6}$ | |
| cell projection morphogenesis | GO:0048858 | $4.090\times10^{-6}$ | |
| neuron projection morphogenesis | GO:0048812 | $4.090\times10^{-6}$ | |
| plasma membrane bounded cell projection morphogene... | GO:0120039 | $4.090\times10^{-6}$ | |
| inorganic ion transmembrane transport | GO:0098660 | $4.263\times10^{-6}$ | |
| head development | GO:0060322 | $5.364\times10^{-6}$ | |
| tissue morphogenesis | GO:0048729 | $5.409\times10^{-6}$ | |
| brain development | GO:0007420 | $7.106\times10^{-6}$ | |
| cell morphogenesis involved in neuron differentiation | GO:0048667 | $7.266\times10^{-6}$ | |
| regionalization | GO:0003002 | $9.512\times10^{-6}$ | |
| axonogenesis | GO:0007409 | $9.729\times10^{-6}$ | |
| chordate embryonic development | GO:0043009 | $1.244\times10^{-5}$ | |
| embryo development ending in birth or egg hatching | GO:0009792 | $1.244\times10^{-5}$ | |
| regulation of signal transduction | GO:0009966 | $1.294\times10^{-5}$ | |
| circulatory system development | GO:0072359 | $2.165\times10^{-5}$ | |
| cell surface receptor protein tyrosine kinase signaling p... | GO:0007169 | $2.257\times10^{-5}$ | |
| response to endogenous stimulus | GO:0009719 | $3.547\times10^{-5}$ | |
| gastrulation | GO:0007369 | $4.102\times10^{-5}$ | |
| neuron projection guidance | GO:0097485 | $4.206\times10^{-5}$ | |
| axon guidance | GO:0007411 | $4.206\times10^{-5}$ | |
| vasculature development | GO:0001944 | $4.943\times10^{-5}$ | |
| cellular response to endogenous stimulus | GO:0071495 | $4.943\times10^{-5}$ | |
| metal ion transport | GO:0030001 | $5.691\times10^{-5}$ | |
| neuron projection development | GO:0031175 | $5.953\times10^{-5}$ | |
| hepaticobiliary system development | GO:0061008 | $6.270\times10^{-5}$ | |
| liver development | GO:0001889 | $8.259\times10^{-5}$ | |
| cell fate specification | GO:0001708 | $9.528\times10^{-5}$ | |
| morphogenesis of an epithelium | GO:0002009 | $9.591\times10^{-5}$ | |
| eye development | GO:0001654 | $1.526\times10^{-4}$ | |
| visual system development | GO:0150063 | $1.526\times10^{-4}$ | |
| plasma membrane bounded cell projection organization | GO:0120036 | $1.546\times10^{-4}$ | |
| regulation of cell differentiation | GO:0045595 | $1.688\times10^{-4}$ | |
| tube morphogenesis | GO:0035239 | $2.168\times10^{-4}$ | |
| blood vessel development | GO:0001568 | $2.932\times10^{-4}$ | |
| negative regulation of developmental process | GO:0051093 | $3.138\times10^{-4}$ | |
| regulation of transcription by RNA polymerase II | GO:0006357 | $3.159\times10^{-4}$ | |
| regulation of biological quality | GO:0065008 | $3.722\times10^{-4}$ | |
| regulation of multicellular organismal development | GO:2000026 | $4.047\times10^{-4}$ | |
| transcription by RNA polymerase II | GO:0006366 | $4.728\times10^{-4}$ | |
| sensory organ morphogenesis | GO:0090596 | $4.939\times10^{-4}$ | |
| positive regulation of cellular process | GO:0048522 | $5.625\times10^{-4}$ | |
| somitogenesis | GO:0001756 | $6.691\times10^{-4}$ | |
| cell projection organization | GO:0030030 | $8.504\times10^{-4}$ | |
| monoatomic anion transport | GO:0006820 | $8.953\times10^{-4}$ | |
| synaptic signaling | GO:0099536 | $9.334\times10^{-4}$ | |
| segmentation | GO:0035282 | $1.052\times10^{-3}$ | |
| inorganic cation transmembrane transport | GO:0098662 | $1.284\times10^{-3}$ | |
| cell-cell signaling | GO:0007267 | $1.583\times10^{-3}$ | |

### (B) *P. nyererei* Forebrain

| GO:BP | | stats | |
|---|---|---|---|
| Term name | Term ID | $p_{adj}$ | $-\log_{10}(p_{adj})$ 0 … ≤16 |
| anatomical structure development | GO:0048856 | $6.677\times10^{-43}$ | |
| developmental process | GO:0032502 | $6.287\times10^{-42}$ | |
| multicellular organism development | GO:0007275 | $5.180\times10^{-39}$ | |
| system development | GO:0048731 | $1.020\times10^{-32}$ | |
| animal organ development | GO:0048513 | $3.289\times10^{-22}$ | |
| anatomical structure morphogenesis | GO:0009653 | $8.222\times10^{-22}$ | |
| cellular developmental process | GO:0048869 | $8.162\times10^{-21}$ | |
| cell differentiation | GO:0030154 | $1.339\times10^{-20}$ | |
| multicellular organismal process | GO:0032501 | $1.221\times10^{-18}$ | |
| nervous system development | GO:0007399 | $2.468\times10^{-16}$ | |
| transport | GO:0006810 | $1.904\times10^{-14}$ | |
| tissue development | GO:0009888 | $4.050\times10^{-14}$ | |
| neuron differentiation | GO:0030182 | $7.455\times10^{-14}$ | |
| generation of neurons | GO:0048699 | $1.112\times10^{-13}$ | |
| localization | GO:0051179 | $7.386\times10^{-13}$ | |
| cell development | GO:0048468 | $1.119\times10^{-12}$ | |
| establishment of localization | GO:0051234 | $2.294\times10^{-12}$ | |
| neurogenesis | GO:0022008 | $4.383\times10^{-12}$ | |
| epithelium development | GO:0060429 | $8.531\times10^{-12}$ | |
| tube development | GO:0035295 | $2.112\times10^{-10}$ | |
| embryo development | GO:0009790 | $1.579\times10^{-9}$ | |
| neuron development | GO:0048666 | $3.015\times10^{-9}$ | |
| anatomical structure formation involved in morphogene... | GO:0048646 | $3.841\times10^{-9}$ | |
| cell morphogenesis | GO:0000902 | $6.270\times10^{-9}$ | |
| Wnt signaling pathway | GO:0016055 | $8.283\times10^{-9}$ | |
| cell morphogenesis involved in neuron differentiation | GO:0048667 | $1.108\times10^{-8}$ | |
| biological regulation | GO:0065007 | $1.109\times10^{-8}$ | |
| axon development | GO:0061564 | $1.410\times10^{-8}$ | |
| neuron projection development | GO:0031175 | $1.866\times10^{-8}$ | |
| regulation of developmental process | GO:0050793 | $2.469\times10^{-8}$ | |
| regulation of cellular process | GO:0050794 | $2.492\times10^{-8}$ | |
| neuron projection morphogenesis | GO:0048812 | $3.483\times10^{-8}$ | |
| plasma membrane bounded cell projection morphogene... | GO:0120039 | $3.483\times10^{-8}$ | |
| cell projection morphogenesis | GO:0048858 | $3.483\times10^{-8}$ | |
| axonogenesis | GO:0007409 | $5.017\times10^{-8}$ | |
| circulatory system development | GO:0072359 | $7.662\times10^{-8}$ | |
| regulation of biological process | GO:0050789 | $1.510\times10^{-7}$ | |
| cellular response to endogenous stimulus | GO:0071495 | $2.735\times10^{-7}$ | |
| response to endogenous stimulus | GO:0009719 | $3.692\times10^{-7}$ | |
| regulation of primary metabolic process | GO:0080090 | $8.037\times10^{-7}$ | |
| sensory organ development | GO:0007423 | $1.419\times10^{-6}$ | |
| phosphate-containing compound metabolic process | GO:0006796 | $2.794\times10^{-6}$ | |
| phosphorus metabolic process | GO:0006793 | $2.794\times10^{-6}$ | |
| tube morphogenesis | GO:0035239 | $3.115\times10^{-6}$ | |
| regulation of RNA metabolic process | GO:0051252 | $6.061\times10^{-6}$ | |
| transmembrane transport | GO:0055085 | $9.876\times10^{-6}$ | |
| regulation of signaling | GO:0023051 | $9.966\times10^{-6}$ | |
| regulation of signal transduction | GO:0009966 | $1.069\times10^{-5}$ | |
| regulation of cell communication | GO:0010646 | $1.096\times10^{-5}$ | |
| sensory system development | GO:0048880 | $1.281\times10^{-5}$ | |
| central nervous system development | GO:0007417 | $1.480\times10^{-5}$ | |
| cellular process | GO:0009987 | $1.861\times10^{-5}$ | |
| regulation of nucleobase-containing compound metabo... | GO:0019219 | $1.929\times10^{-5}$ | |
| enzyme-linked receptor protein signaling pathway | GO:0007167 | $2.882\times10^{-5}$ | |
| vasculature development | GO:0001944 | $5.264\times10^{-5}$ | |
| head development | GO:0060322 | $5.944\times10^{-5}$ | |
| plasma membrane bounded cell projection organization | GO:0120036 | $6.066\times10^{-5}$ | |
| embryonic morphogenesis | GO:0048598 | $6.113\times10^{-5}$ | |
| neuron projection guidance | GO:0097485 | $8.859\times10^{-5}$ | |
| axon guidance | GO:0007411 | $8.859\times10^{-5}$ | |
| cell communication | GO:0007154 | $9.446\times10^{-5}$ | |
| embryo development ending in birth or egg hatching | GO:0009792 | $9.675\times10^{-5}$ | |
| chordate embryonic development | GO:0043009 | $9.675\times10^{-5}$ | |
| blood vessel morphogenesis | GO:0048514 | $1.181\times10^{-4}$ | |
| regulation of DNA-templated transcription | GO:0006355 | $1.410\times10^{-4}$ | |
| regulation of RNA biosynthetic process | GO:2001141 | $1.410\times10^{-4}$ | |
| DNA-templated transcription | GO:0006351 | $1.619\times10^{-4}$ | |
| brain development | GO:0007420 | $2.156\times10^{-4}$ | |
| intracellular signaling cassette | GO:0141124 | $2.227\times10^{-4}$ | |
| angiogenesis | GO:0001525 | $2.333\times10^{-4}$ | |
| muscle structure development | GO:0061061 | $3.040\times10^{-4}$ | |
| cell projection organization | GO:0030030 | $3.291\times10^{-4}$ | |
| blood vessel development | GO:0001568 | $3.736\times10^{-4}$ | |
| regulation of multicellular organismal development | GO:2000026 | $5.990\times10^{-4}$ | |
| vesicle-mediated transport | GO:0016192 | $1.121\times10^{-3}$ | |
| response to growth factor | GO:0070848 | $1.216\times10^{-3}$ | |
| cellular response to growth factor stimulus | GO:0071363 | $1.216\times10^{-3}$ | |
| signaling | GO:0023052 | $1.230\times10^{-3}$ | |
| monoatomic ion transport | GO:0006811 | $1.324\times10^{-3}$ | |
| negative regulation of signaling | GO:0023057 | $1.448\times10^{-3}$ | |
| negative regulation of cell communication | GO:0010648 | $1.448\times10^{-3}$ | |
| signal transduction | GO:0007165 | $1.859\times10^{-3}$ | |
| negative regulation of signal transduction | GO:0009968 | $2.230\times10^{-3}$ | |
| morphogenesis of an epithelium | GO:0002009 | $2.550\times10^{-3}$ | |
| positive regulation of cellular process | GO:0048522 | $3.528\times10^{-3}$ | |
| regulation of Wnt signaling pathway | GO:0030111 | $3.612\times10^{-3}$ | |
| epidermis development | GO:0008544 | $4.006\times10^{-3}$ | |
| monoatomic ion transmembrane transport | GO:0034220 | $4.150\times10^{-3}$ | |
| epithelial cell differentiation | GO:0030855 | $5.087\times10^{-3}$ | |
| regulation of cell differentiation | GO:0045595 | $6.218\times10^{-3}$ | |
| regulation of biological quality | GO:0065008 | $9.811\times10^{-3}$ | |
| negative regulation of response to stimulus | GO:0048585 | $1.048\times10^{-2}$ | |
| canonical Wnt signaling pathway | GO:0060070 | $1.150\times10^{-2}$ | |
| protein phosphorylation | GO:0006468 | $1.495\times10^{-2}$ | |
| organophosphate metabolic process | GO:0019637 | $1.835\times10^{-2}$ | |
| phosphorylation | GO:0016310 | $1.847\times10^{-2}$ | |
| eye development | GO:0001654 | $1.882\times10^{-2}$ | |
| visual system development | GO:0150063 | $1.882\times10^{-2}$ | |
| heart development | GO:0007507 | $2.142\times10^{-2}$ | |
| ameboidal-type cell migration | GO:0001667 | $2.231\times10^{-2}$ | |

### (C) *A. burtoni* Forebrain

| GO:BP | | stats | |
|---|---|---|---|
| Term name | Term ID | $p_{adj}$ | $-\log_{10}(p_{adj})$ (0 … ≤16) |
| system development | GO:0048731 | $2.981\times10^{-14}$ | |
| multicellular organism development | GO:0007275 | $9.199\times10^{-13}$ | |
| anatomical structure development | GO:0048856 | $3.857\times10^{-10}$ | |
| nervous system development | GO:0007399 | $4.212\times10^{-10}$ | |
| developmental process | GO:0032502 | $1.123\times10^{-9}$ | |
| cell differentiation | GO:0030154 | $4.099\times10^{-9}$ | |
| cellular developmental process | GO:0048869 | $4.403\times10^{-9}$ | |
| neurogenesis | GO:0022008 | $5.667\times10^{-7}$ | |
| generation of neurons | GO:0048699 | $3.584\times10^{-6}$ | |
| neuron differentiation | GO:0030182 | $4.243\times10^{-6}$ | |
| transport | GO:0006810 | $5.412\times10^{-6}$ | |
| localization | GO:0051179 | $6.258\times10^{-6}$ | |
| establishment of localization | GO:0051234 | $1.285\times10^{-5}$ | |
| cellular process | GO:0009987 | $2.558\times10^{-5}$ | |
| monoatomic ion transmembrane transport | GO:0034220 | $3.056\times10^{-5}$ | |
| multicellular organismal process | GO:0032501 | $3.879\times10^{-5}$ | |
| transmembrane transport | GO:0055085 | $1.789\times10^{-4}$ | |
| cell development | GO:0048468 | $4.402\times10^{-4}$ | |
| response to hormone | GO:0009725 | $4.506\times10^{-4}$ | |
| inorganic ion transmembrane transport | GO:0098660 | $6.183\times10^{-4}$ | |
| response to endogenous stimulus | GO:0009719 | $6.717\times10^{-4}$ | |
| cellular response to hormone stimulus | GO:0032870 | $7.278\times10^{-4}$ | |
| cellular response to endogenous stimulus | GO:0071495 | $1.206\times10^{-3}$ | |
| animal organ development | GO:0048513 | $2.331\times10^{-3}$ | |
| enzyme-linked receptor protein signaling pathway | GO:0007167 | $2.350\times10^{-3}$ | |
| homeostasis of number of cells | GO:0048872 | $4.358\times10^{-3}$ | |
| regulation of cell communication | GO:0010646 | $4.489\times10^{-3}$ | |
| monoatomic anion transmembrane transport | GO:0098656 | $5.269\times10^{-3}$ | |
| regulation of signaling | GO:0023051 | $6.950\times10^{-3}$ | |
| monoatomic ion transport | GO:0006811 | $8.812\times10^{-3}$ | |
| myeloid cell differentiation | GO:0030099 | $9.018\times10^{-3}$ | |
| cell surface receptor protein tyrosine kinase signaling p... | GO:0007169 | $1.276\times10^{-2}$ | |
| monoatomic anion transport | GO:0006820 | $1.364\times10^{-2}$ | |
| myeloid cell homeostasis | GO:0002262 | $1.586\times10^{-2}$ | |
| positive regulation of cellular process | GO:0048522 | $1.596\times10^{-2}$ | |
| sensory system development | GO:0048880 | $1.974\times10^{-2}$ | |
| hemopoiesis | GO:0030097 | $2.055\times10^{-2}$ | |
| regulation of signal transduction | GO:0009966 | $2.230\times10^{-2}$ | |
| erythrocyte differentiation | GO:0030218 | $2.696\times10^{-2}$ | |
| erythrocyte homeostasis | GO:0034101 | $2.696\times10^{-2}$ | |
| eye development | GO:0001654 | $3.512\times10^{-2}$ | |
| visual system development | GO:0150063 | $3.512\times10^{-2}$ | |
| inorganic anion transport | GO:0015698 | $4.303\times10^{-2}$ | |
| vasculature development | GO:0001944 | $4.650\times10^{-2}$ | |
| hypothalamus development | GO:0021854 | $4.923\times10^{-2}$ | |
| tube morphogenesis | GO:0035239 | $4.940\times10^{-2}$ | |

### (D) *N. brichardi* Forebrain

| GO:BP | | stats | |
|---|---|---|---|
| Term name | Term ID | $p_{adj}$ | $-\log_{10}(p_{adj})$ (0 … ≤16) |
| anatomical structure development | GO:0048856 | $2.646\times10^{-38}$ | |
| developmental process | GO:0032502 | $9.440\times10^{-37}$ | |
| multicellular organism development | GO:0007275 | $1.949\times10^{-31}$ | |
| animal organ development | GO:0048513 | $1.429\times10^{-24}$ | |
| system development | GO:0048731 | $5.869\times10^{-24}$ | |
| cell differentiation | GO:0030154 | $1.333\times10^{-18}$ | |
| tissue development | GO:0009888 | $1.545\times10^{-18}$ | |
| cellular developmental process | GO:0048869 | $1.569\times10^{-18}$ | |
| anatomical structure morphogenesis | GO:0009653 | $2.042\times10^{-18}$ | |
| multicellular organismal process | GO:0032501 | $1.926\times10^{-16}$ | |
| epithelium development | GO:0060429 | $6.885\times10^{-14}$ | |
| nervous system development | GO:0007399 | $1.213\times10^{-12}$ | |
| embryo development | GO:0009790 | $1.474\times10^{-12}$ | |
| localization | GO:0051179 | $7.621\times10^{-12}$ | |
| transport | GO:0006810 | $2.117\times10^{-11}$ | |
| generation of neurons | GO:0048699 | $1.106\times10^{-10}$ | |
| neurogenesis | GO:0022008 | $1.371\times10^{-10}$ | |
| cell development | GO:0048468 | $1.767\times10^{-10}$ | |
| establishment of localization | GO:0051234 | $2.324\times10^{-10}$ | |
| neuron differentiation | GO:0030182 | $5.781\times10^{-10}$ | |
| central nervous system development | GO:0007417 | $3.672\times10^{-9}$ | |
| cellular process | GO:0009987 | $1.823\times10^{-8}$ | |
| Wnt signaling pathway | GO:0016055 | $2.561\times10^{-8}$ | |
| sensory organ development | GO:0007423 | $6.016\times10^{-8}$ | |
| tissue morphogenesis | GO:0048729 | $8.467\times10^{-8}$ | |
| regulation of cell communication | GO:0010646 | $1.353\times10^{-7}$ | |
| anatomical structure formation involved in morphogene... | GO:0048646 | $1.574\times10^{-7}$ | |
| sensory system development | GO:0048880 | $1.827\times10^{-7}$ | |
| regulation of signaling | GO:0023051 | $2.053\times10^{-7}$ | |
| regulation of primary metabolic process | GO:0080090 | $3.158\times10^{-7}$ | |
| regulation of signal transduction | GO:0009966 | $4.110\times10^{-7}$ | |
| regulation of RNA metabolic process | GO:0051252 | $1.351\times10^{-6}$ | |
| embryonic morphogenesis | GO:0048598 | $1.955\times10^{-6}$ | |
| biological regulation | GO:0065007 | $2.038\times10^{-6}$ | |
| embryo development ending in birth or egg hatching | GO:0009792 | $4.084\times10^{-6}$ | |
| chordate embryonic development | GO:0043009 | $4.084\times10^{-6}$ | |
| regulation of nucleobase-containing compound metabo... | GO:0019219 | $5.128\times10^{-6}$ | |
| animal organ morphogenesis | GO:0009887 | $6.078\times10^{-6}$ | |
| tube development | GO:0035295 | $6.893\times10^{-6}$ | |
| regulation of cellular process | GO:0050794 | $7.239\times10^{-6}$ | |
| regulation of biological process | GO:0050789 | $8.928\times10^{-6}$ | |
| morphogenesis of an epithelium | GO:0002009 | $1.141\times10^{-5}$ | |
| regulation of RNA biosynthetic process | GO:2001141 | $2.018\times10^{-5}$ | |
| regulation of DNA-templated transcription | GO:0006355 | $2.018\times10^{-5}$ | |
| vesicle-mediated transport | GO:0016192 | $3.017\times10^{-5}$ | |
| circulatory system development | GO:0072359 | $4.575\times10^{-5}$ | |
| neuron development | GO:0048666 | $4.703\times10^{-5}$ | |
| camera-type eye development | GO:0043010 | $5.676\times10^{-5}$ | |
| DNA-templated transcription | GO:0006351 | $7.597\times10^{-5}$ | |
| visual system development | GO:0150063 | $1.055\times10^{-4}$ | |
| eye development | GO:0001654 | $1.055\times10^{-4}$ | |
| head development | GO:0060322 | $1.733\times10^{-4}$ | |
| axon development | GO:0061564 | $1.902\times10^{-4}$ | |
| cellular localization | GO:0051641 | $2.928\times10^{-4}$ | |
| establishment of localization in cell | GO:0051649 | $4.520\times10^{-4}$ | |
| embryonic organ development | GO:0048568 | $4.781\times10^{-4}$ | |
| plasma membrane bounded cell projection organization | GO:0120036 | $4.972\times10^{-4}$ | |
| regulation of developmental process | GO:0050793 | $5.760\times10^{-4}$ | |
| brain development | GO:0007420 | $6.930\times10^{-4}$ | |
| cell fate commitment | GO:0045165 | $9.559\times10^{-4}$ | |
| regionalization | GO:0003002 | $1.016\times10^{-3}$ | |
| retina development in camera-type eye | GO:0060041 | $1.024\times10^{-3}$ | |
| phosphate-containing compound metabolic process | GO:0006796 | $1.102\times10^{-3}$ | |
| phosphorus metabolic process | GO:0006793 | $1.102\times10^{-3}$ | |
| macromolecule localization | GO:0033036 | $1.431\times10^{-3}$ | |
| cell projection morphogenesis | GO:0048858 | $1.620\times10^{-3}$ | |
| plasma membrane bounded cell projection morphogene... | GO:0120039 | $1.620\times10^{-3}$ | |
| neuron projection morphogenesis | GO:0048812 | $1.620\times10^{-3}$ | |
| axonogenesis | GO:0007409 | $1.779\times10^{-3}$ | |
| protein transport | GO:0015031 | $2.081\times10^{-3}$ | |
| negative regulation of signal transduction | GO:0009968 | $2.220\times10^{-3}$ | |
| pattern specification process | GO:0007389 | $2.494\times10^{-3}$ | |
| cell morphogenesis | GO:0000902 | $2.530\times10^{-3}$ | |
| negative regulation of cell communication | GO:0010648 | $2.743\times10^{-3}$ | |
| negative regulation of signaling | GO:0023057 | $2.743\times10^{-3}$ | |
| tube morphogenesis | GO:0035239 | $2.839\times10^{-3}$ | |
| negative regulation of response to stimulus | GO:0048585 | $3.027\times10^{-3}$ | |
| nitrogen compound transport | GO:0071705 | $3.136\times10^{-3}$ | |
| positive regulation of cellular process | GO:0048522 | $3.178\times10^{-3}$ | |
| cell morphogenesis involved in neuron differentiation | GO:0048667 | $3.914\times10^{-3}$ | |
| regulation of multicellular organismal development | GO:2000026 | $4.855\times10^{-3}$ | |
| neuron projection development | GO:0031175 | $4.920\times10^{-3}$ | |
| regulation of transcription by RNA polymerase II | GO:0006357 | $5.337\times10^{-3}$ | |
| growth | GO:0040007 | $5.428\times10^{-3}$ | |
| cell-cell signaling | GO:0007267 | $5.917\times10^{-3}$ | |
| developmental growth | GO:0048589 | $6.081\times10^{-3}$ | |
| cell communication | GO:0007154 | $6.843\times10^{-3}$ | |
| cell projection organization | GO:0030030 | $6.885\times10^{-3}$ | |
| enzyme-linked receptor protein signaling pathway | GO:0007167 | $6.921\times10^{-3}$ | |
| chemical synaptic transmission | GO:0007268 | $7.063\times10^{-3}$ | |
| anterograde trans-synaptic signaling | GO:0098916 | $7.063\times10^{-3}$ | |
| trans-synaptic signaling | GO:0099537 | $7.063\times10^{-3}$ | |
| central nervous system neuron differentiation | GO:0021953 | $7.342\times10^{-3}$ | |
| cell fate specification | GO:0001708 | $7.630\times10^{-3}$ | |
| synaptic signaling | GO:0099536 | $7.685\times10^{-3}$ | |
| transcription by RNA polymerase II | GO:0006366 | $7.819\times10^{-3}$ | |
| sensory organ morphogenesis | GO:0090596 | $8.199\times10^{-3}$ | |
| cellular response to endogenous stimulus | GO:0071495 | $9.549\times10^{-3}$ | |
| protein modification process | GO:0036211 | $1.014\times10^{-2}$ | |
| canonical Wnt signaling pathway | GO:0060070 | $1.058\times10^{-2}$ | |

**Fig. S8 – Gene Ontology (GO) enrichment of gene promoter peak gains versus *O. niloticus* peaks in forebrain tissue of four species.** Bar plot (with table) of significance (-$log_{10}$ FDR <0.05, y-axis) of each Nile tilapia term shown for each species. Table of terms calculated by the g:GOSt module of g:Profiler (https://biit.cs.ut.ee/gprofiler/gost).

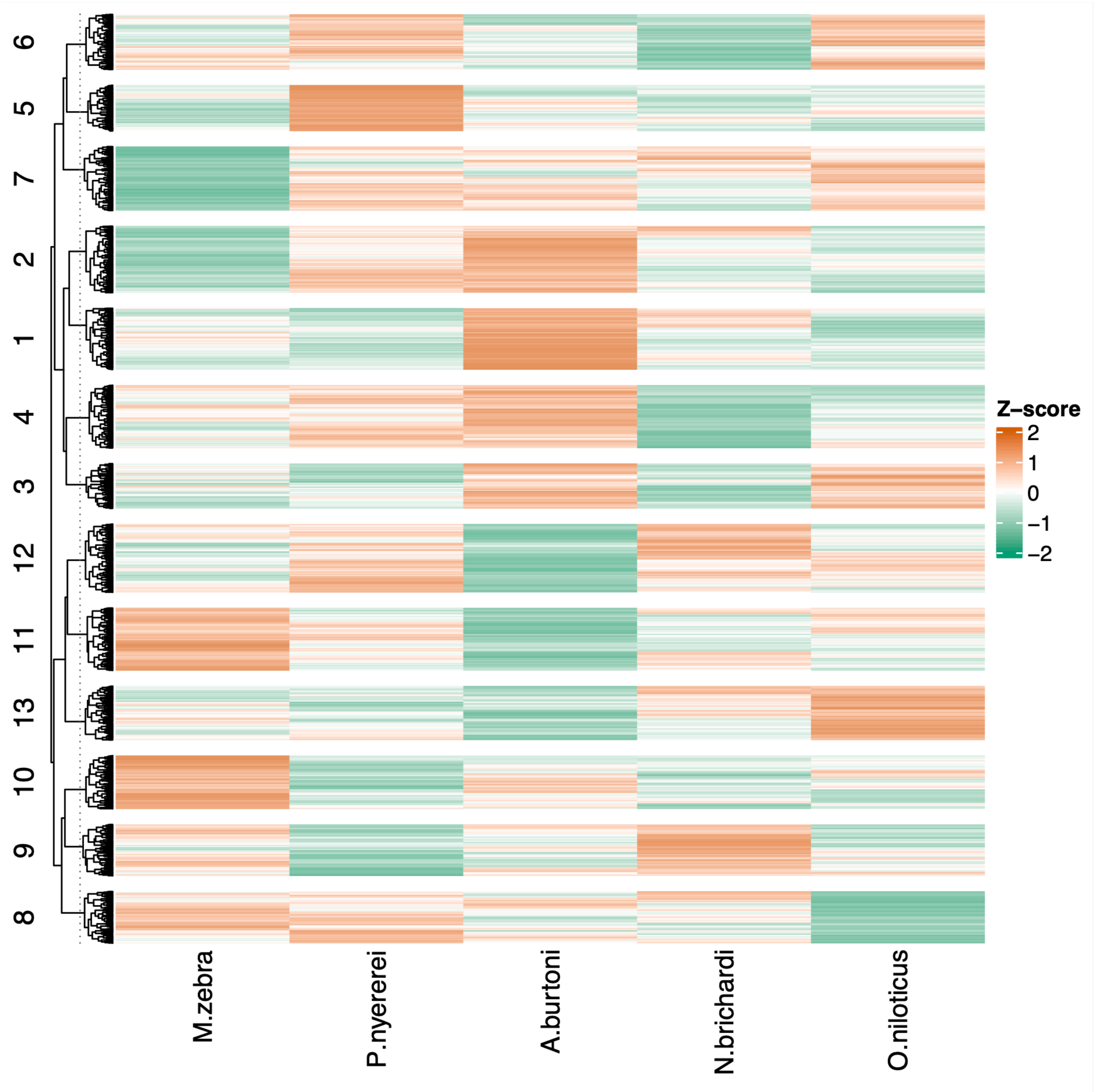


**Fig. S9 – Clustering of TF activities in forebrain tissue across the five species.** Each row represents a single, clustered TF, and each column a species. Mean TF bit-score of motif match across gene promoters (as a measure of activity) from HINT-ATAC [1] are *Z*-score transformed across rows. Green colour indicates low activity and orange colour indicates high activity.

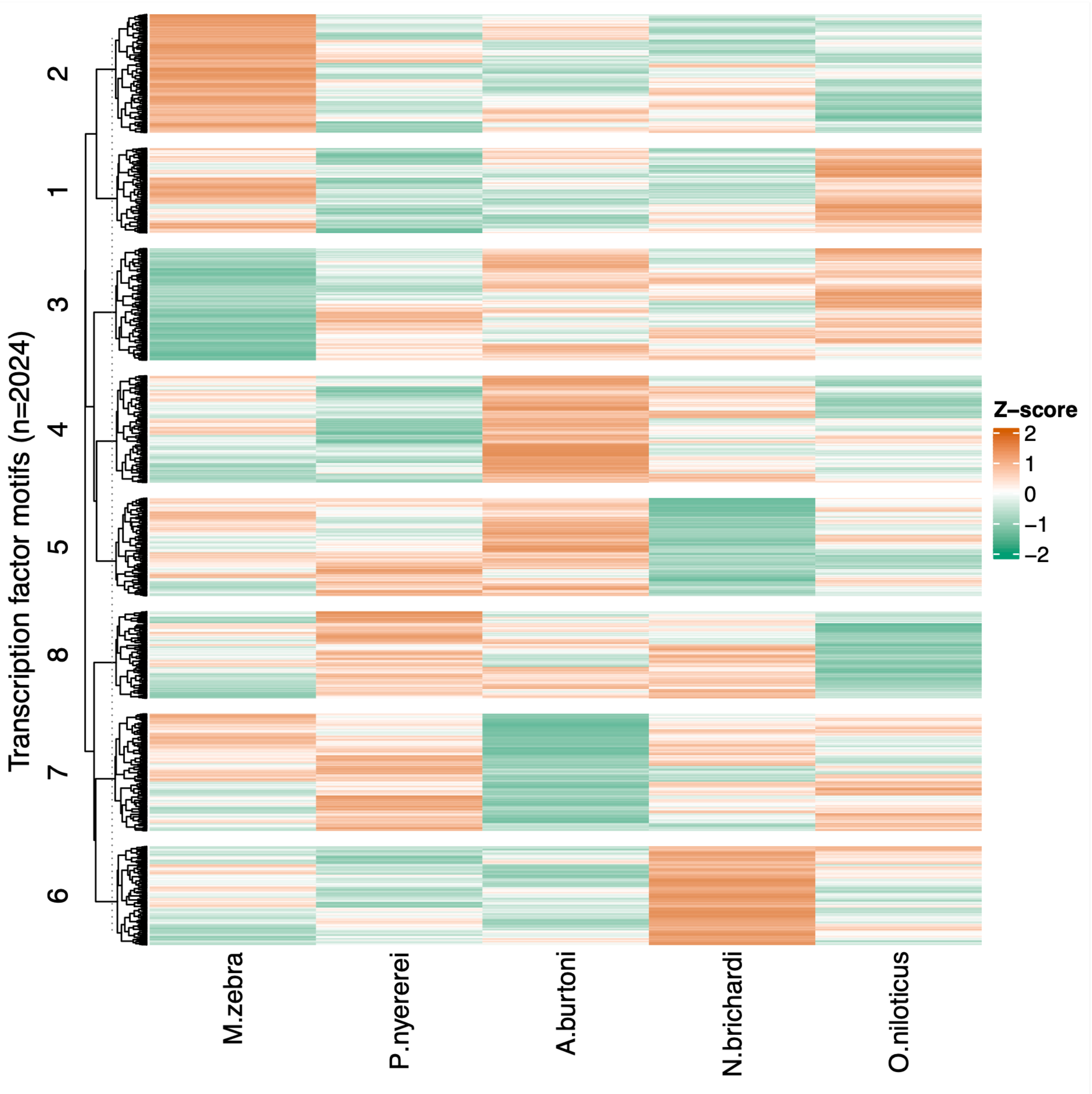


**Fig. S10 – Clustering of TF activities in liver tissue across the five species.**

Each row represents a single, clustered TF, and each column a species. Mean TF bit-score of motif match across gene promoters (as a measure of activity) from HINT-ATAC [1] are *Z*-score transformed across rows. Green colour indicates low activity and orange colour indicates high activity.

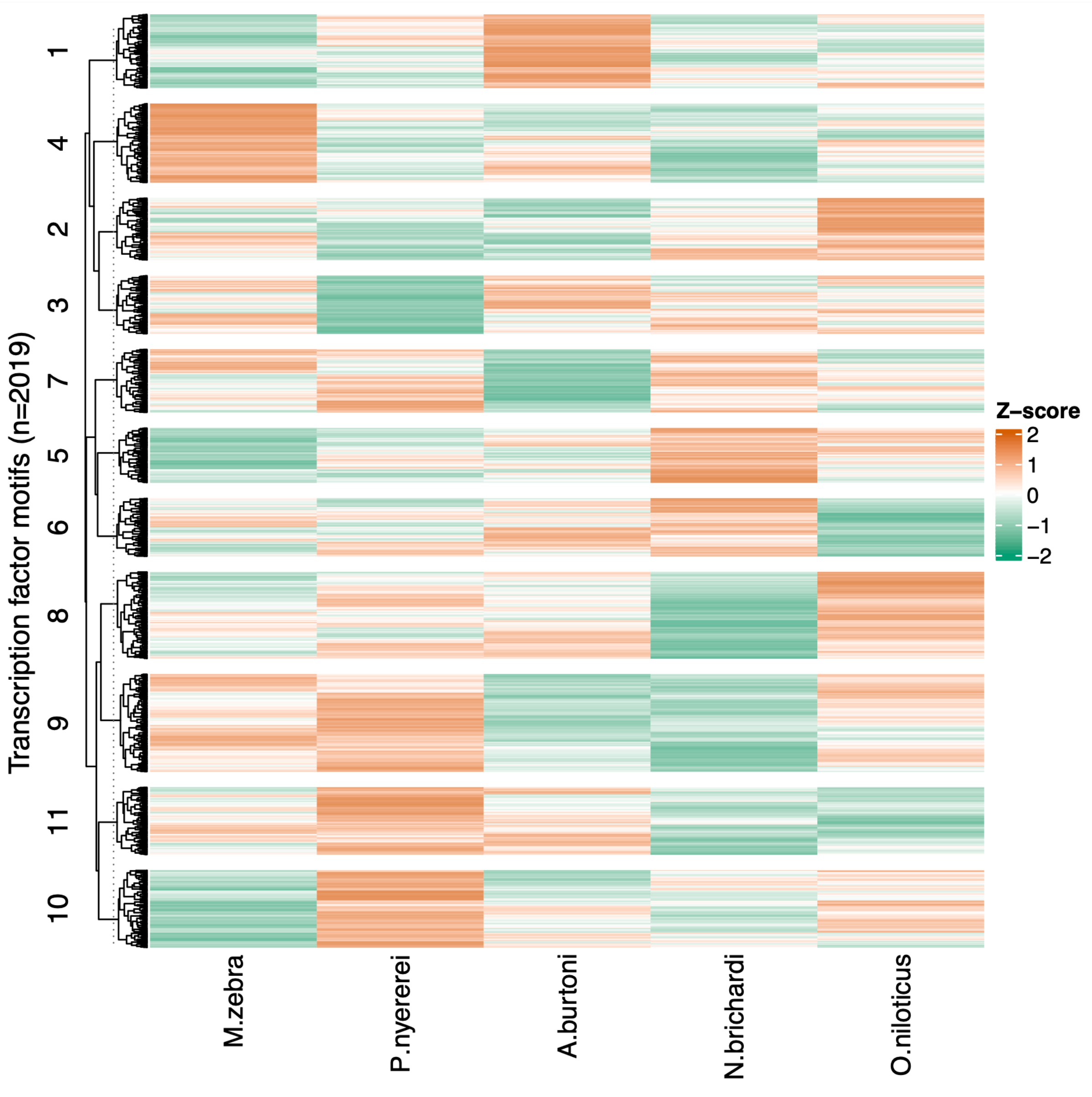


**Fig. S11 – Clustering of TF activities in testis tissue across the five species.** Each row represents a single, clustered TF, and each column a species. Mean TF bit-score of motif match across gene promoters (as a measure of activity) from HINT-ATAC [1] are *Z*-score transformed across rows. Green colour indicates low activity and orange colour indicates high activity.

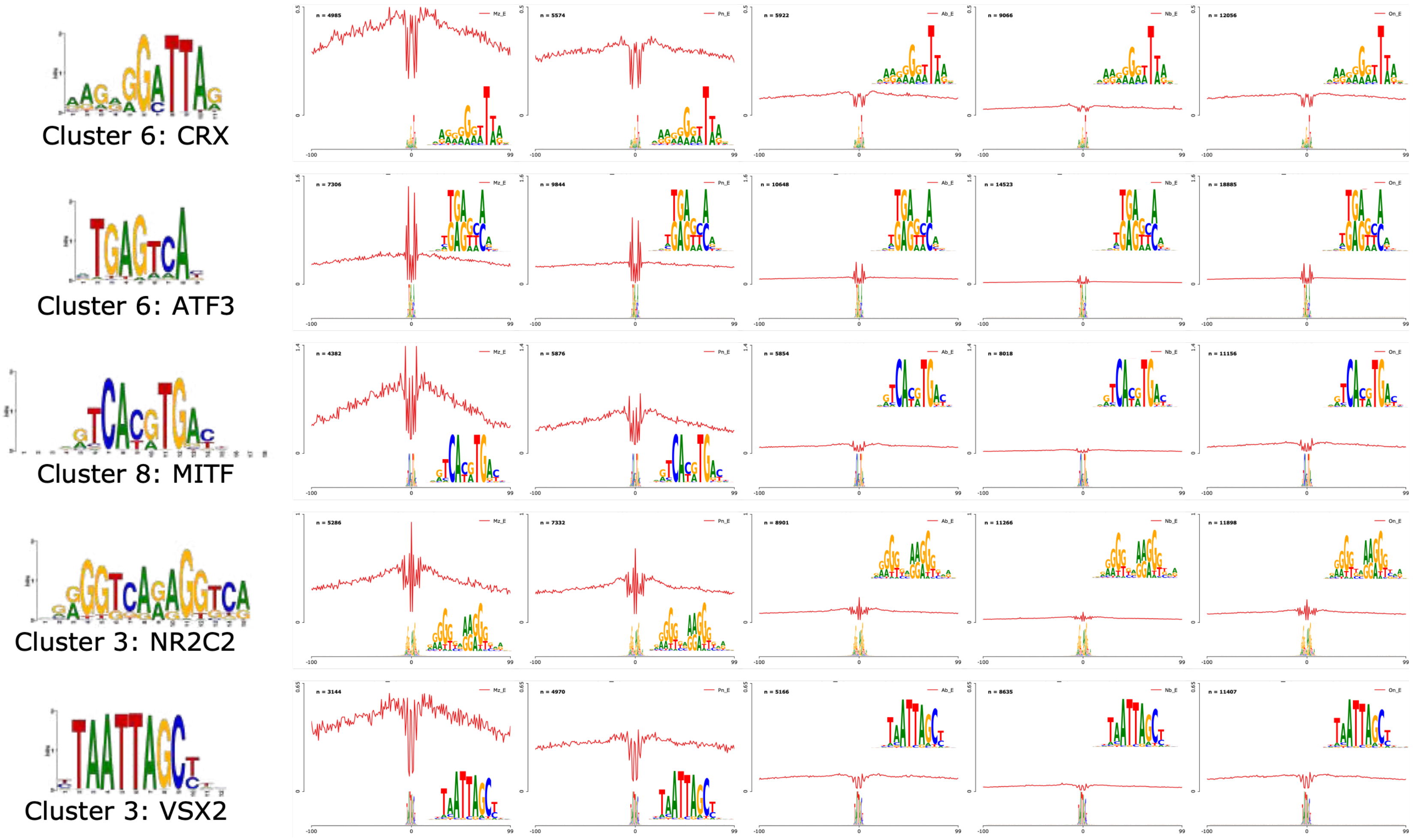


**Fig. S12 – Activity profile of five eye-related TFs across retina tissue of five cichlid species.** Line plot profile of the CRX, ATF3, MITF, NR2C2, and VSX2 TF footprints across retina tissue of five cichlid species (Mz – *M. zebra;* Pn – *P. nyererei;* Ab – *A.*

*burtoni;* Nb – *N. brichardi;* On – *O. niloticus*) gene promoter regions showing positional (x-axis) signal of activity (red line, y-axis) centred on the footprint (at position 0) with flanking sequence. Number (n=) of footprints profiled for each TF shown on top left of each plot. Expanded TF footprint shown with each line plot and original position weight matrices (PWM) of input motif used for profiling the site shown to left, with retina cluster assignment (from Fig. 2C and Fig. 2D).

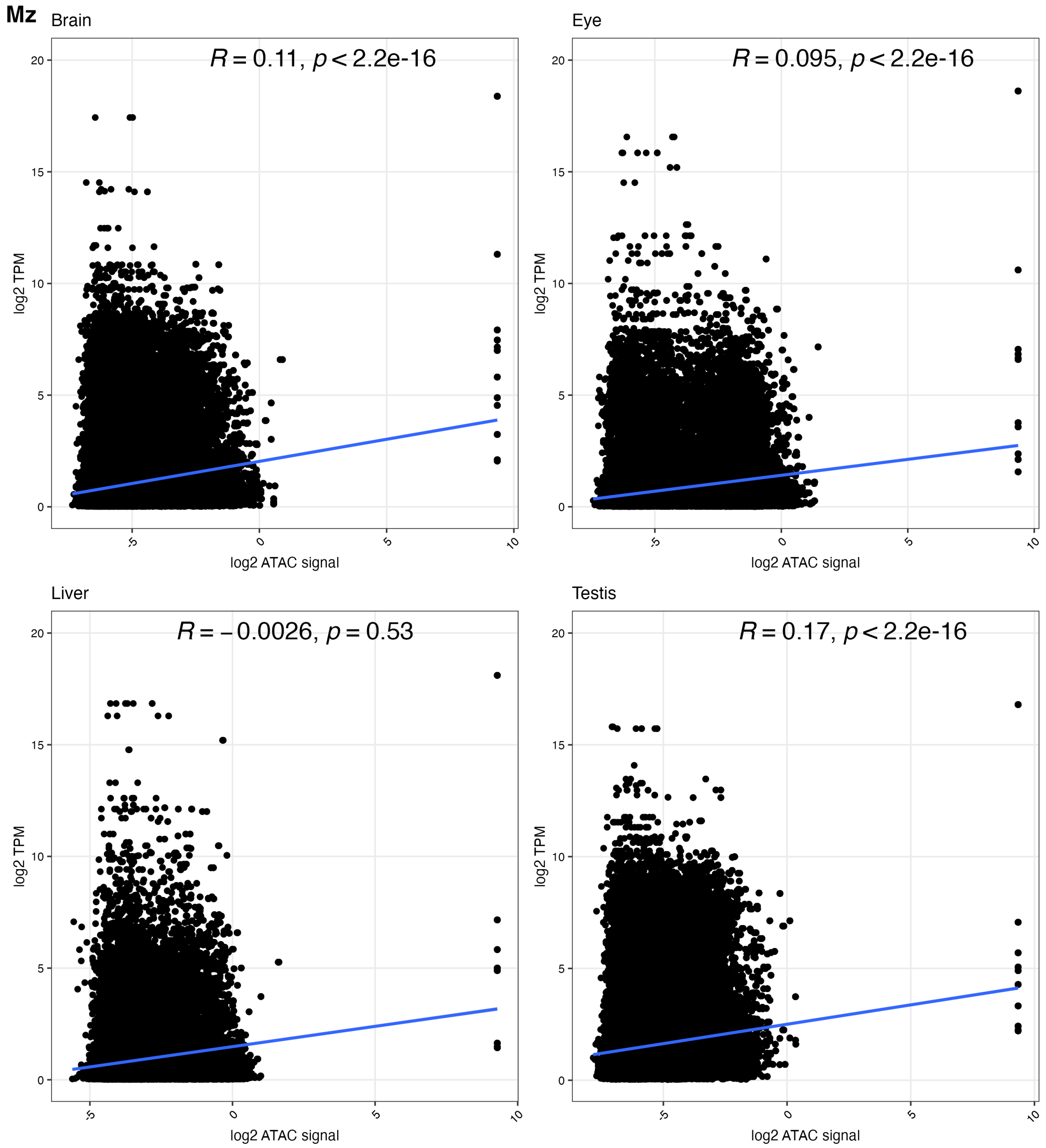


**Fig. S13 – Correlation of accessible peaks accounting for gene expression change in *M. zebra* tissues.** Average log2 gene expression as transcripts per million (TPM, y-axis) and average log2 ATAC signal as gene promoter peak counts (x-axis) for each gene across biological replicates, split by tissue (Brain = Forebrain and Eye = Retina). Correlation co-efficient (*R,* blue trendline) and *p-value* shown for each category.

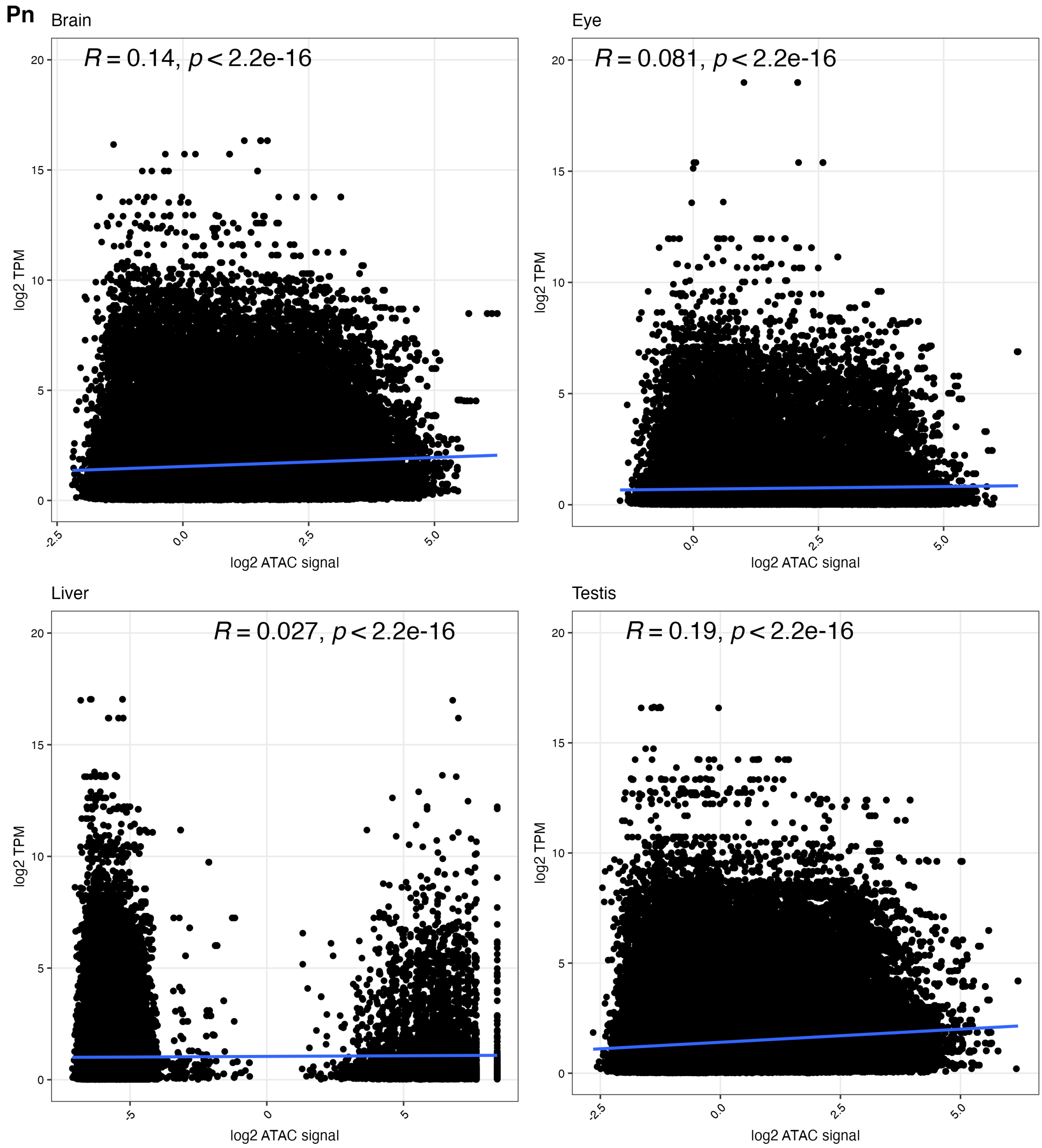


**Fig. S14 – Correlation of accessible peaks accounting for gene expression change in *P. nyererei* tissues.** Average log2 gene expression as transcripts per million (TPM, y-axis) and average log2 ATAC signal as gene promoter peak counts (x-axis) for each gene across biological replicates, split by tissue (Brain = Forebrain and Eye = Retina). Correlation co-efficient (*R,* blue trendline) and *p-value* shown for each category.

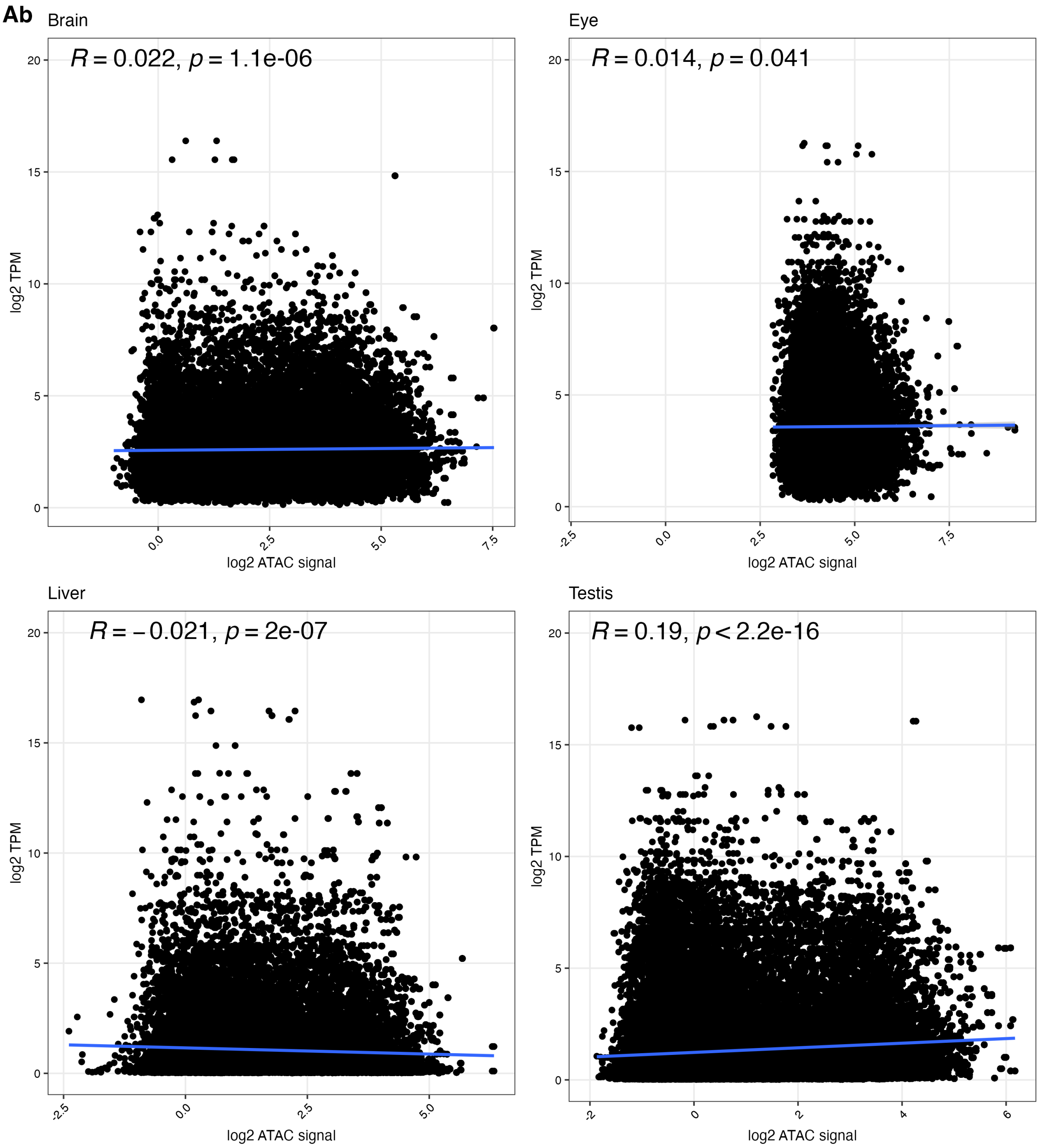


**Fig. S15 – Correlation of accessible peaks accounting for gene expression change in *A. burtoni* tissues.** Average log2 gene expression as transcripts per million (TPM, y-axis) and average log2 ATAC signal as gene promoter peak counts (x-axis) for each gene across biological replicates, split by tissue (Brain = Forebrain and Eye = Retina). Correlation co-efficient (*R,* blue trendline) and *p-value* shown for each category.

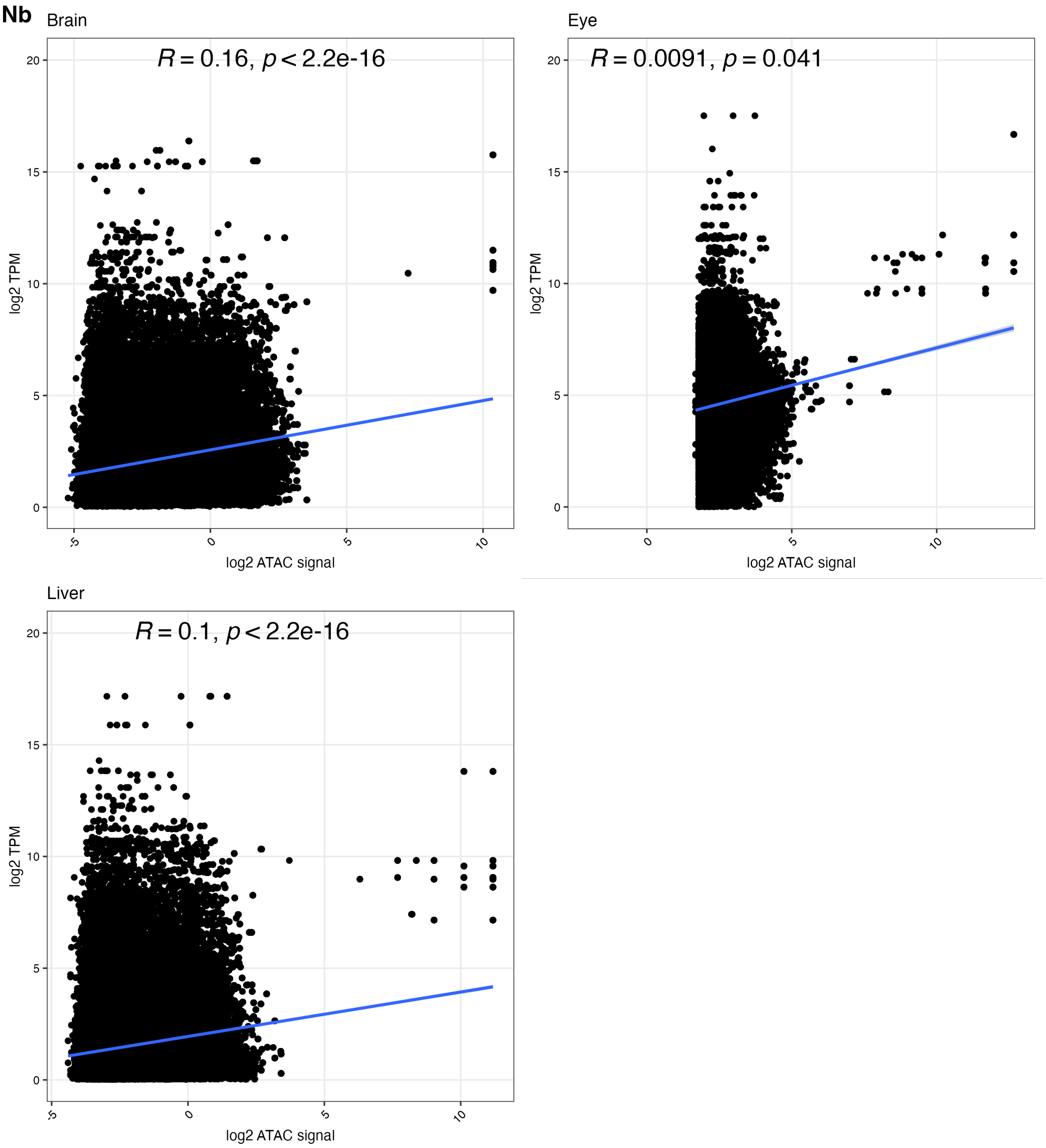


**Fig. S16 – Correlation of accessible peaks accounting for gene expression change in *N. brichardi* tissues.** Average log*2* gene expression as transcripts per million (TPM, y-axis) and average log2 ATAC signal as gene promoter peak counts (x-axis) for each gene across biological replicates, split by tissue (Brain = Forebrain and Eye = Retina). Correlation co-efficient (*R,* blue trendline) and *p-value* shown for each category.

**On**

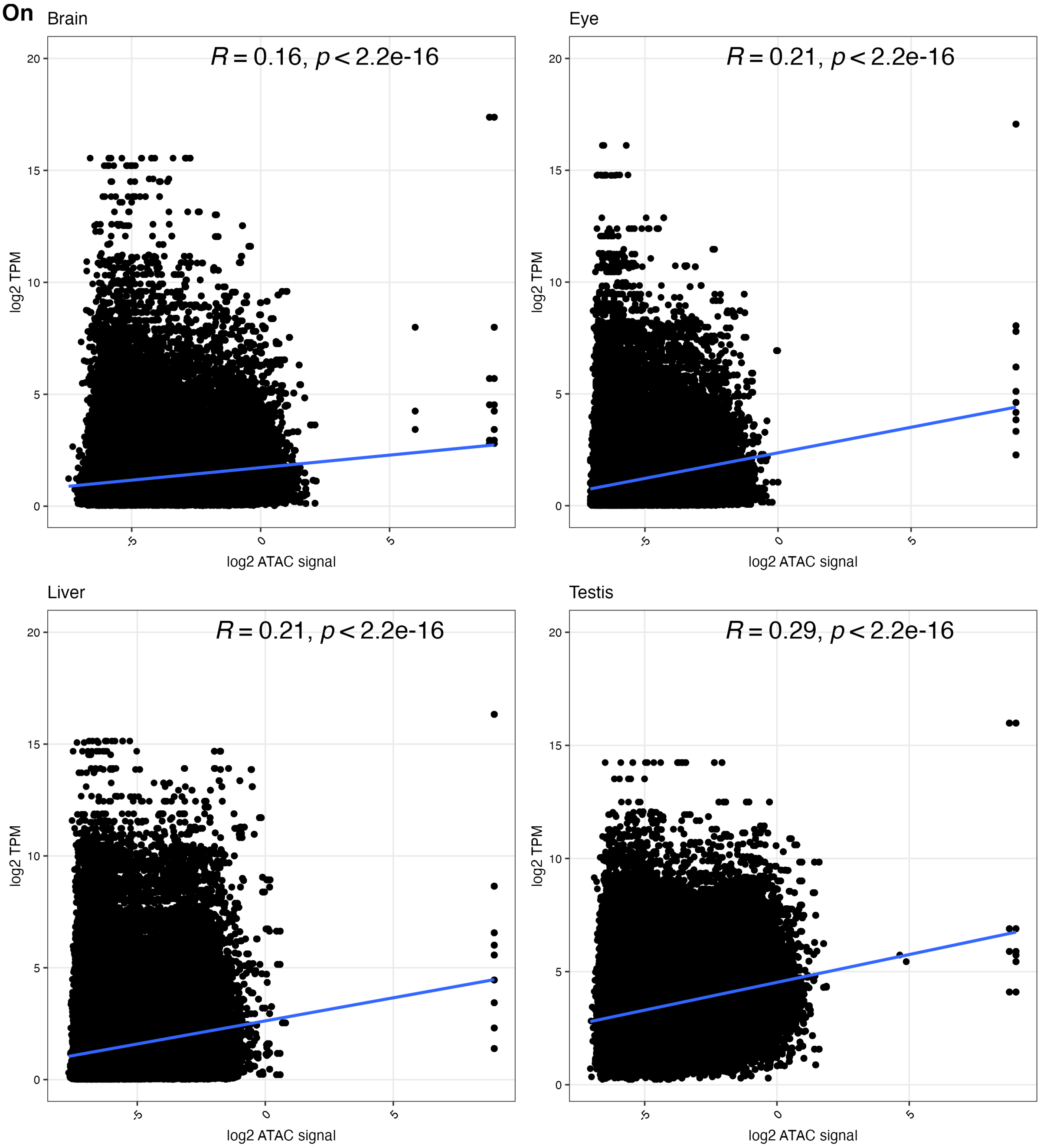


**Fig. S17 – Correlation of accessible peaks accounting for gene expression change in *O. niloticus* tissues.** Average log*2* gene expression as transcripts per million (TPM, y-axis) and average log2 ATAC signal as gene promoter peak counts (x-axis) for each gene across biological replicates, split by tissue (Brain = Forebrain and Eye = Retina). Correlation co-efficient (*R,* blue trendline) and *p-value* shown for each category.

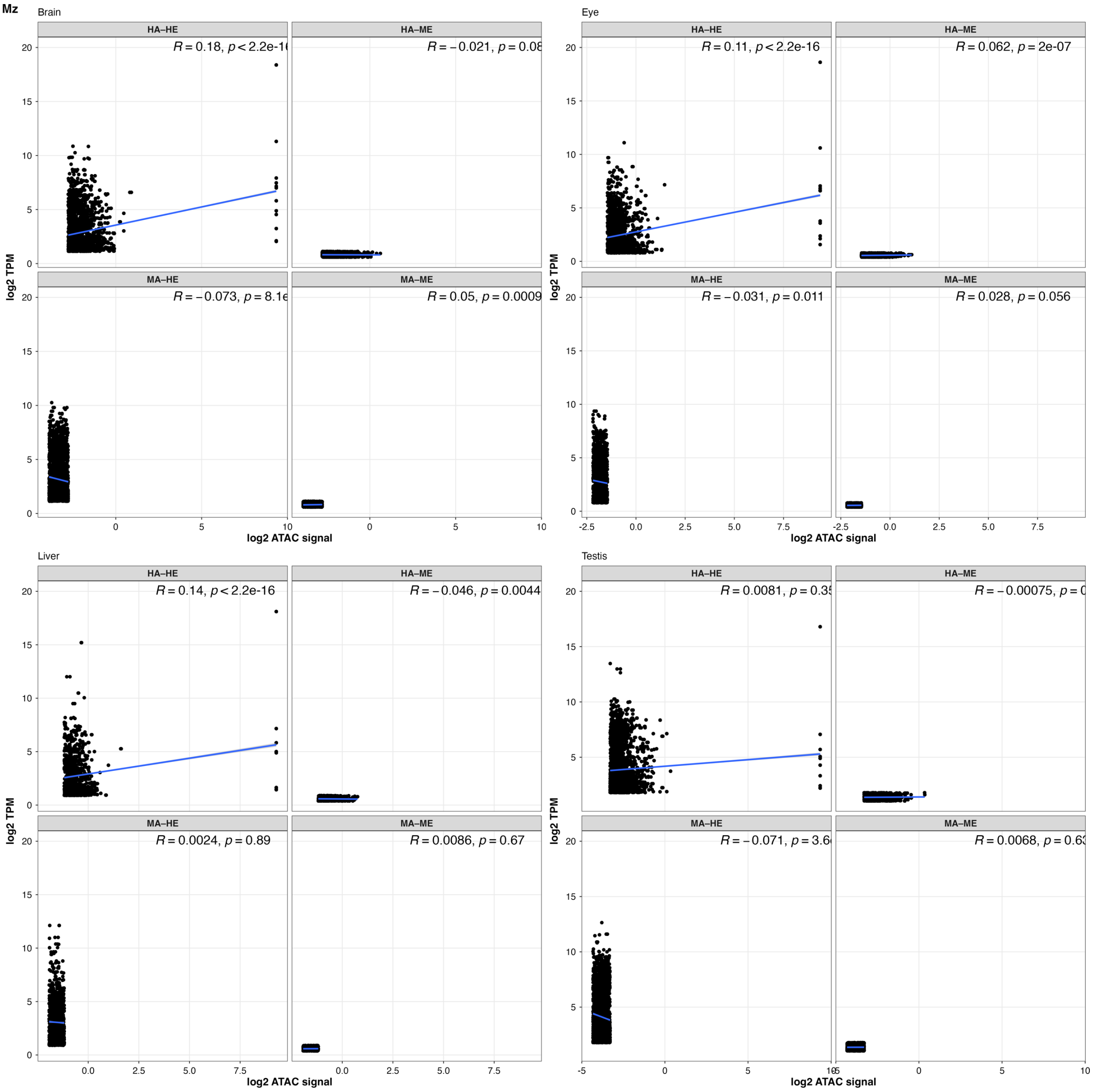


**Fig. S18 – Categorised correlation of accessible peaks accounting for gene expression change in *M. zebra* tissues.** Tissue-specific average *log2* gene expression as transcripts per million (TPM, y-axis) and average *log2* ATAC signal as gene promoter peak counts (x-axis) for each gene across same tissue biological replicates (Brain = Forebrain and Eye = Retina), split by high accessibility and high expression (HA-HE - both average TPM and ATAC signal values are more than the 70th percentiles), high accessibility and medium-low expression (HA-ME - ATAC signal values more than the 70th percentiles and average TPM less than the 50th

percentile), medium-low accessibility and high expression (MA-HE - ATAC signal values less than the 50th percentiles and average TPM more than the 70th percentile), and medium-low accessibility and medium-low expression (MA-ME - average ATAC signal count less than the 50th percentile and average TPM more than the 70th percentile). Correlation co-efficient (*R,* blue trendline) and *p-value* shown for each category.

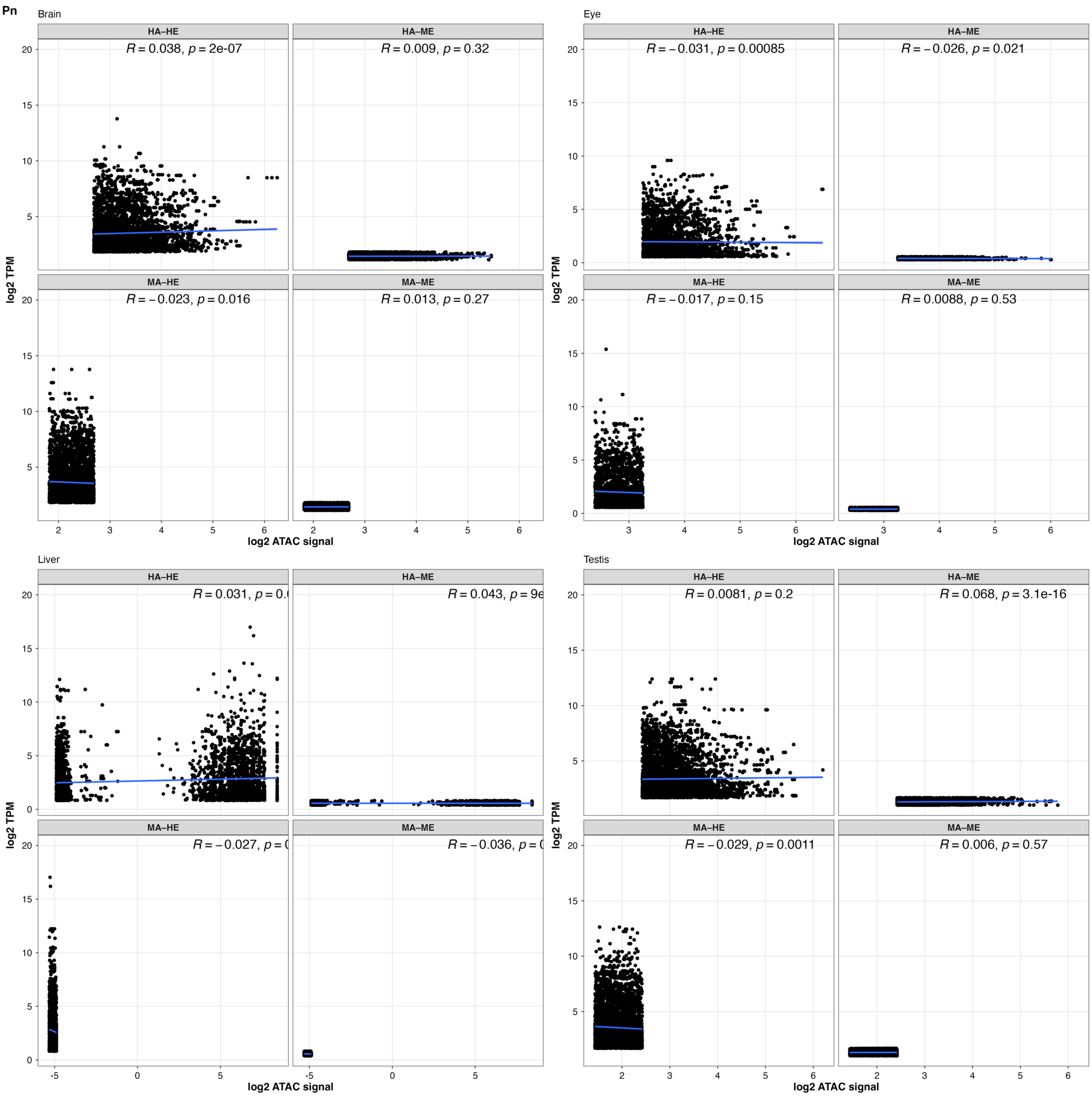


**Fig. S19 – Categorised correlation of accessible peaks accounting for gene expression change in *P. nyererei* tissues.** Tissue-specific average *log2* gene expression as transcripts per million (TPM, y-axis) and average *log2* ATAC signal as gene promoter peak counts (x-axis) for each gene across same tissue biological replicates (Brain = Forebrain and Eye = Retina), split by high accessibility and high expression (HA-HE - both average TPM and ATAC signal values are more than the 70th percentiles), high accessibility and medium-low expression (HA-ME - ATAC signal values more than the 70th percentiles and average TPM less than the 50th

percentile), medium-low accessibility and high expression (MA-HE - ATAC signal values less than the 50th percentiles and average TPM more than the 70th percentile), and medium-low accessibility and medium-low expression (MA-ME - average ATAC signal count less than the 50th percentile and average TPM more than the 70th percentile). Correlation co-efficient (*R,* blue trendline) and *p-value* shown for each category.

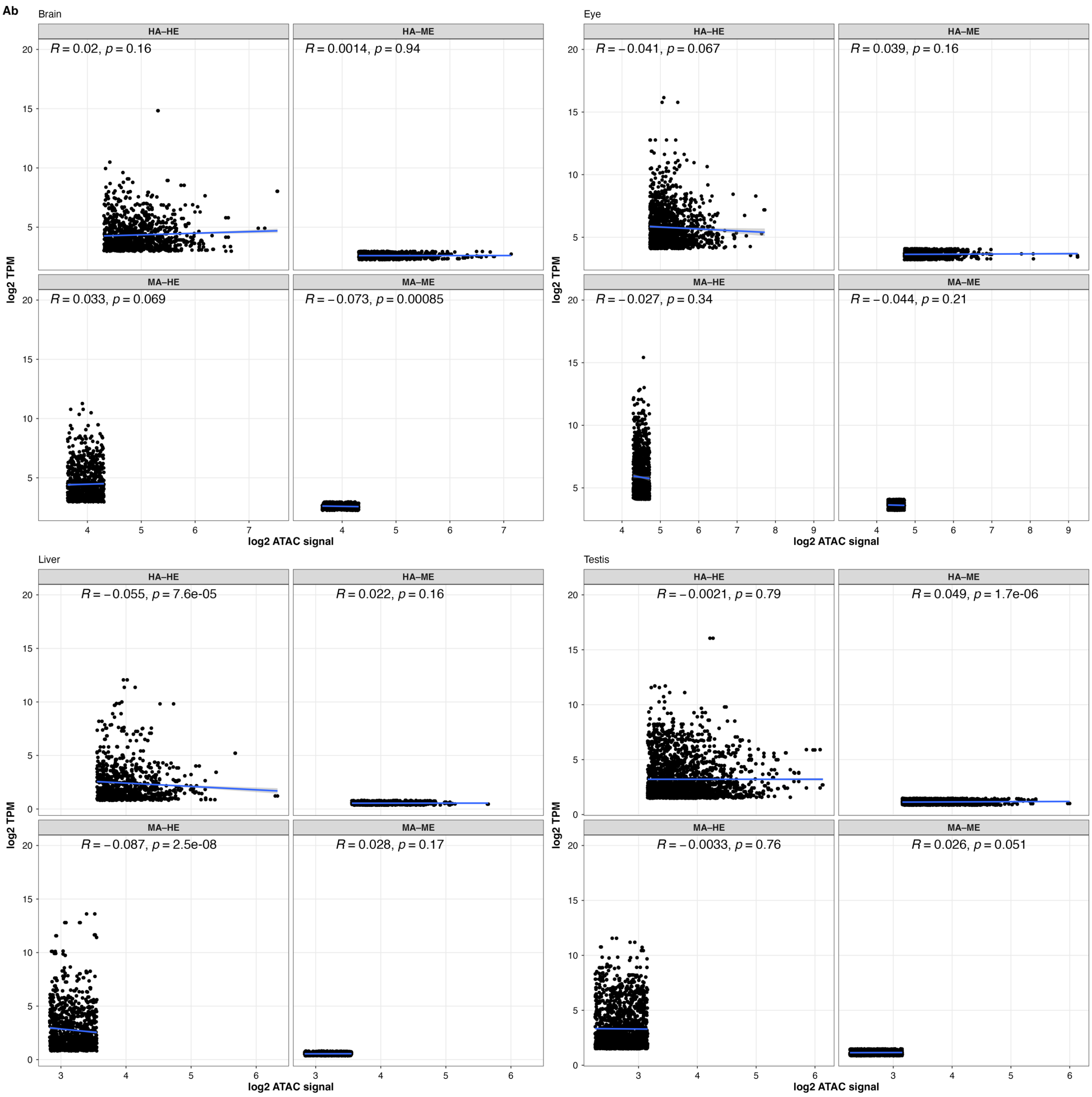


**Fig. S20 – Categorised correlation of accessible peaks accounting for gene expression change in *A. burtoni* tissues.** Tissue-specific average *log2* gene expression as transcripts per million (TPM, y-axis) and average *log2* ATAC signal as gene promoter peak counts (x-axis) for each gene across same tissue biological replicates (Brain = Forebrain and Eye = Retina), split by high accessibility and high expression (HA-HE - both average TPM and ATAC signal values are more than the 70th percentiles), high accessibility and medium-low expression (HA-ME - ATAC signal values more than the 70th percentiles and average TPM less than the 50th

percentile), medium-low accessibility and high expression (MA-HE - ATAC signal values less than the 50th percentiles and average TPM more than the 70th percentile), and medium-low accessibility and medium-low expression (MA-ME - average ATAC signal count less than the 50th percentile and average TPM more than the 70th percentile). Correlation co-efficient (*R,* blue trendline) and *p-value* shown for each category.

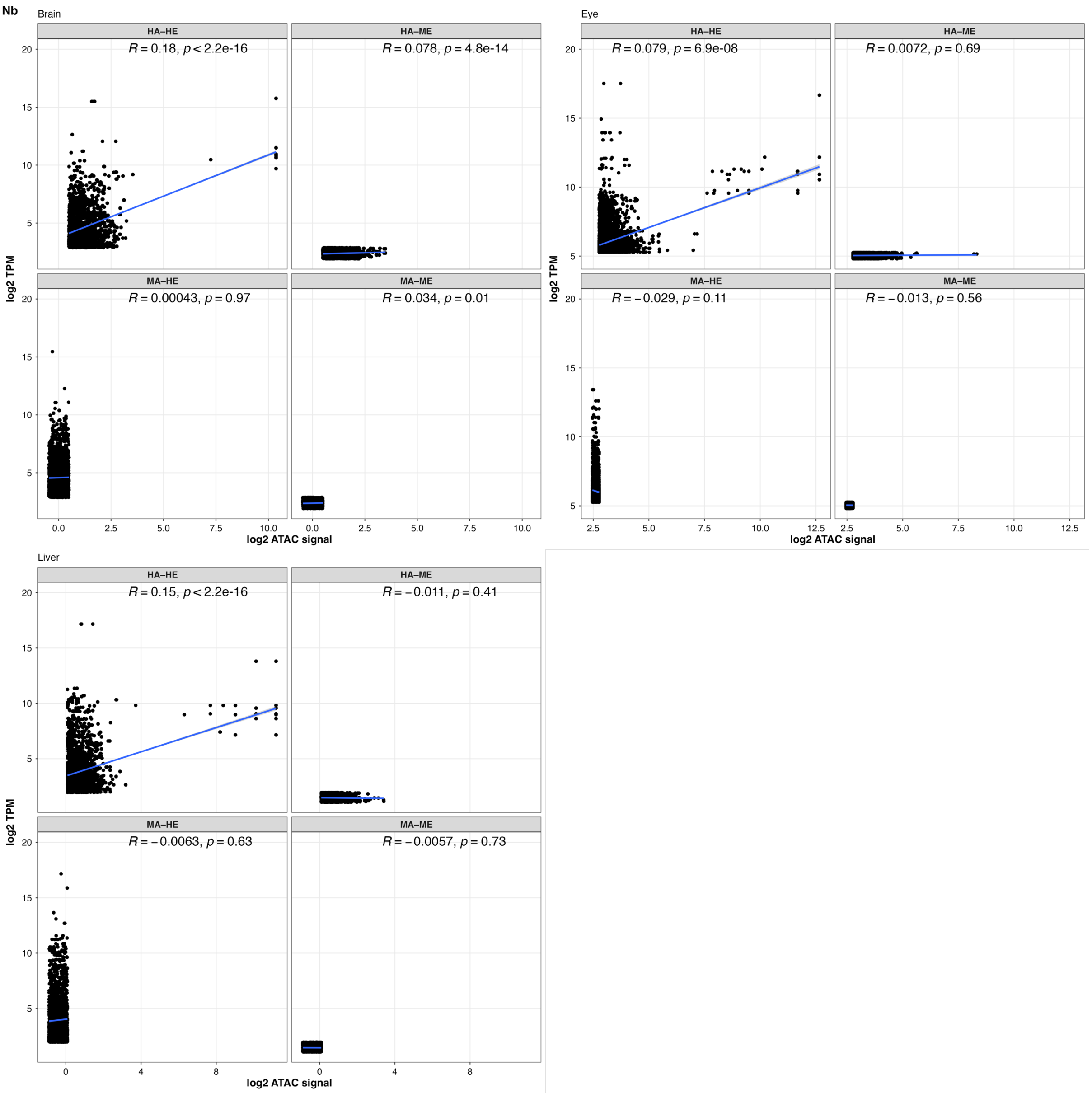


**Fig. S21 – Categorised correlation of accessible peaks accounting for gene expression change in *N. brichardi* tissues.** Tissue-specific average *log2* gene expression as transcripts per million (TPM, y-axis) and average *log2* ATAC signal as gene promoter peak counts (x-axis) for each gene across same tissue biological replicates (Brain = Forebrain and Eye = Retina), split by high accessibility and high expression (HA-HE - both average TPM and ATAC signal values are more than the 70th percentiles), high accessibility and medium-low expression (HA-ME - ATAC signal values more than the 70th percentiles and average TPM less than the 50th

percentile), medium-low accessibility and high expression (MA-HE - ATAC signal values less than the 50th percentiles and average TPM more than the 70th percentile), and medium-low accessibility and medium-low expression (MA-ME - average ATAC signal count less than the 50th percentile and average TPM more than the 70th percentile). Correlation co-efficient (*R,* blue trendline) and *p-value* shown for each category.

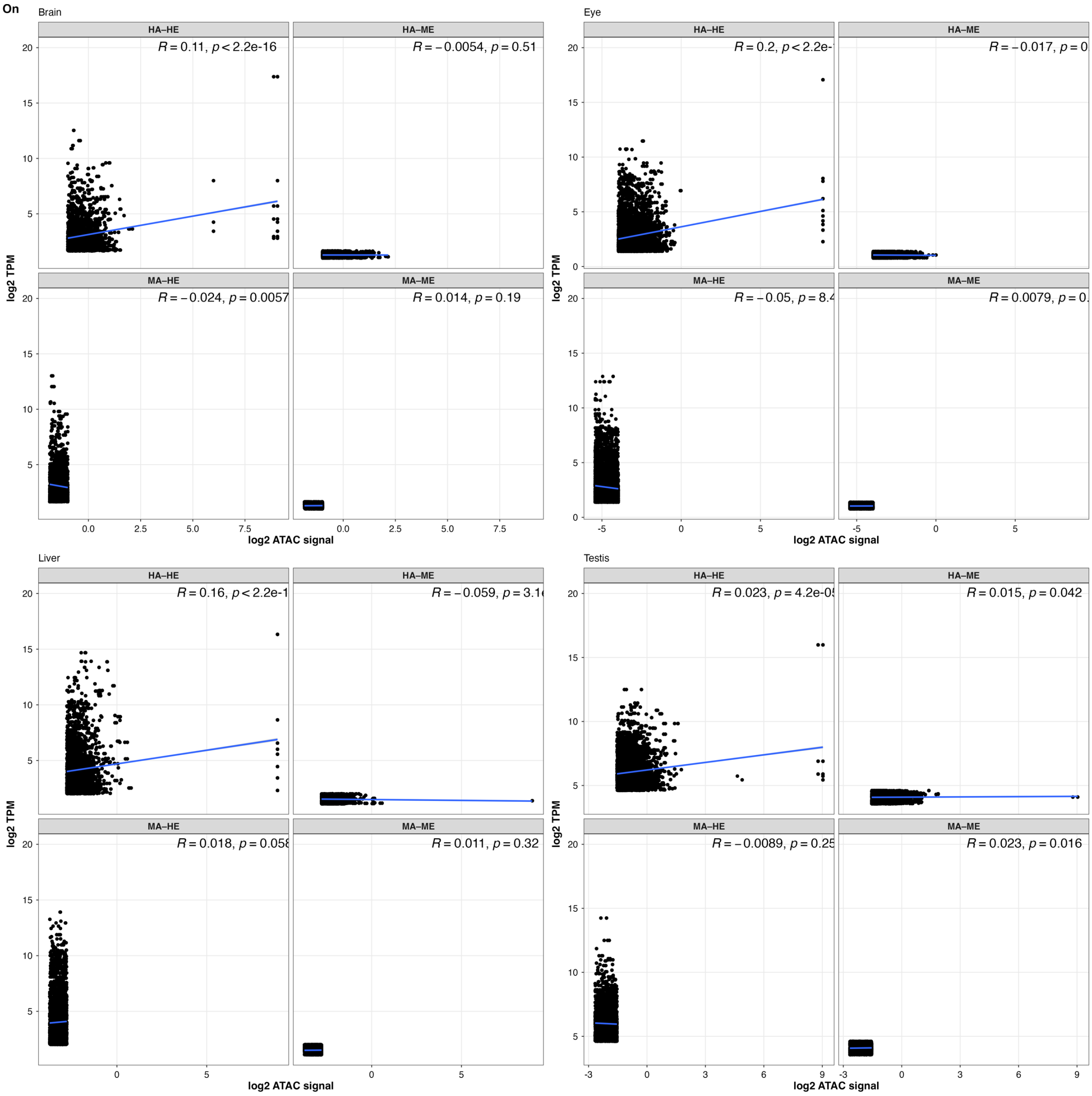


**Fig. S22 – Categorised correlation of accessible peaks accounting for gene expression change in *O. niloticus* tissues.** Tissue-specific average *log2* gene expression as transcripts per million (TPM, y-axis) and average *log2* ATAC signal as gene promoter peak counts (x-axis) for each gene across same tissue biological replicates (Brain = Forebrain and Eye = Retina), split by high accessibility and high expression (HA-HE - both average TPM and ATAC signal values are more than the 70th percentiles), high accessibility and medium-low expression (HA-ME - ATAC signal values more than the 70th percentiles and average TPM less than the 50th

percentile), medium-low accessibility and high expression (MA-HE - ATAC signal values less than the 50th percentiles and average TPM more than the 70th percentile), and medium-low accessibility and medium-low expression (MA-ME - average ATAC signal count less than the 50th percentile and average TPM more than the 70th percentile). Correlation co-efficient (*R,* blue trendline) and *p-value* shown for each category.

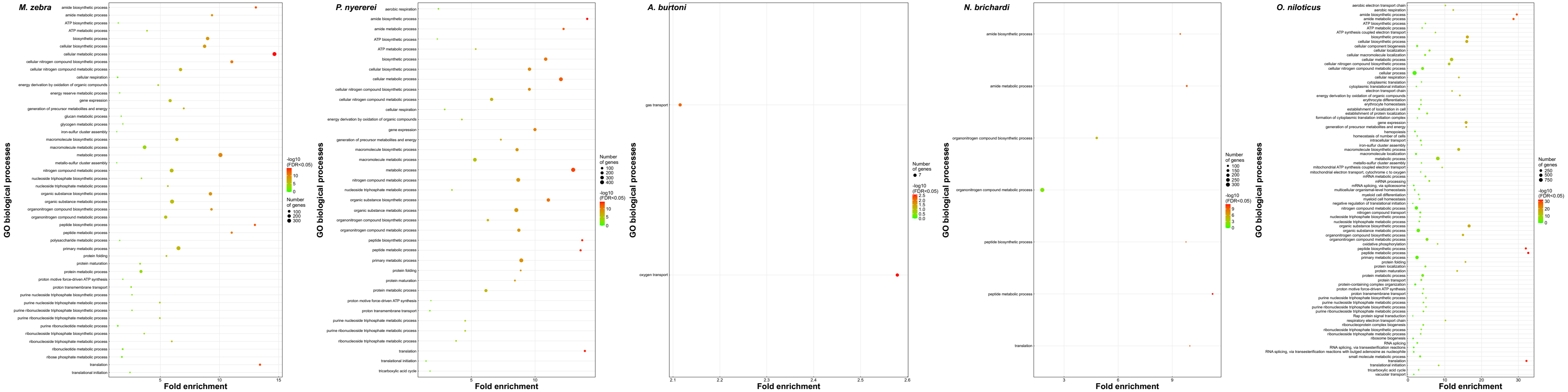


**Fig. S23 – Gene Ontology (GO) enrichment of genes categorised as high accessibility (HA) and high expression (HE) in retina tissue of five cichlid species.** GO biological processes enrichment of HA-HE genes in retina tissue of five cichlids. Circles show enriched biological processes (y-axis) of significance ($-log_{10}$ FDR <0.05, heatmap to right) and $-log_{10}$ FDR <0.05 (x-axis) values of each term across each tissue. Number of enriched genes for each term shown by size of each circle.

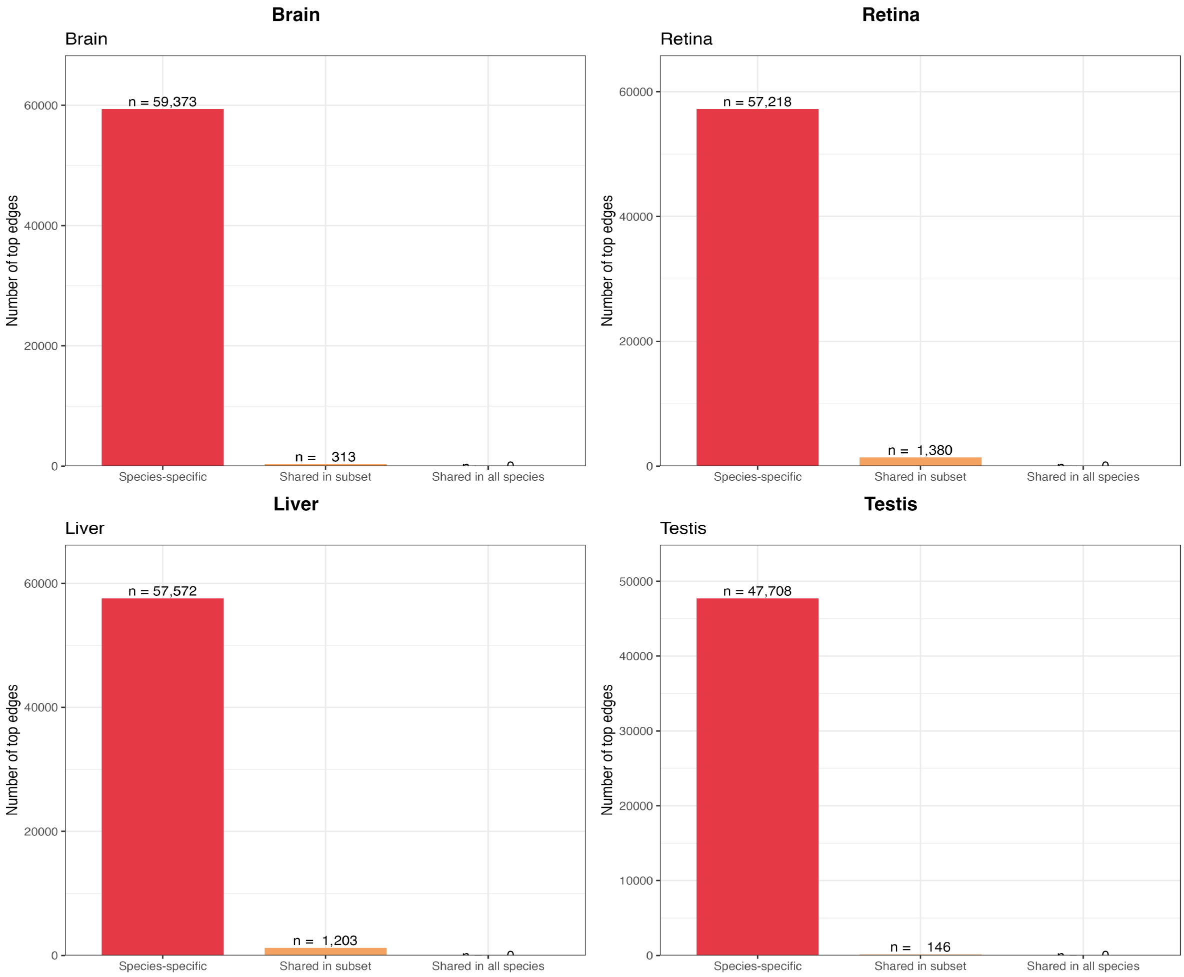


**Fig. S24 – Conservation or divergence of top 12,000 expression only inferred GRN edges across tissues.** Distribution of top 12k GENIE3 edges classified as species-specific, shared in a subset of species, or shared in all five species for forebrain, retina, liver, and testis. Counts are shown on the bars and where absent, are zero-count categories.

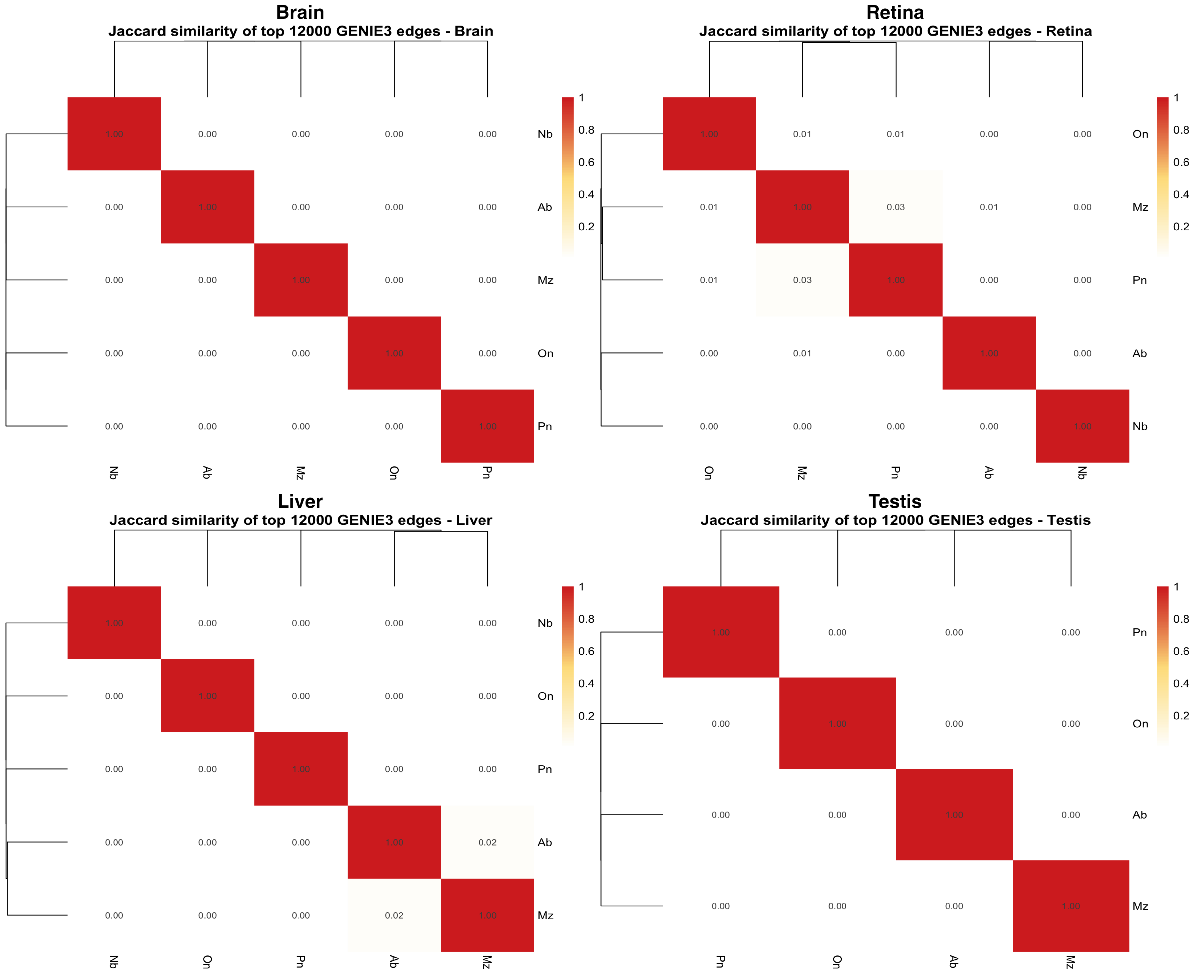


**Fig. S25 - Jaccard similarity of the top 12,000 GENIE3 edges across species in forebrain, retina, liver, and testis.** Heatmaps show pairwise Jaccard similarity among the top 12,000 GENIE3 edges for each tissue, with panels for forebrain, retina, liver, and testis. Species are ordered by hierarchical clustering on both axes, and the same species labels are shown on the x- and y-axes in each panel. Cell values are displayed within the matrix and range from 0 to 1, with darker red indicating higher similarity and pale/white shading indicating lower similarity.

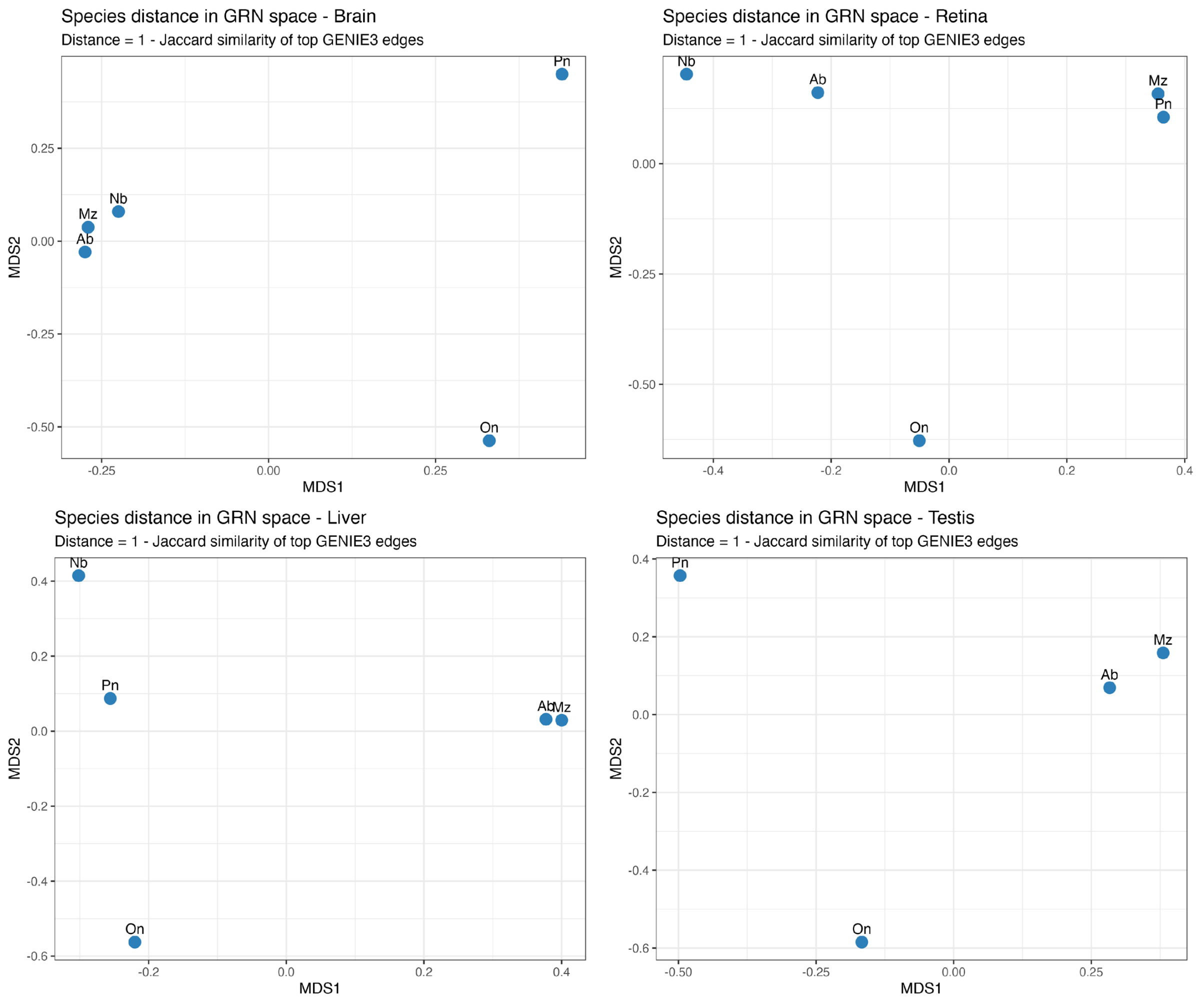


**Fig. S26 - Species distance in GENIE3 edge GRN space across tissues.**

MDS plots of species distances in gene regulatory network (GRN) space for forebrain, retina, liver, and testis, where distance is defined as 1 minus Jaccard similarity of top GENIE3 edges. Points are coloured blue and labelled by species (Mz, Pn, Ab, Nb, On), plotted against MDS1 and MDS2 dimensions.

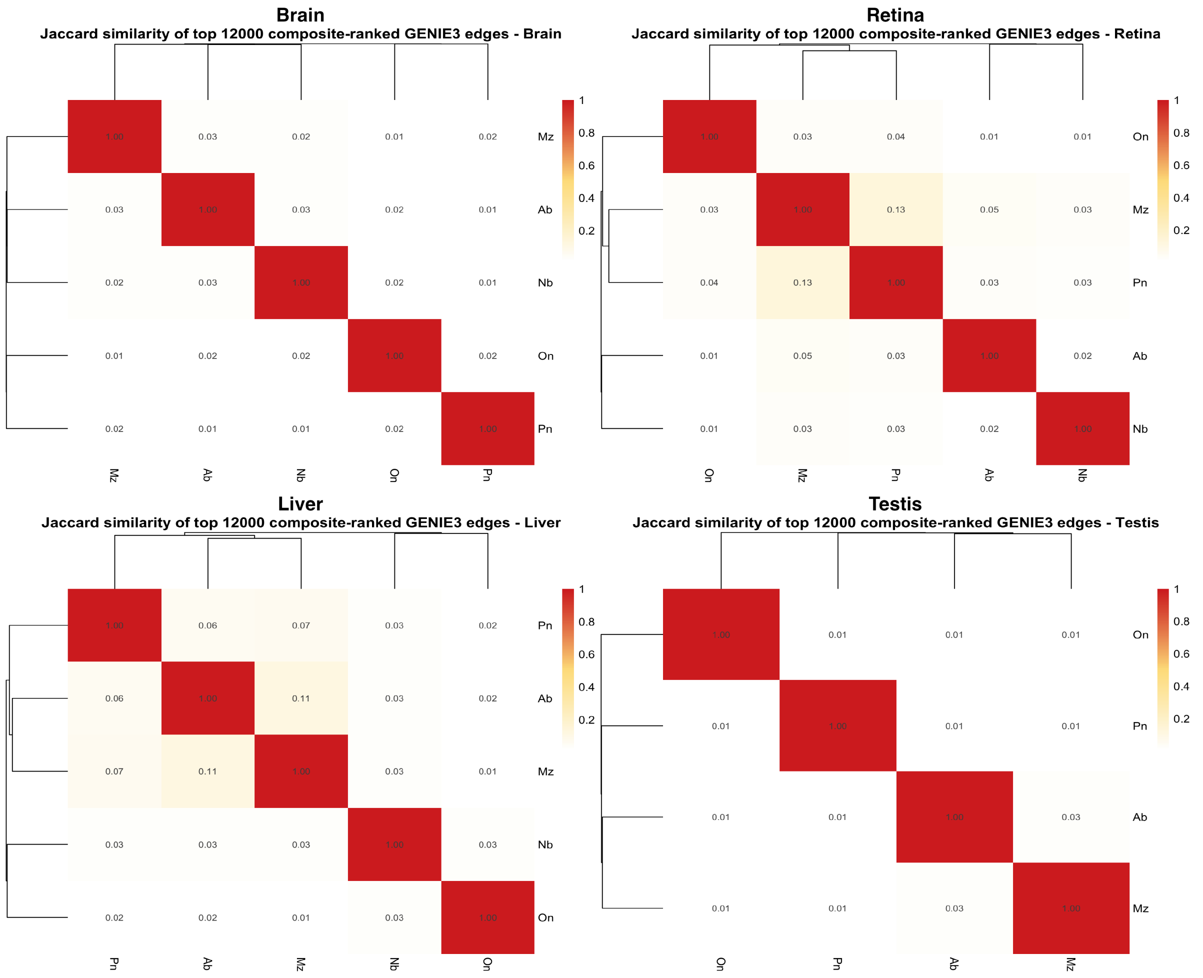


**Fig. S27 - Jaccard similarity of top composite-ranked (motif-support integrated) GENIE3 edges across species.** Heatmaps show pairwise Jaccard similarity among the top 12,000 composite-ranked (integrating motif-support) GENIE3 edges for forebrain, retina, liver, and testis (as separate panels). Species are ordered by hierarchical clustering on both axes, with identical species labels (Mz, Pn, Ab, Nb, On) on x- and y-axes per panel. Cell values are shown in the matrix, ranging from 0 to 1, with darker red for higher similarity and lighter shades for lower similarity.

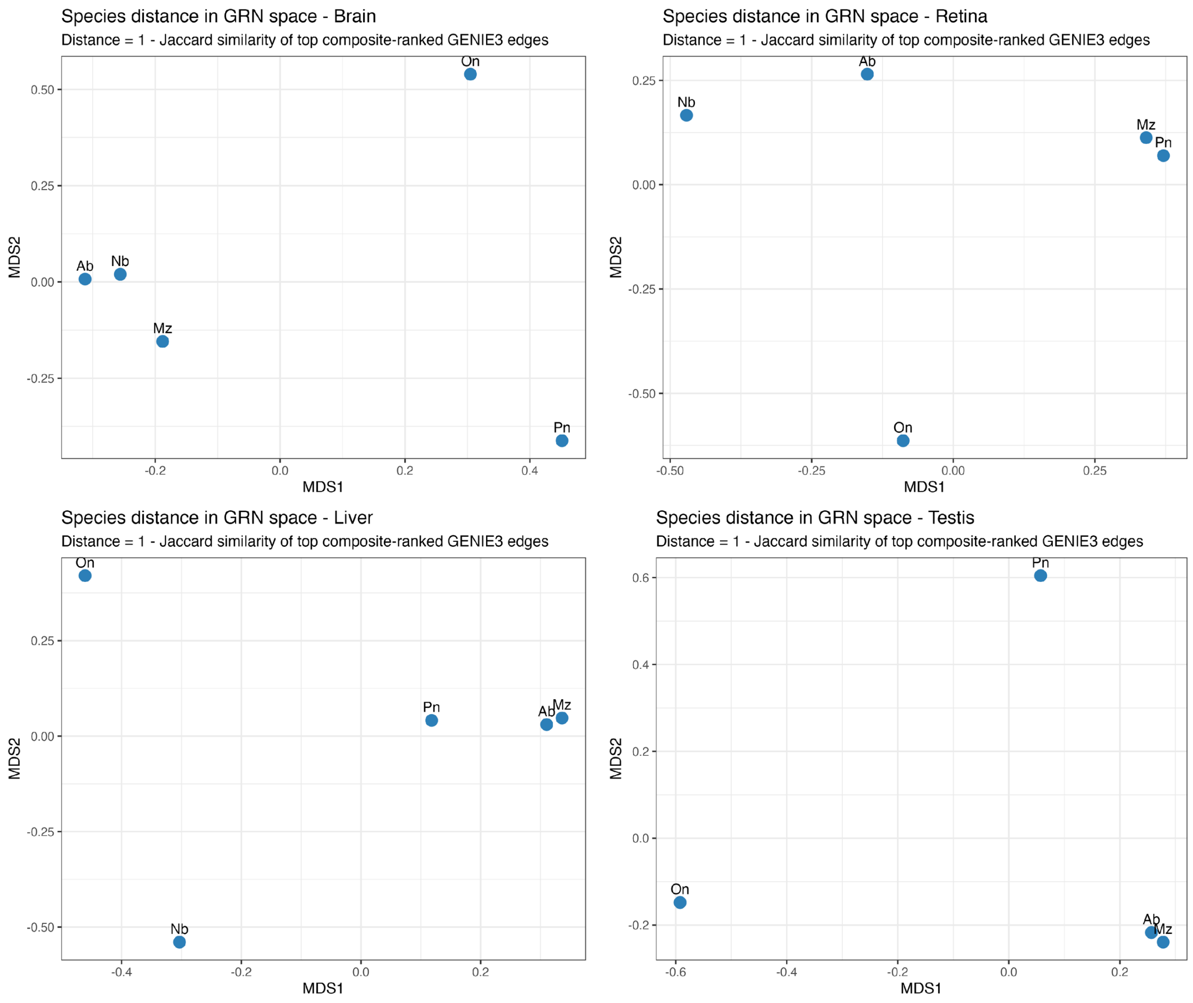


**Fig. S28 - Species distance in composite-ranked (motif-support integrated) GENIE3 edge GRN space across tissues.**

MDS plots of species distances in gene regulatory network (GRN) space for forebrain, retina, liver, and testis, where distance is defined as 1 minus Jaccard similarity of top composite-ranked (motif-support integrated) GENIE3 edges. Points are coloured blue and labelled by species (Mz, Pn, Ab, Nb, On), plotted against MDS1 and MDS2 dimensions.

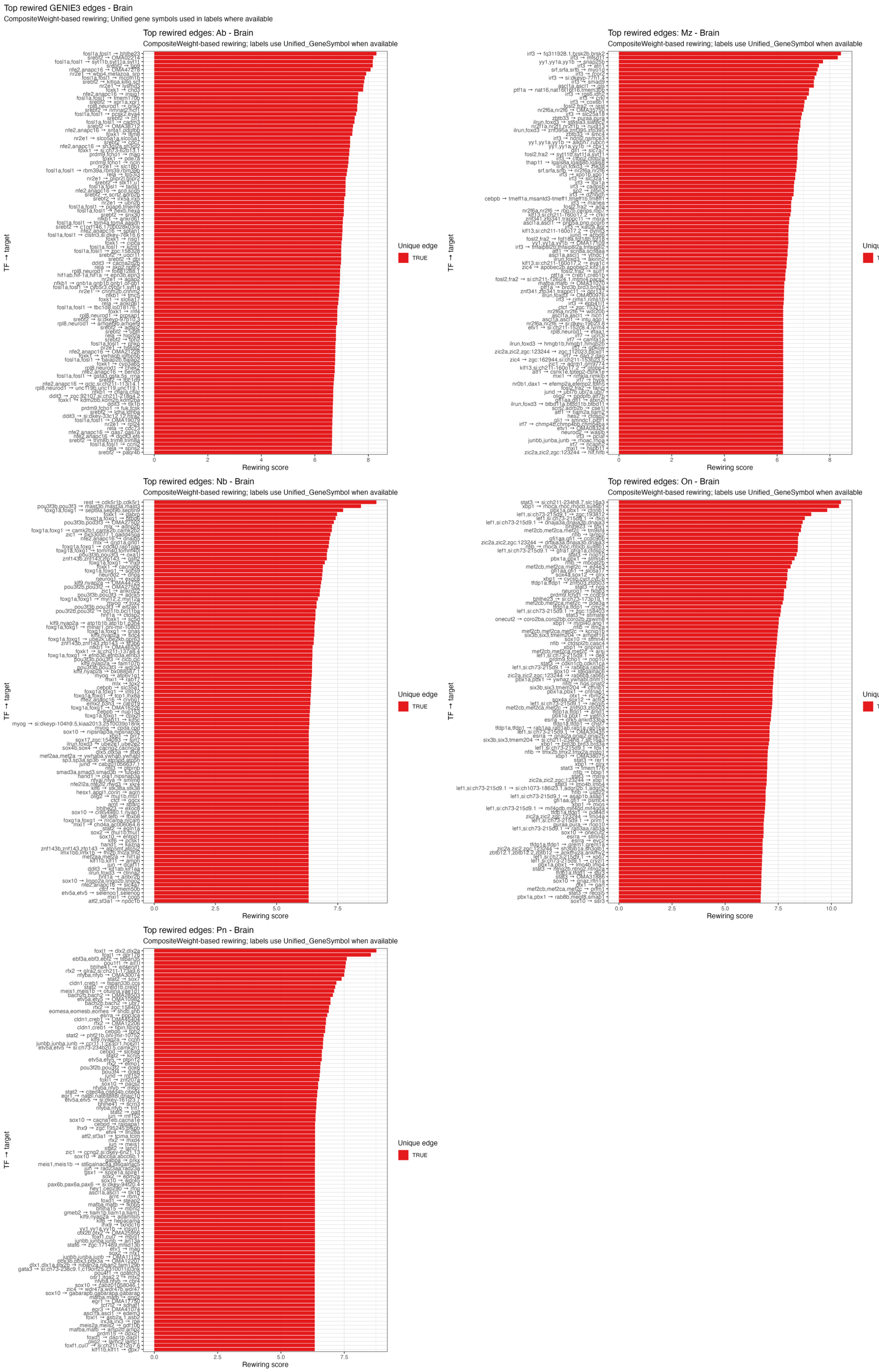

Top rewired GENIE3 edges - Brain
CompositeWeight-based rewiring; Unified gene symbols used in labels where available
Top rewired edges: Ab - Brain
CompositeWeight-based rewiring; labels use Unified_GeneSymbol when available
Top rewired edges: Mz - Brain
CompositeWeight-based rewiring; labels use Unified_GeneSymbol when available
Top rewired edges: Nb - Brain
CompositeWeight-based rewiring; labels use Unified_GeneSymbol when available
Top rewired edges: On - Brain
CompositeWeight-based rewiring; labels use Unified_GeneSymbol when available
Top rewired edges: Pn - Brain
CompositeWeight-based rewiring; labels use Unified_GeneSymbol when available
TF → target
Rewiring score
Unique edge
TRUE

**Fig. S29 - Top rewired GENIE3 edges in forebrain across species.** Bar plots ranking the top composite-weighted rewired GENIE3 edges in brain for each of the five species. Bars are coloured red and show rewiring score on the x-axis; gene symbols (unified where available) label the y-axis. Red horizontal lines indicate edges confirmed as unique in motif-supported GENIE3 edges.

Top rewired GENIE3 edges - Retina

CompositeWeight-based rewiring; Unified gene symbols used in labels where available

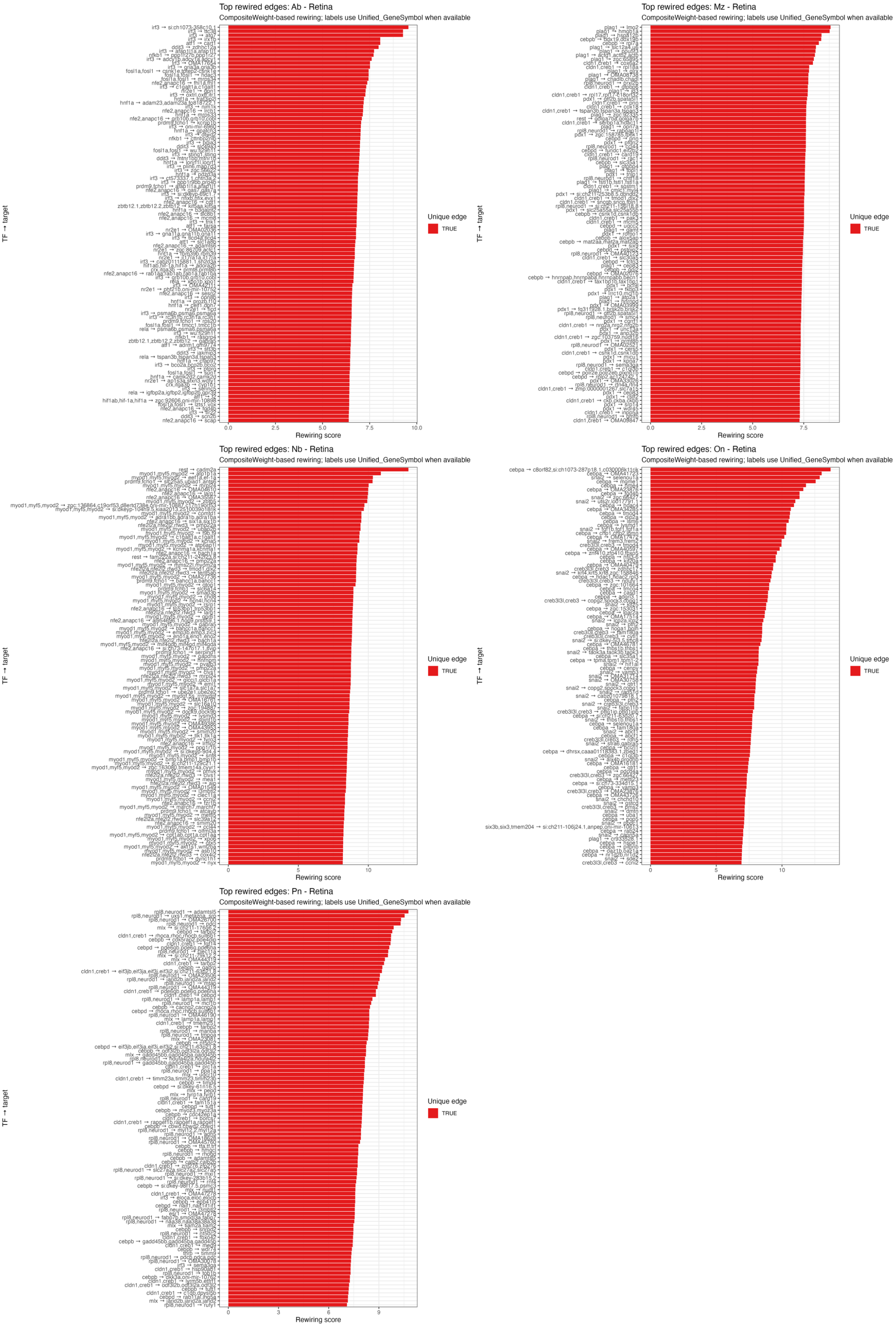

**Fig. S30 - Top rewired GENIE3 edges in retina across species.** Bar plots ranking the top composite-weighted rewired GENIE3 edges in retina for each of the five species. Bars are coloured red and show rewiring score on the x-axis; gene symbols (unified where available) label the y-axis. Red horizontal lines indicate edges confirmed as unique in motif-supported GENIE3 edges.

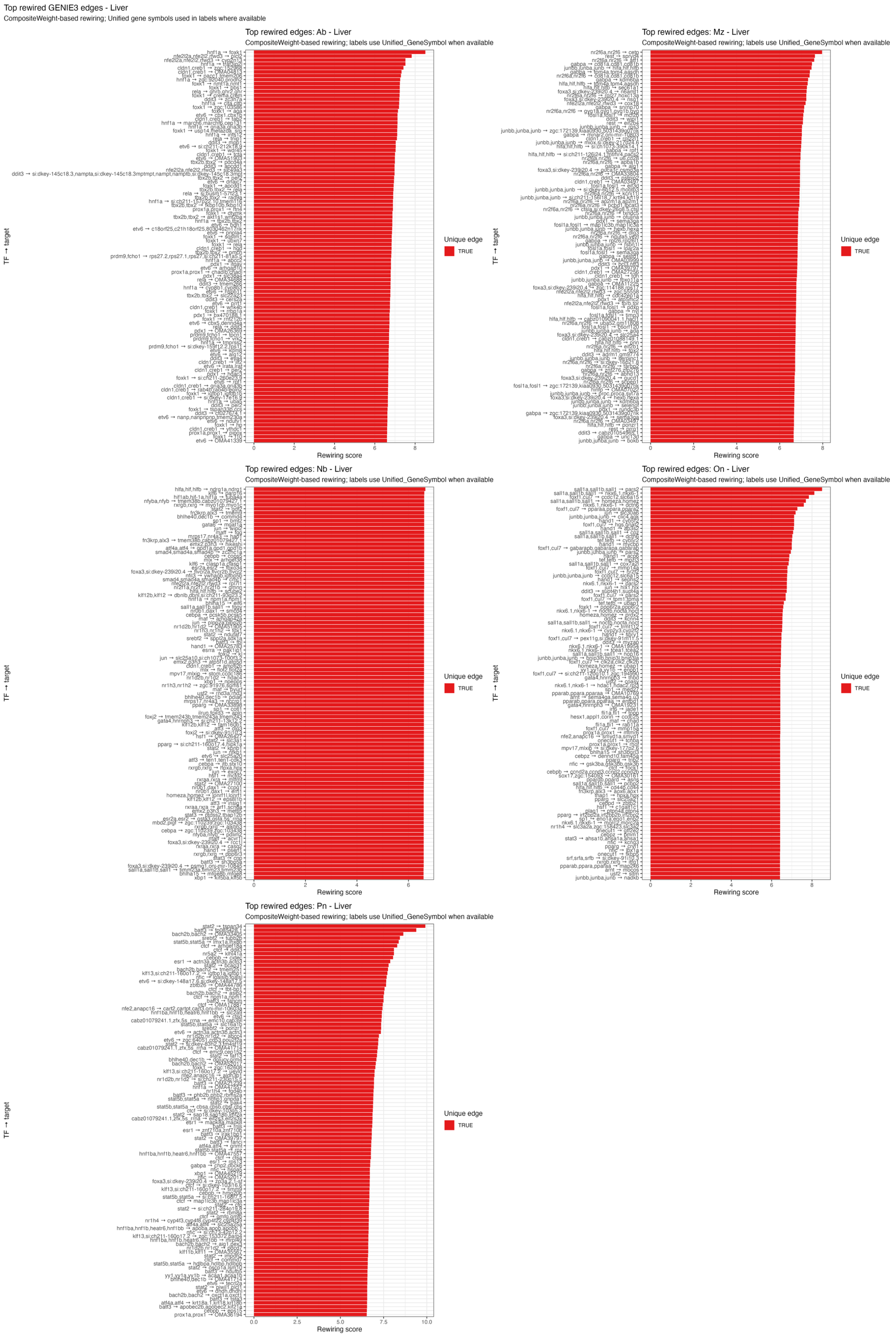

Top rewired GENIE3 edges - Liver
CompositeWeight-based rewiring; Unified gene symbols used in labels where available
Top rewired edges: Ab - Liver
CompositeWeight-based rewiring; labels use Unified_GeneSymbol when available
Top rewired edges: Mz - Liver
CompositeWeight-based rewiring; labels use Unified_GeneSymbol when available
Top rewired edges: Nb - Liver
CompositeWeight-based rewiring; labels use Unified_GeneSymbol when available
Top rewired edges: On - Liver
CompositeWeight-based rewiring; labels use Unified_GeneSymbol when available
Top rewired edges: Pn - Liver
CompositeWeight-based rewiring; labels use Unified_GeneSymbol when available
TF → target
Rewiring score
Unique edge
TRUE

**Fig. S31 - Top rewired GENIE3 edges in liver across species.** Bar plots ranking the top composite-weighted rewired GENIE3 edges in liver for each of the five species. Bars are coloured red and show rewiring score on the x-axis; gene symbols (unified where available) label the y-axis. Red horizontal lines indicate edges confirmed as unique in motif-supported GENIE3 edges.

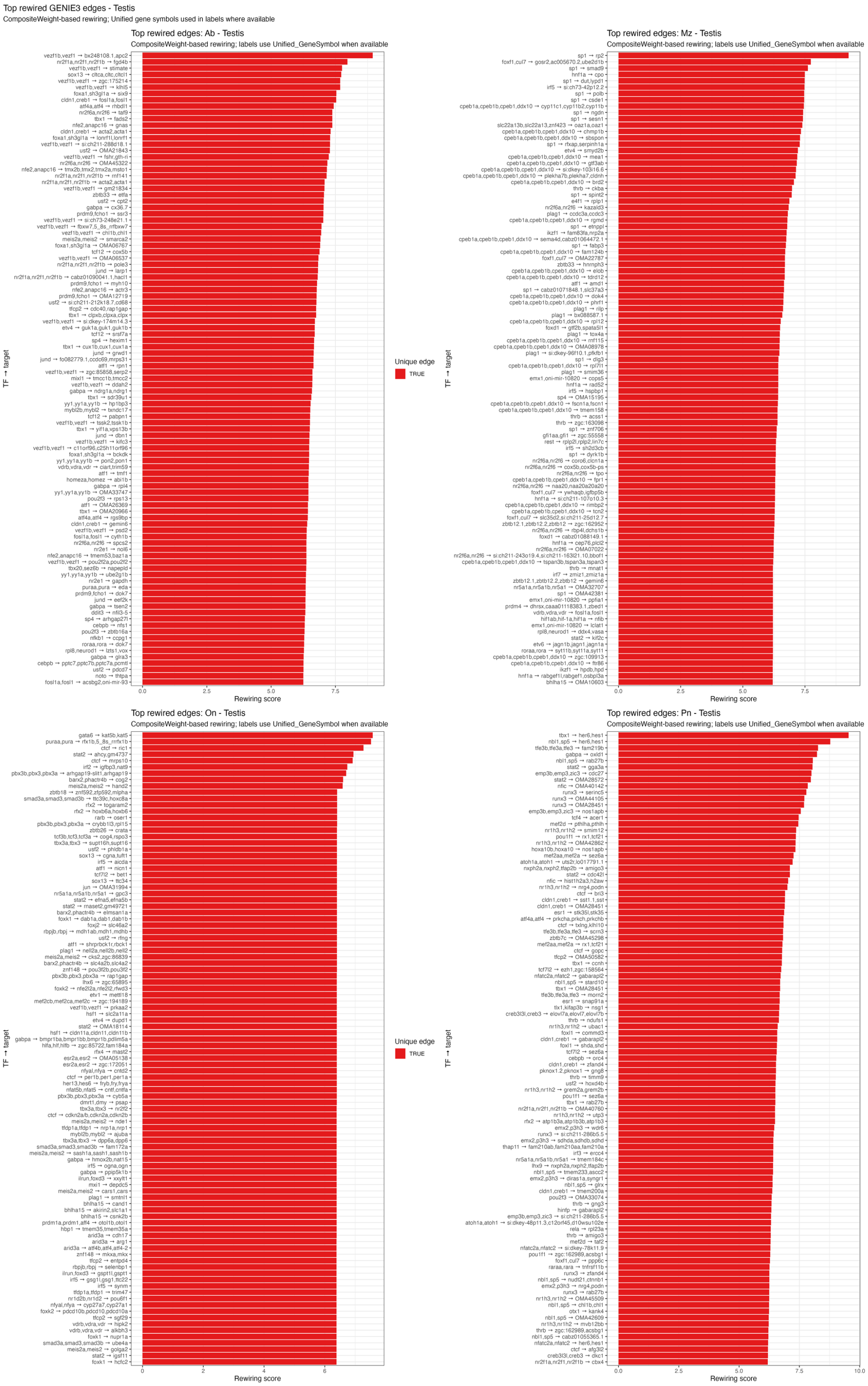
Top rewired GENIE3 edges - Testis
CompositeWeight-based rewiring; Unified gene symbols used in labels where available
Top rewired edges: Ab - Testis
CompositeWeight-based rewiring; labels use Unified_GeneSymbol when available
Top rewired edges: Mz - Testis
CompositeWeight-based rewiring; labels use Unified_GeneSymbol when available
Top rewired edges: On - Testis
CompositeWeight-based rewiring; labels use Unified_GeneSymbol when available
Top rewired edges: Pn - Testis
CompositeWeight-based rewiring; labels use Unified_GeneSymbol when available
TF → target
Rewiring score
Unique edge
TRUE

**Fig. S32 - Top rewired GENIE3 edges in testis across species.** Bar plots ranking the top composite-weighted rewired GENIE3 edges in testis for of four species. Bars are coloured red and show rewiring score on the x-axis; gene symbols (unified where available) label the y-axis. Red horizontal lines indicate edges confirmed as unique in motif-supported GENIE3 edges.

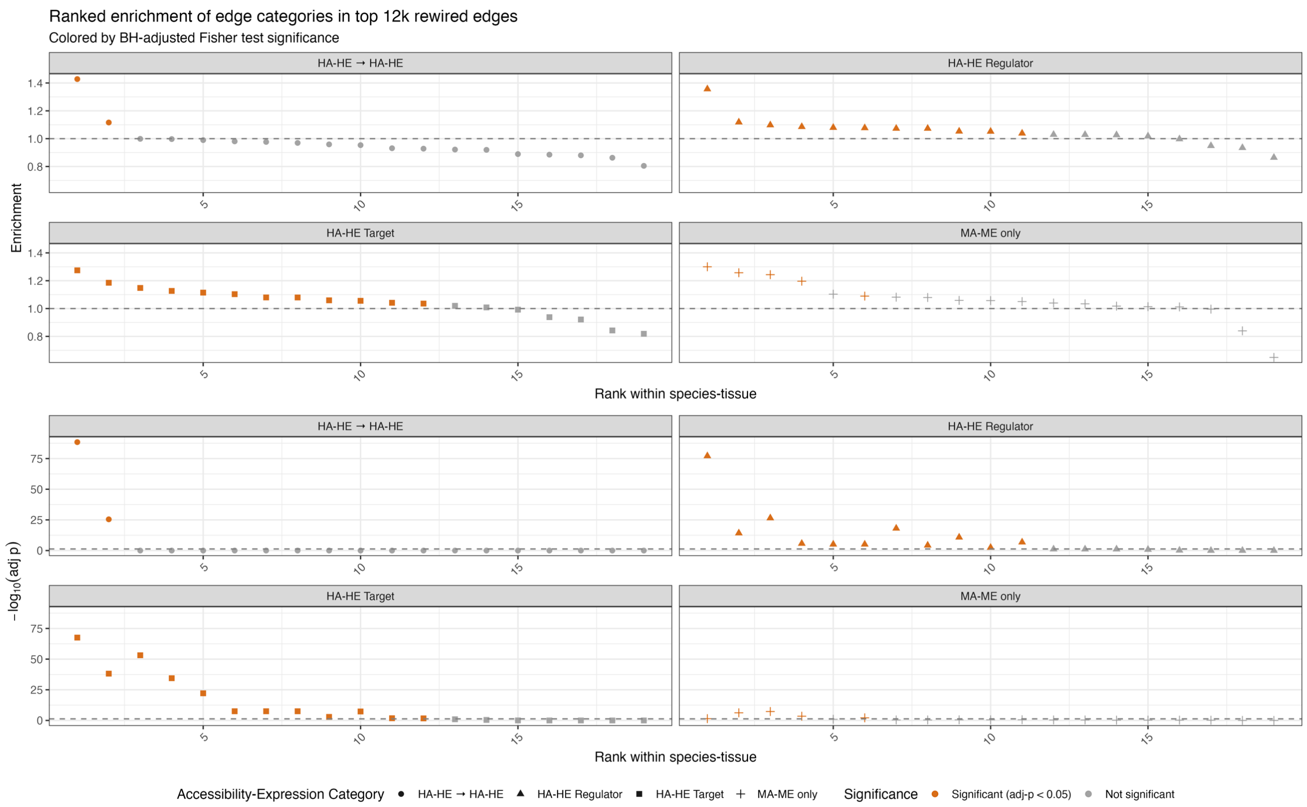


**Fig. S33 – Collated ranked enrichment of HA/HE categories in rewired edges.** Line plots show BH-adjusted Fisher test enrichment of accessibility-expression categories (HA-HE → HA+HE, HA-HE regulator/target, MA-ME only) among the top 12K rewired edges, ranked by enrichment rank within species-tissue. Points are coloured by significance (orange for adj. p<0.05, grey for non-significant) with shapes distinguishing categories (circles for HA-HE → HA+HE, triangles for HA-HE as regulators, square as targets, and cross for MA-ME only). Separate panels for enrichment and their -$\log_{10}$(p-adj).

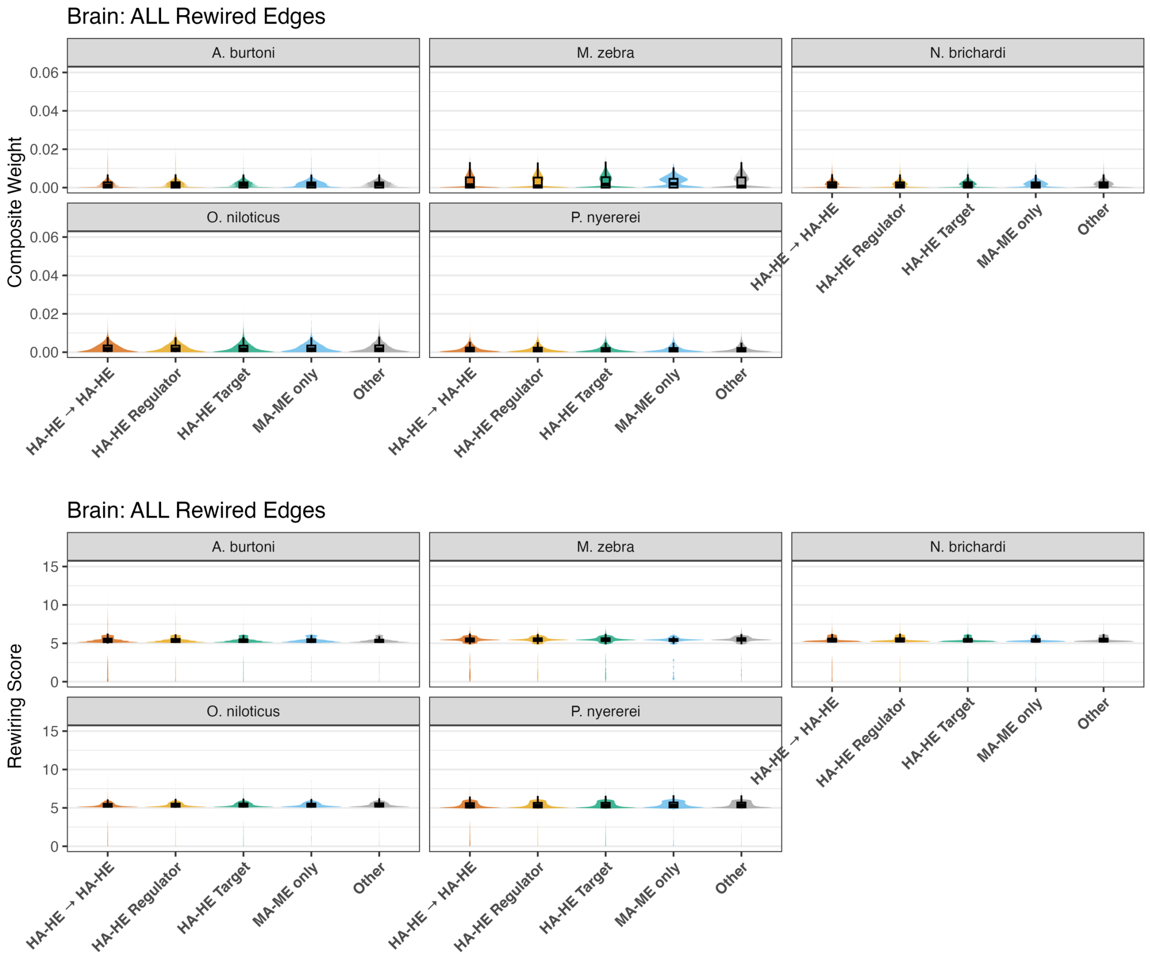


**Fig. S34 – Accessibility-Expression category distribution in all rewired edges for forebrain across species.** Top row: violin/box plots of composite weight (y-axis) for rewired edges by all accessibility-expression categories (x-axis: HA-HE → HA+HE, HA-HE regulator/target, MA-ME, other) across all five species. Bottom row: violin/box plots of rewiring scores (y-axis) for the same categories and species. All categories coloured distinctly (HA-HE → HA+HE: orange; HA-HE regulator: yellow; HA-HE target: green; MA-ME: blue; other: grey).

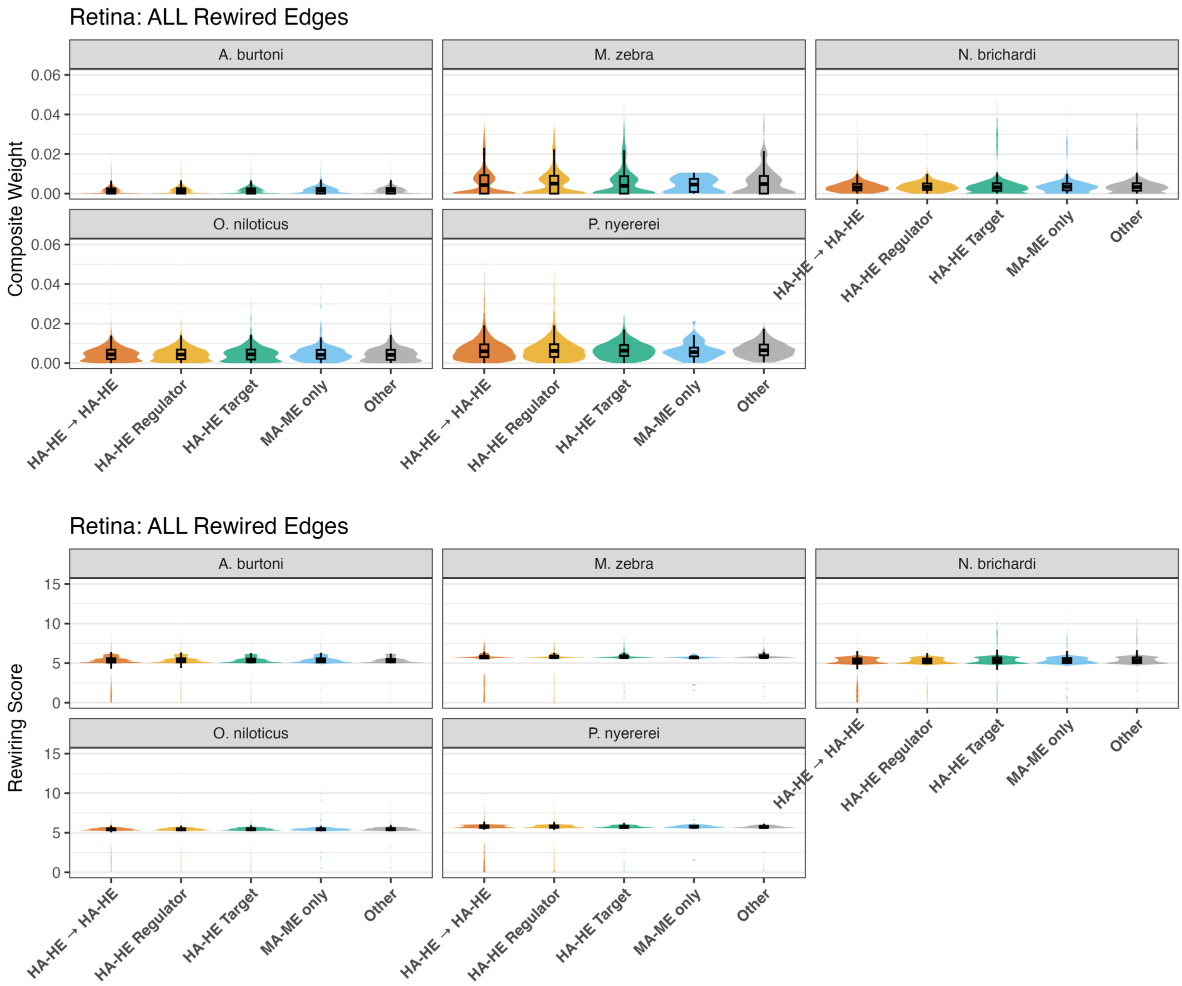


**Fig. S35 – Accessibility-Expression category distribution in all rewired edges for retina across species.** Top row: violin/box plots of composite weight (y-axis) for rewired edges by all accessibility-expression categories (x-axis: HA-HE → HA+HE, HA-HE regulator/target, MA-ME, other) across all five species. Bottom row: violin/box plots of rewiring scores (y-axis) for the same categories and species. All categories coloured distinctly (HA-HE → HA+HE: orange; HA-HE regulator: yellow; HA-HE target: green; MA-ME: blue; other: grey).

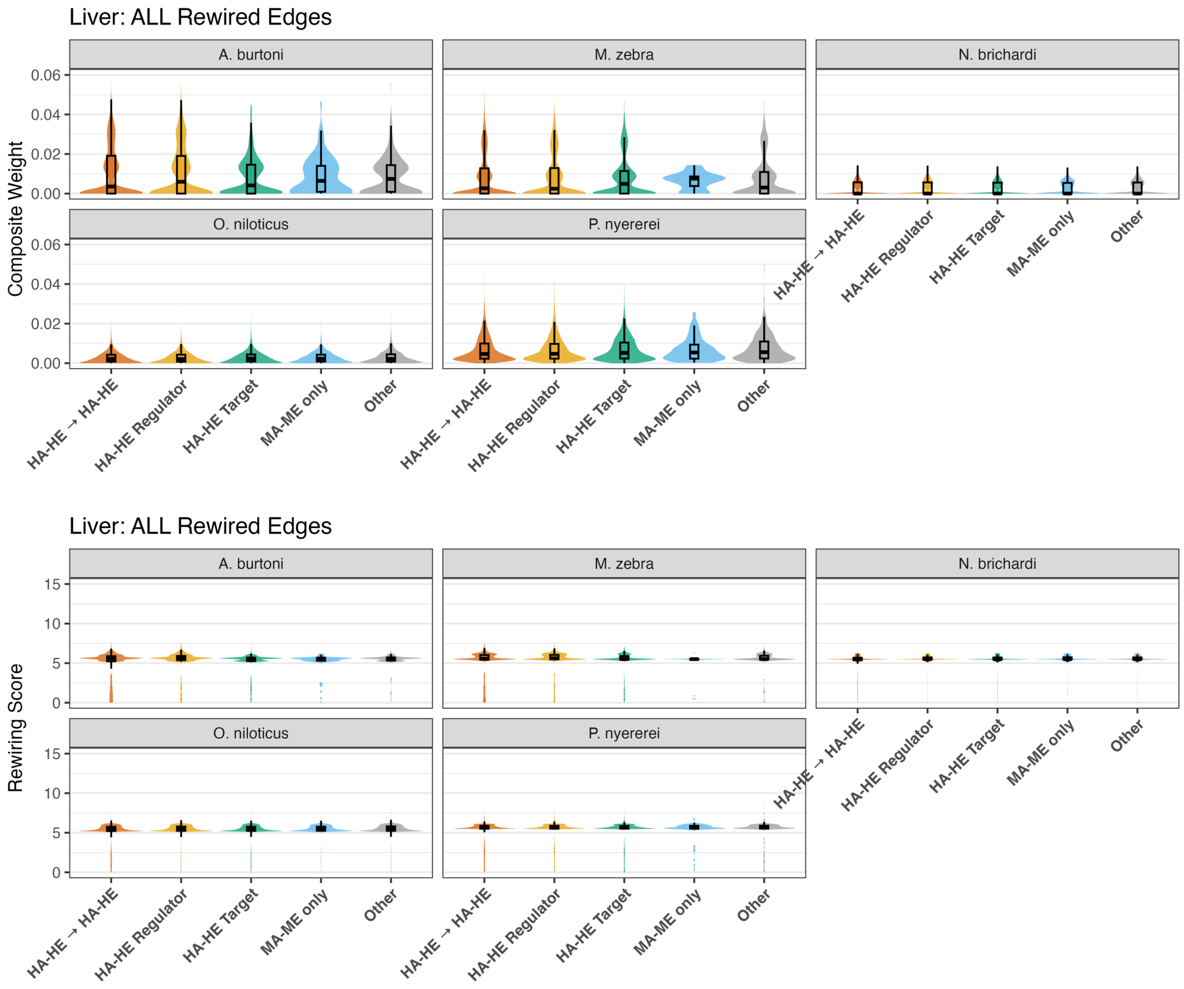


**Fig. S36 – Accessibility-Expression category distribution in all rewired edges for liver across species.** Top row: violin/box plots of composite weight (y-axis) for rewired edges by all accessibility-expression categories (x-axis: HA-HE → HA+HE, HA-HE regulator/target, MA-ME, other) across all five species. Bottom row: violin/box plots of rewiring scores (y-axis) for the same categories and species. All categories coloured distinctly (HA-HE → HA+HE: orange; HA-HE regulator: yellow; HA-HE target: green; MA-ME: blue; other: grey).

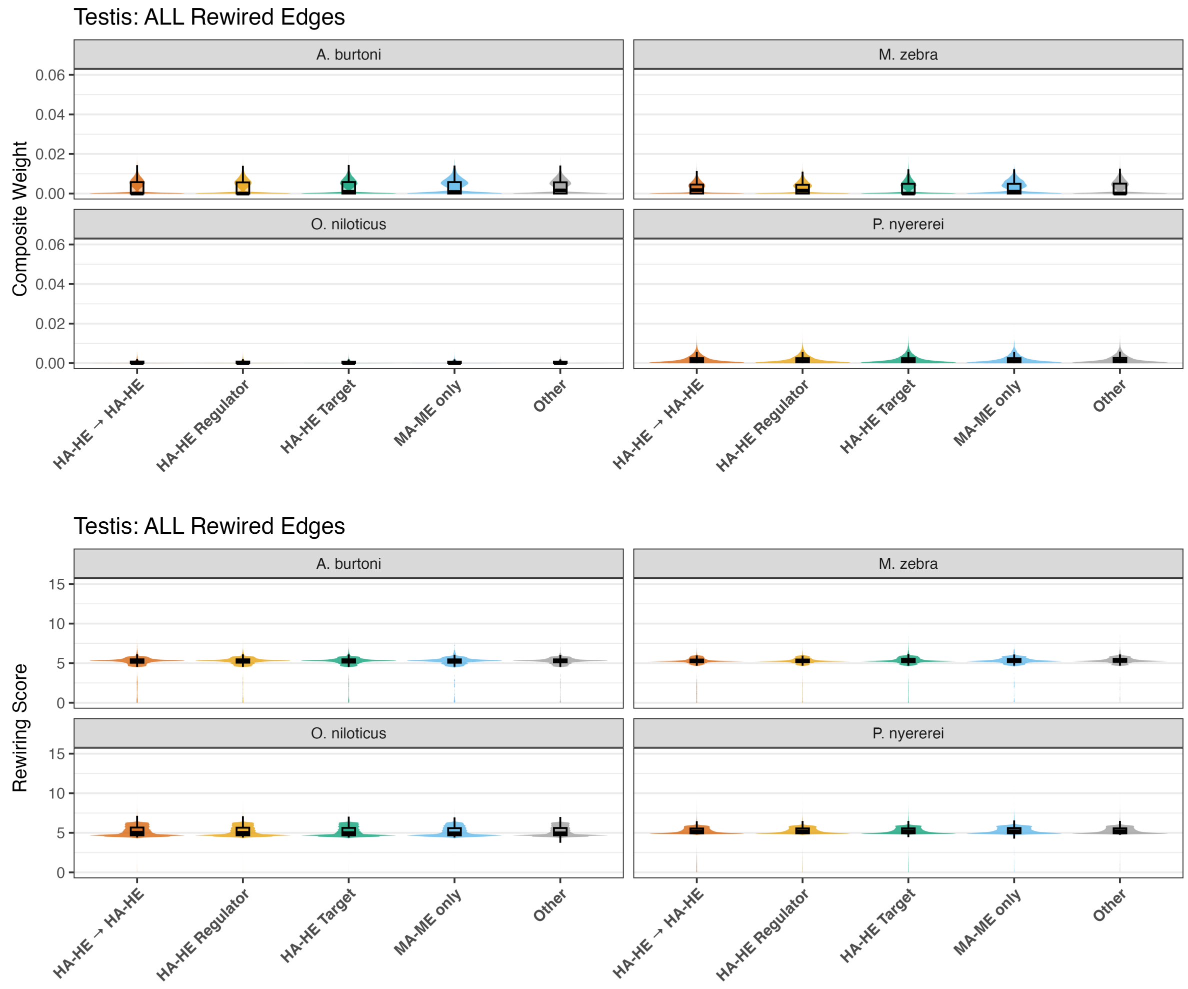


**Fig. S37 – Accessibility-Expression category distribution in all rewired edges for testis across species.** Top row: violin/box plots of composite weight (y-axis) for rewired edges by all accessibility-expression categories (x-axis: HA-HE → HA+HE, HA-HE regulator/target, MA-ME, other) across four species. Bottom row: violin/box plots of rewiring scores (y-axis) for the same categories and species. All categories coloured distinctly (HA-HE → HA+HE: orange; HA-HE regulator: yellow; HA-HE target: green; MA-ME: blue; other: grey).

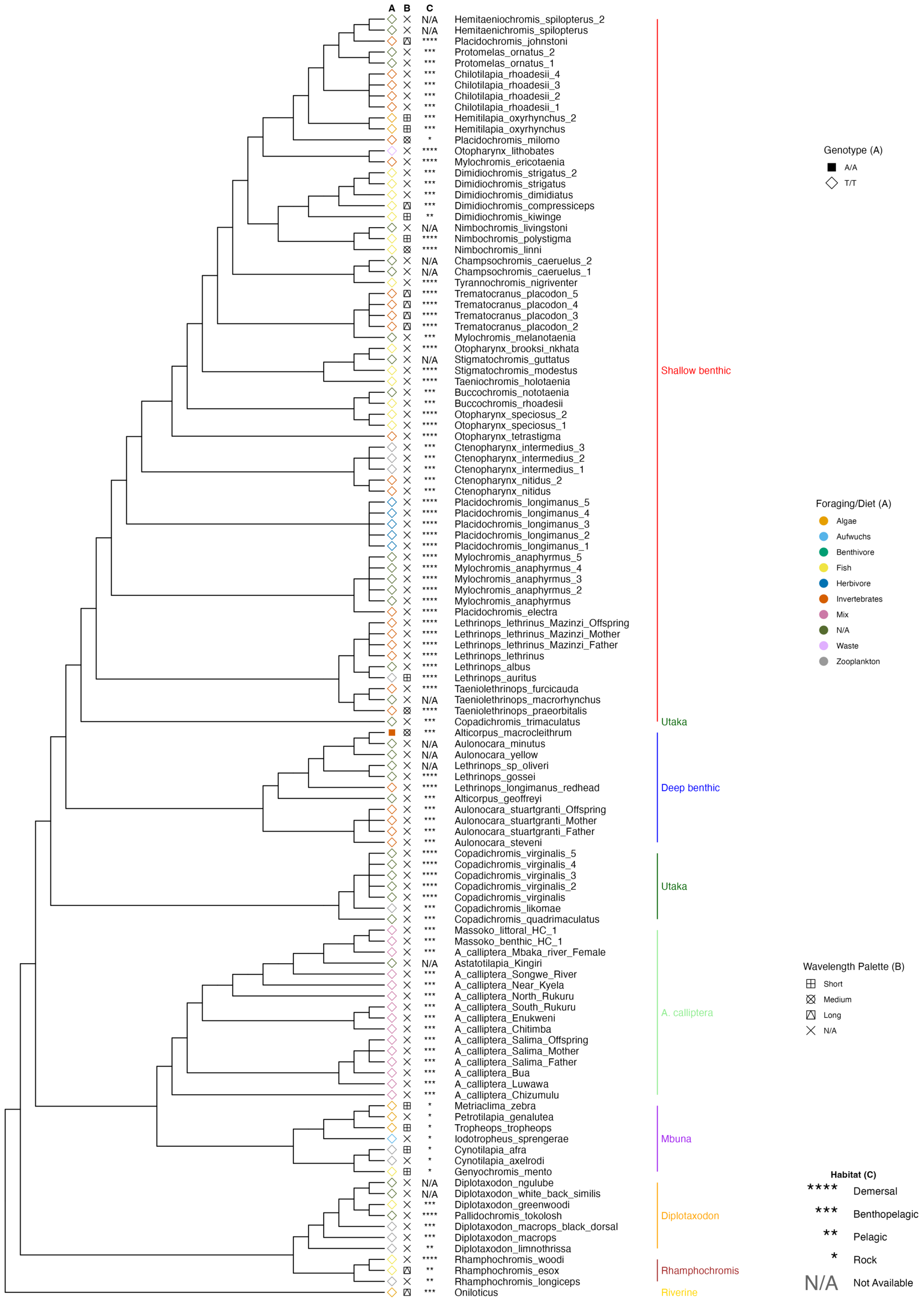


**Fig. S38 – SNP genotypes overlapping MAF TFBS in *M. zebra actr1* promoter and other Lake Malawi species.** Lake Malawi phylogeny reproduced from published least controversial and all included species ASTRAL phylogeny [2], including *O. niloticus* as an outgroup. Phylogenetic branches labelled with species sample name and clade according to legends (right): (a) species foraging/diet habit (color) [3] and phased SNP genotype (shape) [2,4]; (b) adult opsin wavelength palette utilised [3] and (c) species habitat [3,5].

```
                                              ....:..970....:..980....:..990....:.1000....:.1010....:.1020
O. niloticus actr1 gene promoter          180 ATTATTAGATAAATGTGATAGTTAACTGTGATCTGCCAGCACACCAGCAGAAACGCTGCC
R. 'chilingali' actr1 gene promoter       945 ATTATTTGTTAAATGTGATAGTTAACTGTGATCTGCCAGCAGACCAGCAGAAACGCTGCC
T. 'mauve' actr1 gene promoter            946 ATTATTTGTTAAATGTGATAGTTAACTGTGATCTGCCAGCAGACCAGCAGAAACGCTGCC
A. stuartgranti actr1 gene promoter       837 ATTATTTGTTAAATGTGATAGTTAACTGTGATCTGCCAGCAGACCAGCAGAAACGCTGCC
M. zebra actr1 gene promoter              945 ATTATTTGTTAAATGTGATAGTTAACTGTGATCTGCCAGCAGACCAGCAGAAACGCTGCC
A. calliptera actr1 gene promoter         944 ATTATTTGTTAAATGTGATAGTTAACTGTGATCTGCCAGCAGACCAGCAGAAACGCTGCC
A. macrocleithrum actr1 gene promoter       1 ---------------------------------------------------------GCC

                                              ....:.1030....:.1040....:.1050....:.1060....:.1070....:.1080
O. niloticus actr1 gene promoter          240 GCTCGTTGACAGCGTACAGACATGCTGATTAGCTCTCCGTGCTGCCAGTCAGGATCTCTG
R. 'chilingali' actr1 gene promoter      1005 ACTCGTTGACAGCGTATAGACATGCTGATTGGCTCTCCGTGATGCCAGTCAGGATCTCTG
T. 'mauve' actr1 gene promoter           1006 ACTCGTTGACAGCGTATAGACATGCTGATTGGCTCTCCGTGATGCCAGTCAGGATCTCTG
A. stuartgranti actr1 gene promoter       897 ACTCGTTGACAGCGTATAGACATGCTGATTGGCTCTCCGTGATGCCAGTCAGGATCTCTG
M. zebra actr1 gene promoter             1005 ACTCGTTGACAGCGTATAGACATGCTGATTGGCTCTCCGTGATGCCAGTCAGGATCTCTG
A. calliptera actr1 gene promoter        1004 ACTCGTTGACAGCGTATAGACATGCTGATTGGCTCTCCGTGATGCCAGTCAGGATCTCTG
A. macrocleithrum actr1 gene promoter       4 ACTCGTTGACAGCGTATAGACAAGCTGATTGGCTCTCCGTGATGCCAGTCAGGATCTCTG

                                              ....:.1090....:.1100....:.1110....:.1120....:.1130....:.1140
O. niloticus actr1 gene promoter          300 GAGCTTGATTGGCTGTCTGCACCGTGAGCCCGCCGCGATGCTGTGGTTGTTGTGGTCGGG
R. 'chilingali' actr1 gene promoter      1065 GAGCTTGATTGGCTGTCTGCACCGTGAGCCCGCCTCGGTGCTGTGGTTGTTGTGGACGGG
T. 'mauve' actr1 gene promoter           1066 GAGCTTGATTGGCTGTCTGCACCGTGAGCCCGCCTCGGTGCTGTGGTTGTTGTGGACGGG
A. stuartgranti actr1 gene promoter       957 GAGCTTGATTGGCTGTCTGCACCGTGAGCCCGCCTCGGTGCTGTGGTTGTTGTGGACGGG
M. zebra actr1 gene promoter             1065 GAGCTTGATTGGCTGTCTGCACCGTGAGCCCGCCTCGGTGCTGTGGTTGTTGTGGACGGG
A. calliptera actr1 gene promoter        1064 GAGCTTGATTGGCTGTCTGCACCGTGAGCCCGCCTCGGTGCTGTGGTTGTTGTGGACGGG
A. macrocleithrum actr1 gene promoter      64 GAGCTTGATTGGCTGTCTGCACCGTGAGCCCGCCTCGGTGCTGTGGTTGTTGTGGACGG-

                                              ....:.1150....:.1160....:.1170....:.1180....:.1190....:.1200
O. niloticus actr1 gene promoter          360 GCGGCAGCAGCAGCGGCAGAGGAGAGTCGACAGGCGTAGAGCGGAGCGGCTAACCGGCGC
R. 'chilingali' actr1 gene promoter      1125 GCG---GCAGCAGCGGCAGAGGAGCGTCGACAGGCGTAGAGCGGAGCTGCTAACCGGCGC
T. 'mauve' actr1 gene promoter           1126 GCG---GCAGCAGCGGCAGAGGAGCGTCGACAGGCGTAGAGCGGAGCTGCTAACCGGCGC
A. stuartgranti actr1 gene promoter      1017 GCG---GCAGCAGCGGCAGAGGAGCGTCGACAGGCGTAGAGCGGAGCTGCTAACCGGCGC
M. zebra actr1 gene promoter             1125 GCG---GCAGCAGCGGCAGAGGAGCGTCGACAGGCGTAGAGCGGAGCTGCTAACCGGCGC
A. calliptera actr1 gene promoter        1124 GCG---GCAGCAGCGGCAGAGGACCGTCGACAGGCGTAGAGCGGAGCTGCTAACCGGCGC
A. macrocleithrum actr1 gene promoter     123 ------------------------------------------------------------

                                              ....:.1210....:.1220....:.1230....:.1240....:.1250....:.1260
O. niloticus actr1 gene promoter          420 TGATTTTTACATGATTAAAACACTAAACGGGAACGAAGACTCACGCGGATCGATAGAAAC
R. 'chilingali' actr1 gene promoter      1182 TGATTTTTACATGATTAAAACACTAAACGGGCACGAAGACTCACGCGGATCGATAGAAAC
T. 'mauve' actr1 gene promoter           1183 TGATTTTTACATGATTAAAACACTAAACGGGCACGAAGACTCACGTGGATCGATAGAAAC
A. stuartgranti actr1 gene promoter      1074 TGATTTTTACATGATTAAAACACTAAACGGGCACGAAGACTCACGTGGATCGATAGAAAC
M. zebra actr1 gene promoter             1182 TGATTTTTACATGATTAAAACACTAAACGGGCACGAAGACTCACGCGGATCGATAGAAAC
A. calliptera actr1 gene promoter        1181 TGATTTTTACATGATTAAAACACTAAACGGGCACGAAGACTCACGCGGATCGATAGAAAC
A. macrocleithrum actr1 gene promoter     123 ------------------------------------------------------------

                                              ....:.1270....:.1280...
O. niloticus actr1 gene promoter          480 ACACCTGAAGCGCACGTGCGGTC > O. niloticus actr1 gene
R. 'chilingali' actr1 gene promoter      1242 ACACCTGAAGCGCACGTGCGGTC
T. 'mauve' actr1 gene promoter           1243 ACACCTGAAGCGCACGTGCGGTC
A. stuartgranti actr1 gene promoter      1134 ACACCTGAAGCGCACGTGCGGTC
M. zebra actr1 gene promoter             1242 ACACCTGAAGCGCACGTGCGGTC > M. zebra actr1 gene
A. calliptera actr1 gene promoter        1241 ACACCTGAAGCGCACGTGCGGTC
A. macrocleithrum actr1 gene promoter     123 -----------------------
```

**MAF_HUMAN.H11MO.1.B TFBS Motif**

**T Segregating TFBS variant**

**Fig. S39 – Alignment of actr1 gene promoter region in Lake Malawi cichlids, *O. niloticus*, and *A. calliptera*.** Predicted MAF motifs (blue shading) and polymorphic sites (orange text) in *O. niloticus* and *M. zebra actr1* gene promoter region. Orthologous alignment with other cichlid genomes and consensus *A. macrocleithrum* reads [2,4] shown with light grey or white shading indicating polymorphic sites.

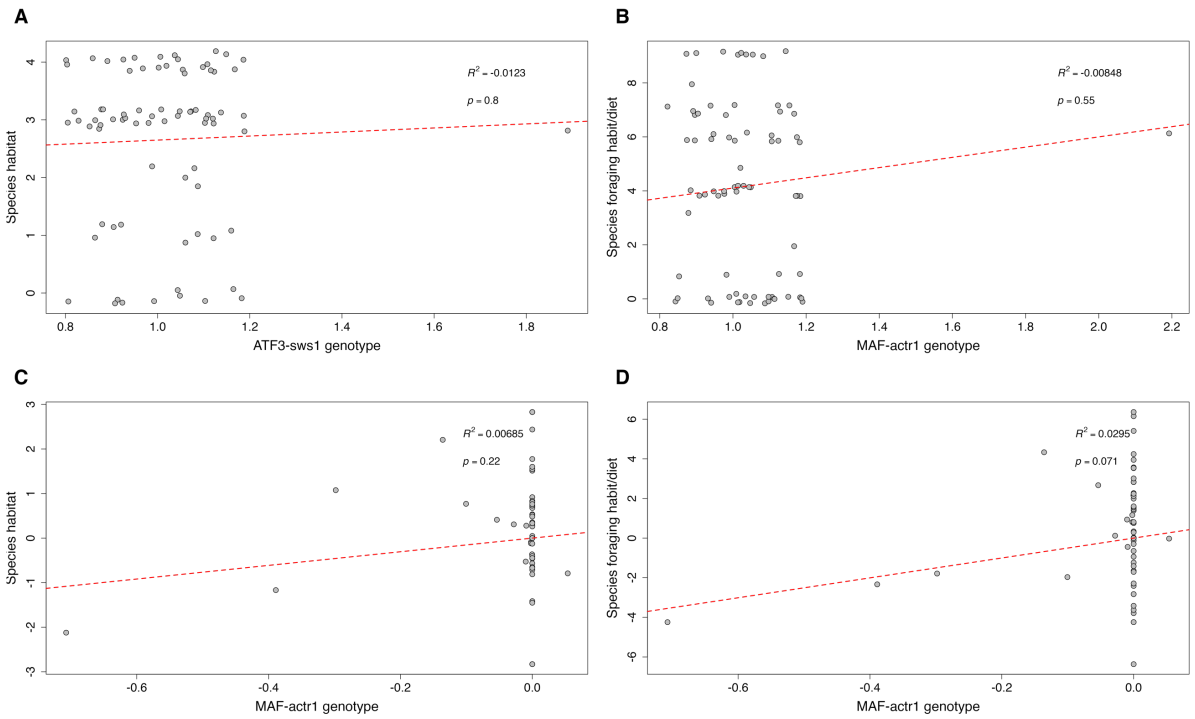


**Fig. S40 – Phylogenetic independent contrast analysis of MAF-actr1 target site genotypes of Lake Malawi species against their ecology.** Phylogenetic independent scatterplots of MAF-actr1 TFBS genotypes (1=T/T, 2=A/A) in 119 Lake Malawi individuals (73 species) against their respective (a) habitat (0=N/A, 1=Rock, 2=Pelagic, 3=Benthopelagic, 4=Demersal); and (b) foraging habit/diet (0=N/A, 1=Algae, 2=Aufwuchs, 3=Benthivore, 4=Fish, 5=Herbivore, 6=Invertebrates, 7=Mix, 8=Waste, 9=Zooplankton). Corresponding scatterplots of Lake Malawi ASTRAL phylogeny 3 and regression model fitted to MAF-actr1 TFBS genotypes (1=T/T, 2=A/A) of 119 Lake Malawi individuals (73 species) against their respective (c) habitat (0=N/A, 1=Rock, 2=Pelagic, 3=Benthopelagic, 4=Demersal); and (d) foraging habit/diet (0=N/A, 1=Algae, 2=Aufwuchs, 3=Benthivore, 4=Fish, 5=Herbivore, 6=Invertebrates, 7=Mix, 8=Waste, 9=Zooplankton). All data points used as per Supplementary Fig. S24, with overlapping coordinates 'jittered' around their respective point to highlight density. Adjusted r2 and p-value of each regression line shown in top right of each plot.

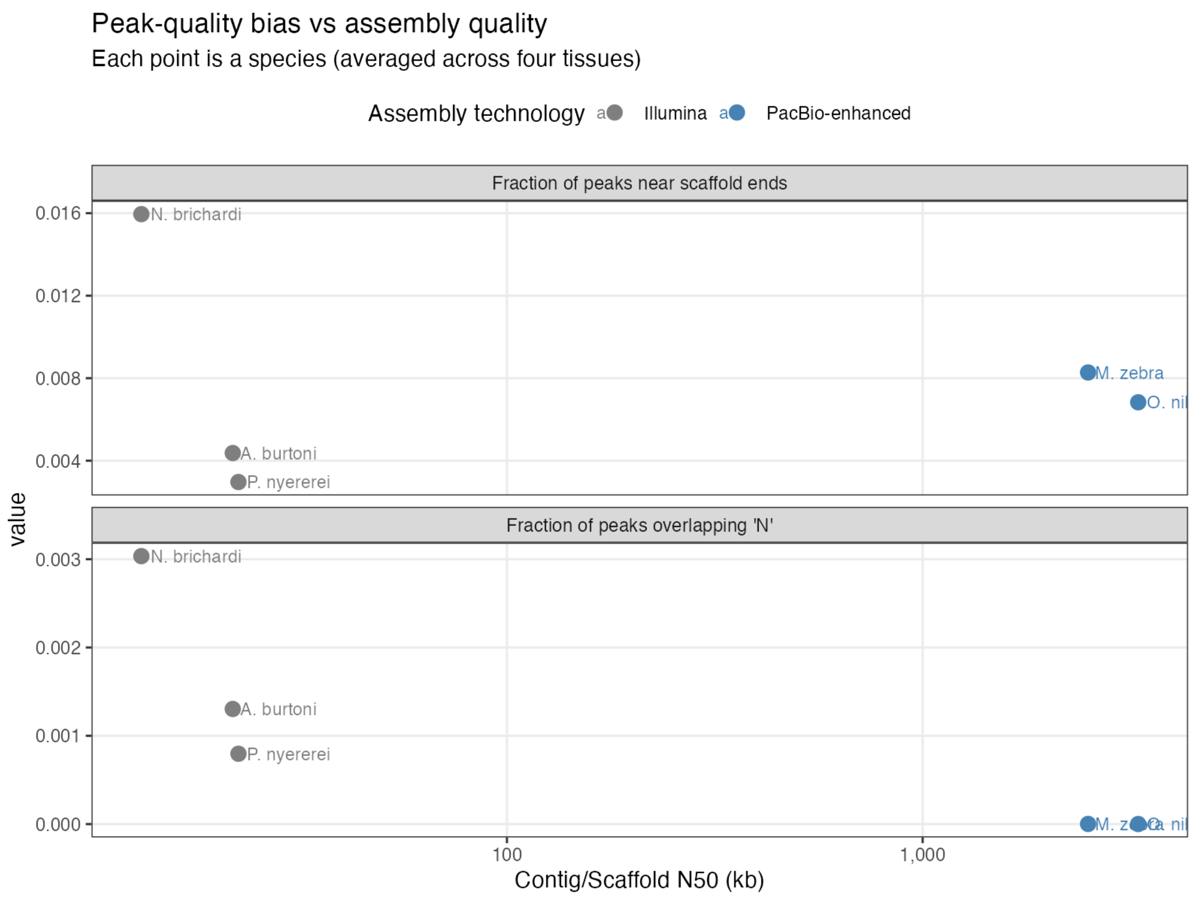


**Fig. S41 – Raw relationship between assembly contiguity and peak bias.** Species-level mean fractions of ATAC peaks overlapping at least 10% of gap-masked sequence (N/n) or lying within 100 bp of scaffold ends, plotted against assembly contiguity (N50, kb) for the five cichlid species. For each species, peak metrics were averaged across the four tissues examined. Points are coloured by assembly technology, with Illumina-only assemblies in grey and PacBio-enhanced assemblies in blue. The plot shows the unadjusted association between assembly quality and peak placement relative to gap-rich or terminal scaffold regions.

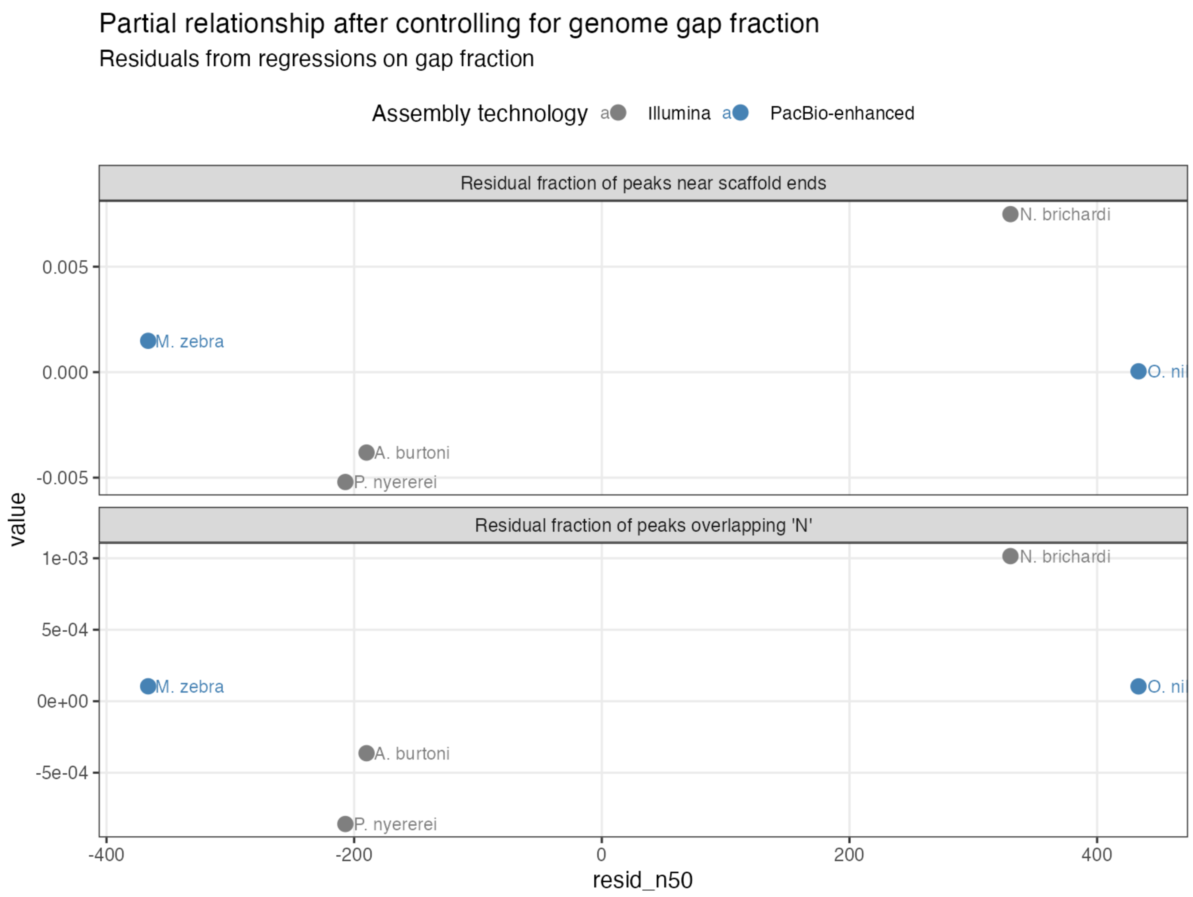


**Fig. S42 – Gap-adjusted relationship between assembly contiguity and peak bias.** Residual species-level mean fractions of ATAC peaks overlapping at least 10% of gap-masked sequence (N/n) or lying within 100 bp of scaffold ends, plotted against residual assembly contiguity after controlling for genome-wide gap fraction. Residuals were obtained from regressions on genome gap fraction, calculated as the proportion of bases in the genome FASTA encoded as N/n. Peak metrics were averaged across the four tissues for each species, and points are coloured by assembly technology as in Fig. S41. This adjusted analysis assesses whether the apparent association between contiguity and peak placement persists after accounting for species-wide assembly gap burden.
